# Synthetic protein condensates that recruit and release protein activity in living cells

**DOI:** 10.1101/2020.11.09.375766

**Authors:** Tatsuyuki Yoshii, Masaru Yoshikawa, Masahiro Ikuta, Shinya Tsukiji

**Author notes:** These authors contributed equally to this work (Tatsuyuki Yoshii, Masaru Yoshikawa, Masahiro Ikuta). Correspondence should be addressed to S.T.

## Abstract

Compartmentation of proteins into biomolecular condensates or membraneless organelles formed by phase separation is an emerging principle for the regulation of cellular processes. Creating synthetic condensates that accommodate specific intracellular proteins on demand would have various applications in chemical biology, cell engineering and synthetic biology. Here, we report the construction of synthetic protein condensates capable of recruiting and/or releasing proteins of interest in living mammalian cells in response to a small molecule or light. We first present chemogenetic protein-recruiting and -releasing condensates, which rapidly inhibited and activated signaling proteins, respectively. An optogenetic condensate system was successfully constructed that enables reversible release and sequestration of protein activity using light. This proof-of-principle work provides a new platform for chemogenetic and optogenetic control of protein activity in mammalian cells and represents a step towards tailor-made engineering of synthetic protein condensates with various functionalities.

## Introduction

Rapid control of protein activity in living cells with chemogenetics and optogenetics is a valuable tool for investigating the dynamic roles of proteins in biological systems and for biomedical applications such as cell-based therapies. Various approaches have been developed for chemogenetic and optogenetic control of proteins, including inducible oligomerization^1,2^, complementation of split proteins^3,4^ and engineered allosteric or conformational control^5,6^. However, most of these strategies require extensive trial and error to generate engineered proteins whose activity can be switched on or off effectively by small molecules or light. Moreover, engineering of such chemical- and light-switchable proteins is not always successful, depending on the tertiary structure and topology of the target protein. Controlling the subcellular localization of proteins is another powerful strategy for modulating protein activity with broad applicability. Several chemogenetic^7–13^ and optogenetic^8,14^ systems that enable rapid and site-specific protein translocation have been developed. However, localization-based protein control often suffers from elevated background activity in the basal state caused by overexpression of the target protein (in most cases, in a constitutively active form)^15^. The anchor-away^16^ and related techniques^15,17^ inhibit protein function by relocating the target protein to an organelle that is not the site of action. Although useful, concentrating overexpressed proteins to endogenous organelles for sequestering protein activity may lead to unintended negative effects at the targeted organelles. Clearly, the establishment of new methods for chemogenetic and optogenetic control of intracellular proteins is highly desired.

In this report, we set out to develop a conceptually new approach to control protein activity using synthetic protein-based designer organelles. In recent years, it has become increasingly evident that in addition to classical membrane-bound organelles, cells contain different types of organelles, called membraneless organelles or biomolecular condensates, that form by self-assembly and phase separation of proteins^18–21^. Cells regulate diverse biological and signaling processes by dynamically compartmentalizing proteins into such phase-separated condensates. In particular, an important role of condensates in cell physiology is that they can sequester proteins and effectively inhibit their activity. For example, in yeast, TORC1 (target of rapamycin complex 1) is sequestered in stress granules (known as protein and RNA condensates) upon heat stress, which results in the suppression of TORC1 signaling^22^. Inspired by this, we hypothesized that it should be possible to inhibit various proteins of interest by creating a synthetic condensate in cells as a designer membraneless organelle that recruits and sequesters proteins. Conversely, releasing a protein sequestered in a synthetic condensate into the cytoplasm could restore activity to the released protein. Such a mechanism, based on protein sequestration and release to and from synthetic condensates, is expected to be applicable for regulating various types of proteins without relying on endogenous organelles and may overcome many of the limitations of existing approaches. Furthermore, harnessing phase separation of proteins offers a strategy to construct synthetic, genetically encoded, membraneless organelles with desired functionalities in a modular manner^23–28^. We thus sought to create synthetic protein condensate-based designer organelles that are programmed to recruit and/or release proteins of interest in living cells in response to a small molecule or light. While recent efforts have been directed toward developing tools for artificially controlling the assembly and/or disassembly of protein condensates by small molecules^29,30^ or light^24,31–33^, few attempts have been made to engineer synthetic membraneless organelles with controllable protein recruitment and/or release functions. In pioneering work, Schuster and coworkers described enzyme-triggered cargo protein release from RGG-based designer condensates in living cells^23^. However, the system relied on the coexpression of a protease in cells, and rapid temporal control over cargo release was not achieved.

Here, we construct synthetic protein condensate systems, called SPREC (synthetic protein-recruiting/-releasing condensates), that are capable of controlling protein activity by externally regulated protein recruitment (sequestration) and/or release in living mammalian cells. As a scaffold to create SPREC, we used a previously reported fusion protein consisting of the p62 PB1 domain and the fluorescent protein Azami-Green (AG) (PB1-AG), which phase separates to form protein condensates in cells^25^. By rationally and modularly integrating the phase separating PB1-AG tandem protein polymer with a chemically inducible dimerization (CID) tool, we first created a chemogenetic SPREC-In system that rapidly recruits a protein of interest (POI) from the cytoplasm to the PB1-AG-based condensates by addition of a small-molecule dimerizer. Next, we coupled the SPREC-In system with an engineered proximity-dependent protease, leading to the development of a SPREC-Out system by which a POI previously expressed inside PB1-AG-based condensates (by fusion to the PB1-AG construct via a protease recognition sequence) is cleaved and released into the cytoplasm by small molecule-triggered protease recruitment. These chemogenetic SPREC-In and -Out systems achieved single-cycle protein recruitment and release switching, respectively. To realize reversible and repeatable protein recruitment/release control, we further engineered an optogenetic SPREC system by utilizing a light-responsive protein dimerization tool. Importantly, the chemogenetic and optogenetic SPREC systems are demonstrated to be applicable to control protein activity and cellular processes such as membrane ruffling and ERK signaling. The concept of SPREC opens the door to developing novel chemogenetic and optogenetic tools for the conditional control of various protein targets and may also offer a new direction in the design and engineering of protein condensate-based soft materials with tailor-made functions for biological and biomedical applications.

## Results

### Engineering of a chemogenetic SPREC-In system

Tandem fusion of the p62 PB1 domain (a homo-oligomeric protein) and AG (a tetramer protein) has been shown to undergo interconnected self-assembly and forms phase-separated condensates in cells upon expression^25^. We used the PB1-AG fusion protein polymer as a phase-separating scaffold for engineering SPREC systems. To induce protein recruitment into PB1-AG-based condensates with a small molecule, we utilized a CID technique, where the small-molecule rapamycin mediates the heterodimerization of FK506-binding protein (FKBP) and FKBP-rapamycin-binding domain (FRB)^34^. In our strategy, FRB is fused to the C-terminus of the PB1-AG fusion protein (PB1-AG-FRB) to construct an FRB-containing protein-assembled condensate (^FRB^PAC) in cells (**Fig. 1a**). Meanwhile, a target protein is expressed in the cytoplasm as a fusion with FKBP. We hypothesized that because ^FRB^PAC is a membraneless compartment, the FKBP-tagged protein is recruited and trapped in ^FRB^PAC by rapamycin-triggered FKBP-FRB dimerization. To test this, we cotransfected HeLa cells with PB1-AG-FRB and FKBP-tagged mCherry (FKBP-mCherry). Confocal fluorescence imaging showed micrometer-sized green fluorescent puncta in cells (**Fig. 1b**), indicating that PB1-AG-FRB formed phase-separated synthetic condensates incorporating the FRB domain, ^FRB^PAC. In contrast, FKBP-mCherry was distributed throughout the cytoplasm (**Fig. 1b**). Addition of rapamycin caused rapid recruitment of FKBP-mCherry to the ^FRB^PAC (**Fig. 1b**). The punctate fluorescent signals of the ^FRB^PAC were maintained and no noticeable disruption of the ^FRB^PAC was observed even after FKBP-mCherry recruitment. Time-lapse imaging revealed that FKBP-mCherry recruitment was completed within 1 min of adding rapamycin (200 nM) (**Fig. 1c** and **Supplementary Movie 1**). In contrast, no protein recruitment was observed in control experiments using a PB1-AG condensate lacking FRB, untagged mCherry (without FKBP), or DMSO (vehicle control) (**Fig. 1c** and **Supplementary Fig. 1**). These results demonstrated the validity of the SPREC-In system as a chemogenetic platform for controlled protein recruitment into synthetic condensates. This CID-based SPREC-In system was also operational in other cell lines, such as NIH3T3 and Cos-7 cells (**Supplementary Fig. 2**).

**Fig. 1.**
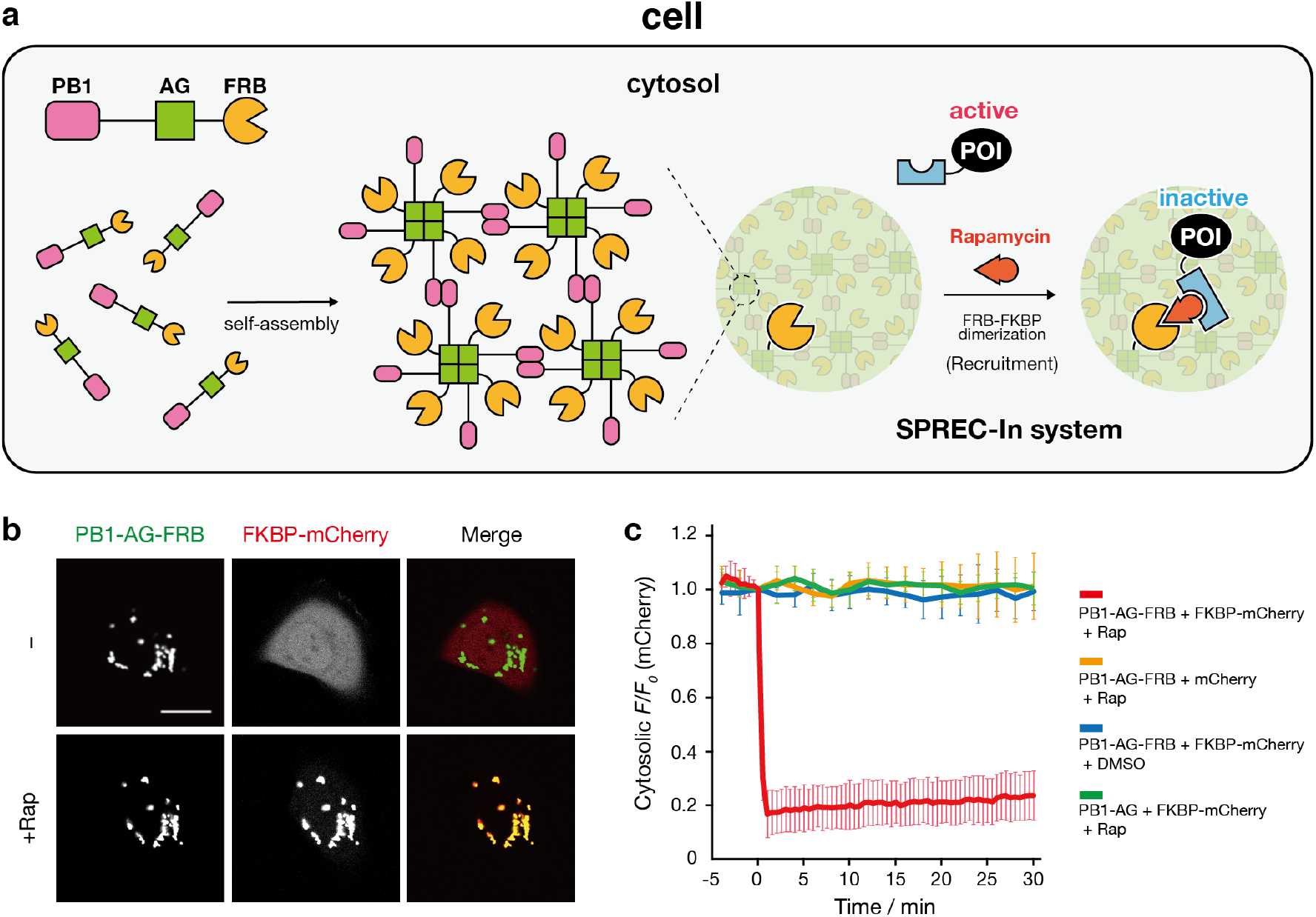
The concept and design of the SPREC-In system. **a** Schematic of the intracellular formation of ^FRB^PAC through self-assembly of PB1-AG-FRB. In the SPREC-In system, a FKBP-fused protein of interest (FKBP-POI) can be acutely recruited to ^FRB^PAC upon rapamycin-induced dimerization of FKBP and FRB. **b** Rapamycin-triggered FKBP-mCherry recruitment to ^FRB^PAC. Confocal fluorescence images of a HeLa cell coexpressing ^FRB^PAC (PB1-AG-FRB) and FKBP-mCherry before (upper panel) and 20 min after incubation with 200 nM rapamycin (lower panel). Scale bar, 20 μm. **c** Time course of FKBP-mCherry recruitment. HeLa cells coexpressing the indicated constructs were treated with 200 nM rapamycin or DMSO (fluorescence images of controls are shown in **Supplementary Fig. 1**). Normalized fluorescence intensities (*F*/*F*0) of FKBP-mCherry (or mCherry) in the cytoplasm were plotted against time. Data are presented as the mean ± SD (*n* > 10 cells).

### Conditional protein activity sequestration by the SPREC-IN system

The activity of a protein diffusing in the cytoplasm would be rendered inactive by sequestrating it into condensates. We investigated whether the SPREC-In system could be used to inactivate signaling proteins. We selected Vav2 as a model protein, which is a guanine nucleotide exchange factor (GEF) that activates Rho small GTPases^35,36^. Expression of a constitutively active catalytic domain of Vav2 (Vav2_cat_) induces plasma membrane protrusion and ruffling by activating the downstream small GTPase Rac^37,38^. We postulated that Vav2_cat_ would be functionally inactivated upon sequestration into synthetic condensates. To test this, we linked mCherry-tagged FKBP (FKBP-mCherry) to the catalytic (DH-PH-CRD) domain of Vav2 (FKBP-mCherry-Vav2_cat_) and coexpressed FKBP-mCherry-Vav2_cat_ in HeLa cells stably expressing the F-actin probe Lifeact-mTagBFP2 (Lifeact-BFP) with PB1-AG-FRB. Before rapamycin addition, PB1-AG-FRB formed ^FRB^PAC, while FKBP-mCherry-Vav2_cat_ distributed throughout the cytoplasm (**Fig. 2a**). Global lamellipodia formation and membrane ruffling was also observed, indicating that FKBP-mCherry-Vav2_cat_ was active. Upon rapamycin addition, rapid recruitment of FKBP-mCherry-Vav2_cat_ into ^FRB^PAC was observed, which was followed by retraction of lamellipodia (**Fig. 2a,b** and **Supplementary Movie 2**). Quantitative analysis of the cell area revealed a significant shrinkage of the cells by Vav2_cat_ sequestration into ^FRB^PAC (**Fig. 2c**). The use of a fluorescent biosensor for Rac^39^ showed that the activity of endogenous Rac was attenuated significantly upon Vav2_cat_ sequestration (**Supplementary Fig. 3**). No significant change in cell area was observed when DMSO was added, when mCherry-Vav2_cat_ lacking FKBP was used, or when FKBP-mCherry-Vav2_cat_ formed a heterodimer with a monomeric version of PB1-AG-FRB (mPB1-mAG-FRB) that does not form condensates (**Fig. 2c** and **Supplementary Fig. 4**). Therefore, Vav2_cat_ activity was suppressed rapidly by sequestering Vav2_cat_ in synthetic condensates using the SPREC-In system.

**Fig. 2.**
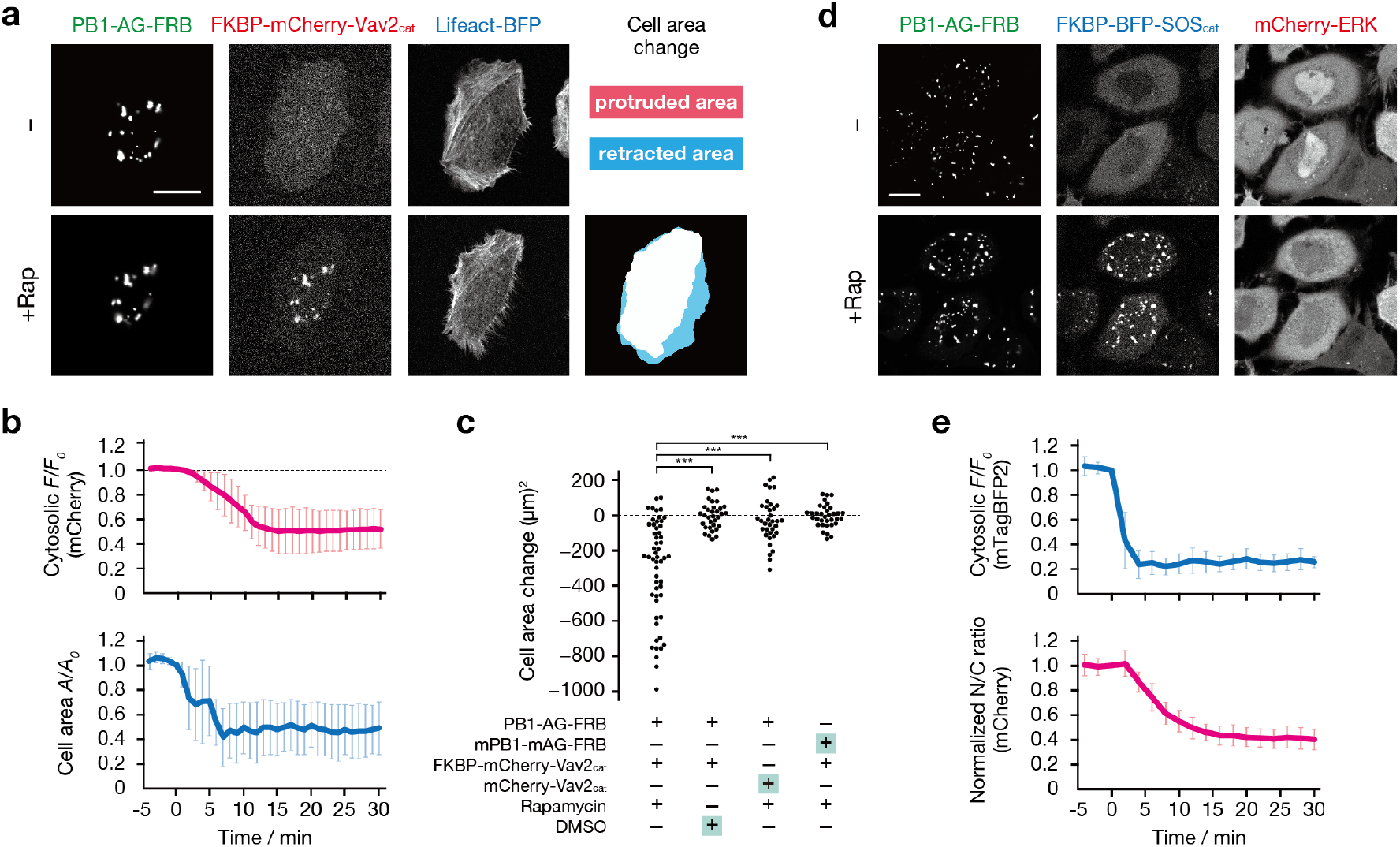
Chemogenetic protein activity sequestration using the SPREC-In system. **a** Vav2 activity sequestration. Confocal fluorescence images of a HeLa cell coexpressing ^FRB^PAC (PB1-AG-FRB), FKBP-mCherry-Vav2_cat_ and Lifeact-BFP before (upper panel) and 20 min after incubation with 200 nM rapamycin (lower panel). The protrusion-retraction map indicates morphological changes before and after rapamycin treatment. Scale bar, 20 μm. **b** Time course of Vav2_cat_ recruitment and cell area change. (Top) Normalized fluorescence intensities (*F*/*F*0) of FKBP-mCherry-Vav2_cat_ in the cytoplasm were plotted against time. (Bottom) Cell areas were calculated from Lifeact-BFP images, and the normalized values (*A*/*A*0) were plotted against time. **c** Quantification of cell retraction. The Vav2 sequestration assays were performed under the conditions shown in the graph. Cell area changes were calculated by determining the cell area difference before and after incubation with 200 nM rapamycin or DMSO for 20 min (*n* > 30 cells). The symbols indicate the results of *t* test analysis. \*\*\**p* < 0.001. **d** Sequestering SOS activity. Confocal fluorescence images of HeLa cells coexpressing ^FRB^PAC (PB1-AG-FRB), FKBP-BFP-SOS_cat_ and mCherry-ERK before (upper panel) and 30 min after incubation with 200 nM rapamycin (lower panel). Scale bar, 20 μm. **e** Time course of SOS_cat_ recruitment and ERK activity. (Top) SOS_cat_ recruitment was evaluated in the same manner as described for Vav2_cat_ (**b**). (Bottom) Normalized ratios of the nuclear to cytoplasmic fluorescence intensities (N/C ratios) of mCherry-ERK were plotted against time. Data are presented as the mean ± SD (*n* > 10 cells).

To further test the utility of the SPREC-In system, we next attempted to control the activity of SOS (son of sevenless), which is a Ras GEF that functions as an activator of the Ras/ERK pathway^40,41^. The catalytic (DH-PH) domain of SOS (SOScat)^42^ was attached to the C-terminus of a FKBP-mTagBFP2 fusion protein (FKBP-BFP-SOS_cat_). mCherry-tagged ERK (mCherry-ERK) was used to monitor SOS activity^12^. As ERK translocates from the cytoplasm to the nucleus upon SOS-mediated activation of the Ras/ERK pathway, the SOS activity can be quantified by the ratio of nuclear to cytoplasmic fluorescence intensities (N/C ratio) of the mCherry-ERK reporter. As expected, coexpression of PB1-AG-FRB, FKBP-BFP-SOS_cat_ and mCherry-ERK in HeLa cells caused formation of ^FRB^PAC and the cytoplasmic localization of FKBP-BFP-SOS_cat_ (**Fig. 2d**). mCherry-ERK was localized predominantly in the nucleus, indicating that the Ras/ERK pathway is active (**Fig. 2d**). By adding rapamycin, FKBP-BFP-SOS_cat_ was efficiently recruited to ^FRB^PAC within 5 min, which was followed by the export of mCherry-ERK to the cytoplasm, indicating ERK inactivation (**Fig. 2d,e** and **Supplementary Movie 3**). The N/C ratio decrease reached a plateau approximately 20 min after rapamycin addition (**Fig. 2e**). Such rapamycin-triggered ERK inactivation was not observed when BFP-SOS_cat_ lacking FKBP was used (**Supplementary Fig. 5**). Taken together, these results demonstrated that the SPREC-In system conditionally inactivated signaling proteins and related cellular processes by sequestering protein activity.

### Development of a chemogenetic SPREC-Out system

The utility of the SPREC platform would be expanded greatly if this platform could be used as a tool to release protein activity in living cells. Thus, we sought to couple the CID-based SPREC-In system with proximity-induced proteolysis using the tobacco etch virus (TEV) protease (TEVp)^43^. TEVp has high sequence-specificity and does not cleave endogenous proteins in mammalian cells^43,44^. In our strategy, a target protein is fused to the C-terminus of PB1-AG-FRB via a cleavage sequence of TEVp (TEVcs or TCS) (**Fig. 3a**). The resulting target fusion protein, PB1-AG-FRB-TEVcs-POI, is expressed in cells together with PB1-AG and FKBP-tagged TEVp (FKBP-TEVp). In the basal state, the target protein is expressed and confined in synthetic PB1-AG-based condensates (^FRB-TCS-POI^PAC) formed by coassembly of PB1-AG-FRB-TEVcs-POI and PB1-AG, while FKBP-TEVp is expressed in the cytoplasm. Upon exposure to rapamycin, FKBP-TEVp is recruited to the ^FRB-TCS-POI^PAC by dimerization with FRB, which leads to proximity-induced cleavage of TEVcs and release of the target protein from the condensates. For this approach to work as a useful protein activity release tool, it is imperative to minimize background cleavage of TEVcs in the absence of rapamycin. Based on recent work by Wang and coworkers, we chose the C-terminally truncated TEVp (Δ220–242) and its low affinity TEVcs sequence (ENLYFQ/L) because this enzyme-substrate combination exhibits low background cleavage and high proximity-induced cleavage efficiency^43^.

**Fig. 3.**
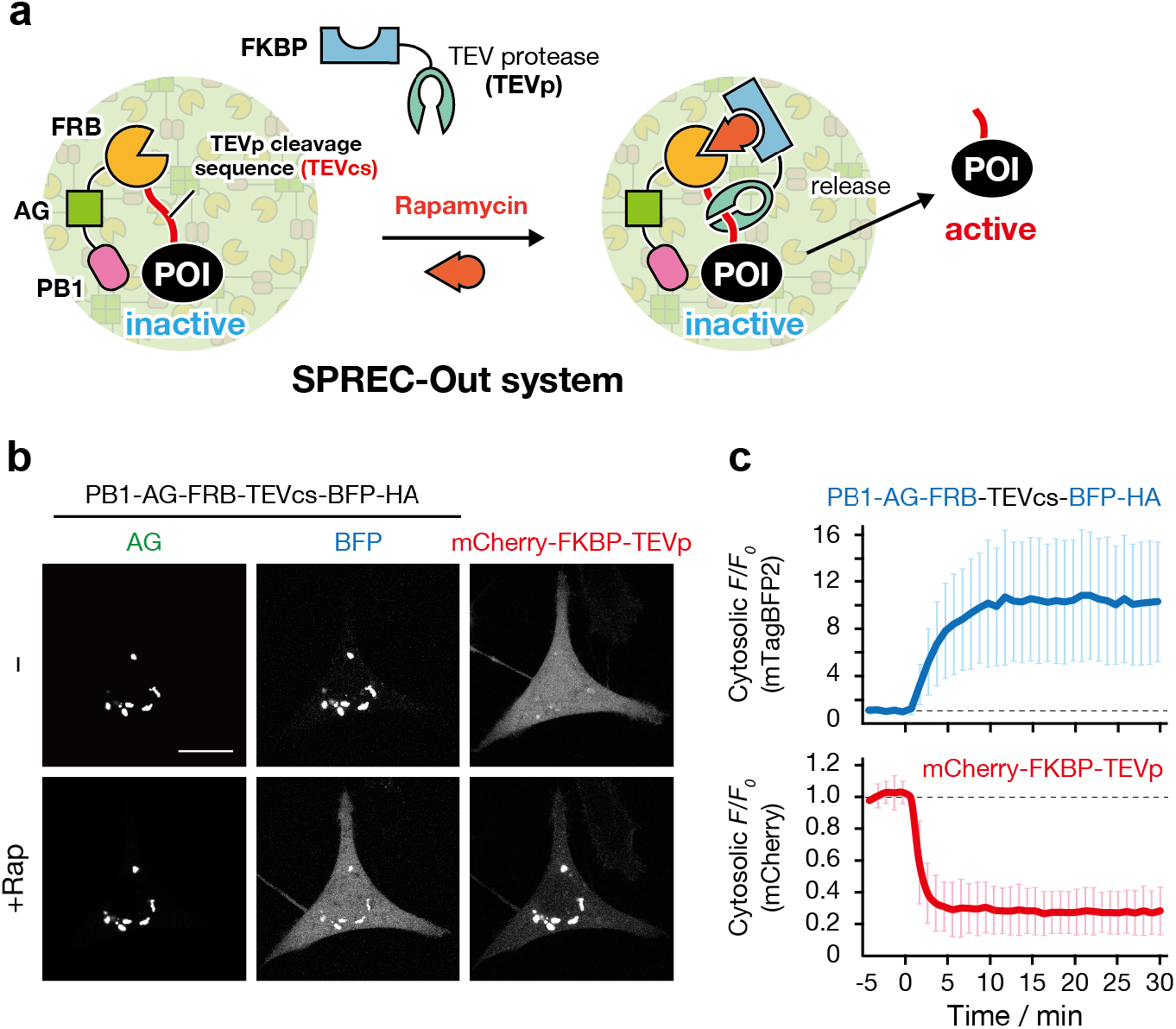
Construction of the SPREC-Out system. **a** Schematic of the concept and design. In this system, a protein of interest (POI) is initially expressed and confined in synthetic PB1-AG-based condensates by fusion to PB1-AG-FRB via a TEV protease (TEVp) cleavage sequence (TEVcs). Upon rapamycin addition, FKBP-fused TEVp is recruited to the condensate and cleaves the TEVcs, releasing the POI into the cytoplasm. **b** Rapamycin-triggered BFP-HA release. Confocal fluorescence images of a HeLa cell coexpressing ^FRB-TCS-BFP-HA^PAC (PB1-AG and PB1-AG-FRB-TEVcs-BFP-HA) and mCherry-FKBP-TEVp before (upper panel) and 30 min after incubation with 200 nM rapamycin (lower panel). Scale bar, 20 μm. **c** Time course of TEVp recruitment and BFP release. Normalized fluorescence intensities (*F*/*F*_0_) of mCherry-FKBP-TEVp (bottom) and BFP-HA (top) in the cytoplasm were plotted against time. Data are presented as the mean ± SD (*n* > 10 cells).

To test the aforementioned strategy, we selected HA-tagged mTagBFP2 (BFP-HA) as a model protein for release and generated PB1-AG-FRB-TEVcs-BFP-HA. BFP fluorescence was mostly found as fluorescent puncta when the protein was coexpressed in cells with PB1-AG and mCherry-FKBP-TEVp, and this fluorescence merged well with AG fluorescence (**Fig. 3b**). Therefore, the target protein BFP-HA was confined efficiently in condensates (^FRB-TCS-BFP-HA^PAC) with almost no background cleavage by the TEVp construct that was expressed in the cytoplasm. However, addition of rapamycin rapidly recruited mCherry-FKBP-TEVp to the ^FRB-TCS-BFP-HA^PAC and caused a marked gradual increase in BFP fluorescence in the cytoplasm, which reached a plateau after approximately 10 min (**Fig. 3b,c** and **Supplementary Movie 4**). Several control experiments were performed to establish that the BFP fluorescence increase in the cytoplasm was due to the cleavage of TEVcs by TEVp and not by the disassembly of the condensates. Recruitment of mCherry-FKBP lacking the TEVp domain into ^FRB-TCS-BFP-HA^PAC had no effect on BFP fluorescence (**Fig. 3c** and **Supplementary Fig. 6**). Additionally, no increase in cytosolic BFP fluorescence was observed upon rapamycin-induced recruitment of TEVp into the condensates when BFP-HA was fused to PB1-AG-FRB via an uncleavable TEVcs (TEVuncs) (**Fig. 3c** and **Supplementary Fig. 6**). Furthermore, we verified the rapamycin- and TEVp-dependent cleavage of PB1-AG-FRB-TEVcs-BFP-HA by western blotting (**Supplementary Fig. 7**). Overall, we have established the SPREC-Out system in which a target protein sequestered in PB1-AG-based condensates can be rapidly released to the cytoplasm in living cells on a time scale of minutes by rapamycin-triggered TEVp recruitment.

### Conditional protein activity release by the SPREC-Out system

We next investigated whether the SPREC-Out system is applicable as a tool to release protein activity in living cells. We chose Vav2_cat_ (BFP-HA-Vav2_cat_) as a model protein again, but here for controlled activation. According to the design strategy established above, we first generated PB1-AG-FRB-TEVcs-BFP-HA-Vav2_cat_ and coexpressed this construct with PB1-AG and mCherry-FKBP-TEVp in cells stably expressing Lifeact-tdiRFP670 (Lifeact-iRFP). Fluorescence of BFP and AG were observed as colocalized puncta, indicating the formation of condensates incorporating the Vav2_cat_ construct (^FRB-TCS-BFP-HA-Vav2^PAC) (**Fig. 4a**). Despite the expression of the constitutively active Vav2_cat_, cells showed only a basal level of membrane ruffling because of the efficient sequestration of Vav2_cat_ activity in condensates. Upon rapamycin-induced recruitment of mCherry-FKBP-TEVp to the ^FRB-TCS-BFP-HA-Vav2^PAC, BFP-HA-Vav2_cat_ was released into the cytoplasm, which caused a pronounced induction of membrane protrusion and ruffling (**Fig. 4a,b** and **Supplementary Movie 5**). The use of a fluorescent biosensor for Rac showed the activation of endogenous Rac upon Vav2_cat_ release (**Supplementary Fig. 8**). The TEVp-mediated cleavage of PB1-AG-FRB-TEVcs-BFP-HA-Vav2_cat_, generating BFP-HA-Vav2_cat_, was also confirmed by western blotting (**Supplementary Fig. 9**). No significant change in cell morphology was observed when recruiting mCherry-FKBP to ^FRB-TCS-BFP-HA-Vav2^PAC, or when Vav2_cat_ was fused to PB1-AG-FRB via TEVuncs (**Fig. 4b** and **Supplementary Fig. 10**).

**Fig. 4.**
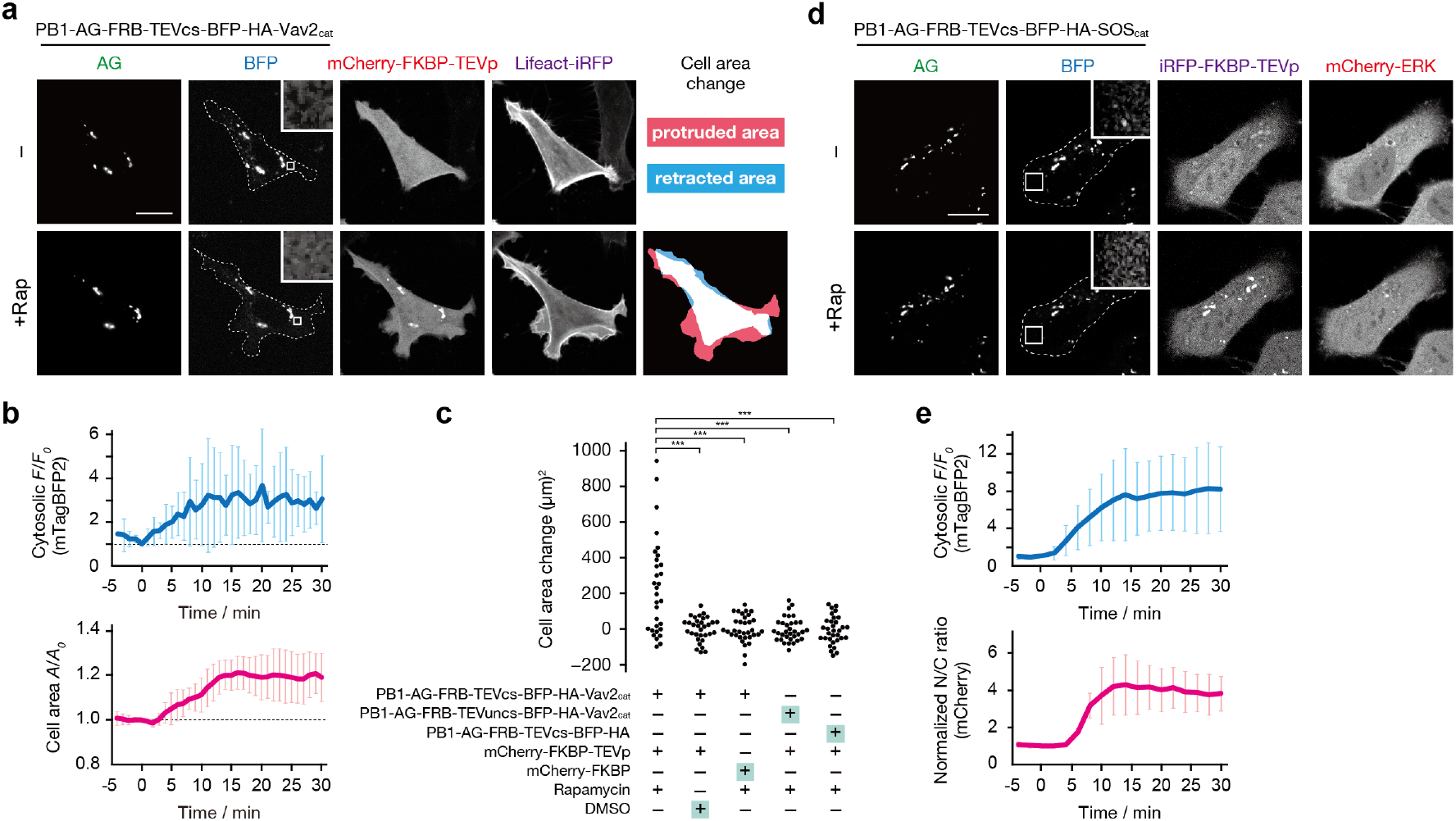
Chemogenetic protein activity release by the SPREC-Out system. **a** Vav2 activity release. Confocal fluorescence images of a HeLa cell coexpressing ^FRB-TCS-BFP-HA-Vav2^PAC (PB1-AG and PB1-AG-FRB-TEVcs-BFP-HA-Vav2_cat_), mCherry-FKBP-TEVp and Lifeact-iRFP before (upper panel) and 20 min after incubation with 200 nM rapamycin (lower panel). The dashed line indicates the cell boundary. The inserted window shows the high magnification and enhanced brightness image of the cytoplasmic region indicated by the box. Scale bar, 20 μm. **b** Time course of Vav2_cat_ release and cell area change. (Top) Normalized fluorescence intensities (*F*/*F*0) of BFP-HA-Vav2_cat_ in the cytoplasm were plotted against time. (Bottom) Cell areas were calculated from Lifeact-iRFP images, and the normalized values (*A*/*A*0) were plotted against time. **c**Quantification of cell protrusions. The Vav2 release assays were performed under the conditions shown in the graph. Cell area changes were calculated by determining the cell area difference before and after incubation with 200 nM rapamycin or DMSO for 20 min (*n* > 30 cells). The symbols indicate the results of *t* test analysis. \*\*\**p* < 0.001. (**d**) Releasing SOS activity. Confocal fluorescence images of HeLa cells coexpressing ^FRB-TCS-BFP-HA-SOS^PAC (PB1-AG and PB1-AG-FRB-TEVcs-BFP-HA-SOScat) and mCherry-ERK before (upper panel) and 30 min after incubation with 200 nM rapamycin (lower panel). Scale bar, 20 μm. (**e**) Time course of SOS_cat_ release and ERK activity. (Top) SOS_cat_ release was evaluated in the same manner as described for Vav2_cat_ (**b**). (Bottom) Normalized ratios of the nuclear to cytoplasmic fluorescence intensities (N/C ratios) of mCherry-ERK were plotted against time. Data are presented as the mean ± SD (*n* > 10 cells).

The SPREC-Out system could also induce activation of the Ras/ERK pathway. We cotransfected PB1-AG-FRB-TEVcs-BFP-HA-SOS_cat_ with PB1-AG and tdiRFP670-FKBP-TEVp in cells expressing mCherry-ERK. In the absence of rapamycin, the SOS_cat_ construct was trapped in condensates, and mCherry-ERK was observed predominantly in the cytoplasm in an inactive state. Upon rapamycin addition, BFP-HA-SOS_cat_ was released into the cytoplasm, which triggered activation and translocation of mCherry-ERK into the nucleus (**Fig. 4c,d** and **Supplementary Fig. 11** and **Supplementary Movie 6**). No ERK activation occurred in the presence of PD184352 (50 μM), a MEK inhibitor, or in control experiments that do not release SOS activity (**Supplementary Fig. 12**). These results clearly demonstrated that the SPREC-Out system rapidly turns on cellular processes such as membrane protrusion/ruffling and Ras/ERK signaling by chemogenetic release of protein activity.

### An optogenetic SPREC system for reversible protein activity control

The chemogenetic SPREC systems described above allow only unidirectional protein recruitment or release because of the irreversible nature of the rapamycin CID tool^45^. Controlled reversible sequestration and release of protein activity in cells would have various applications for cell engineering and synthetic biology. Therefore, we sought to further generate a SPREC system with reversibility. For this purpose, we focused our attention on the LOVTRAP system, which is an optogenetic tool for reversible light-induced protein dissociation using a pair of photoreceptor LOV2 and a small protein named Zdk1^46^. Zdk1 associates with LOV2 in the dark but dissociates upon blue light illumination. To integrate the LOVTRAP system into SPREC, we replaced FRB in the PB1-AG-FRB construct with the LOV2 domain (**Fig. 5a**). To avoid undesired stimulation of LOV2 during fluorescence imaging of condensates, we also replaced AG with a tetrameric red fluorescent protein Monti-Red (MR) (PB1-MR-LOV2). We then cotransfected HeLa cells with PB1-MR-LOV2 and miRFP670-tagged Zdk1 (miRFP-Zdk1). In the dark state, the formation of LOV2-containing protein-assembled condensates (^LOV2^PAC) was observed clearly as red fluorescent puncta (**Fig. 5b**). Notably, miRFP fluorescence colocalized well with the red fluorescence from the ^LOV2^PAC and was barely visible in the cytoplasm, indicating that miRFP-Zdk1 was incorporated efficiently into ^LOV2^PAC in the dark state. However, upon blue light illumination, cytosolic miRFP fluorescence increased rapidly as miRFP-Zdk1 was released from LOV2PMC into the cytoplasm (**Fig. 5b,c**). The diffusing miRFP-Zdk1 was recruited back to the condensate when the blue light illumination was switched off (**Fig. 5b,c**). The light-controlled protein release-and-recruitment cycle could be repeated at least several times (**Supplementary Movie 7**). The duration the protein was present in the cytoplasm could be adjusted by varying the time the blue light illumination was switched on (**Supplementary Fig. 13**). Such light-dependent cytoplasmic release of miRFP-Zdk1 never occurred in a control experiment using PB1-MR condensates containing a LOV2 dark mutant^46^ (PB1-MR-LOV2dark) (**Supplementary Fig. 14**). Furthermore, when we applied this optogenetic SPREC (optoSPREC) system to Vav2, we were able to stimulate cell protrusion and retraction cycles in a reversible and repeatable manner using blue light (**Fig. 5e,f**, **Supplementary Movie 8**, and **Supplementary Fig. 15**). Therefore, the LOVTRAP-based optoSPREC system is a new tool to manipulate cellular processes based on light-controlled protein activity release from synthetic condensates.

**Fig. 5.**
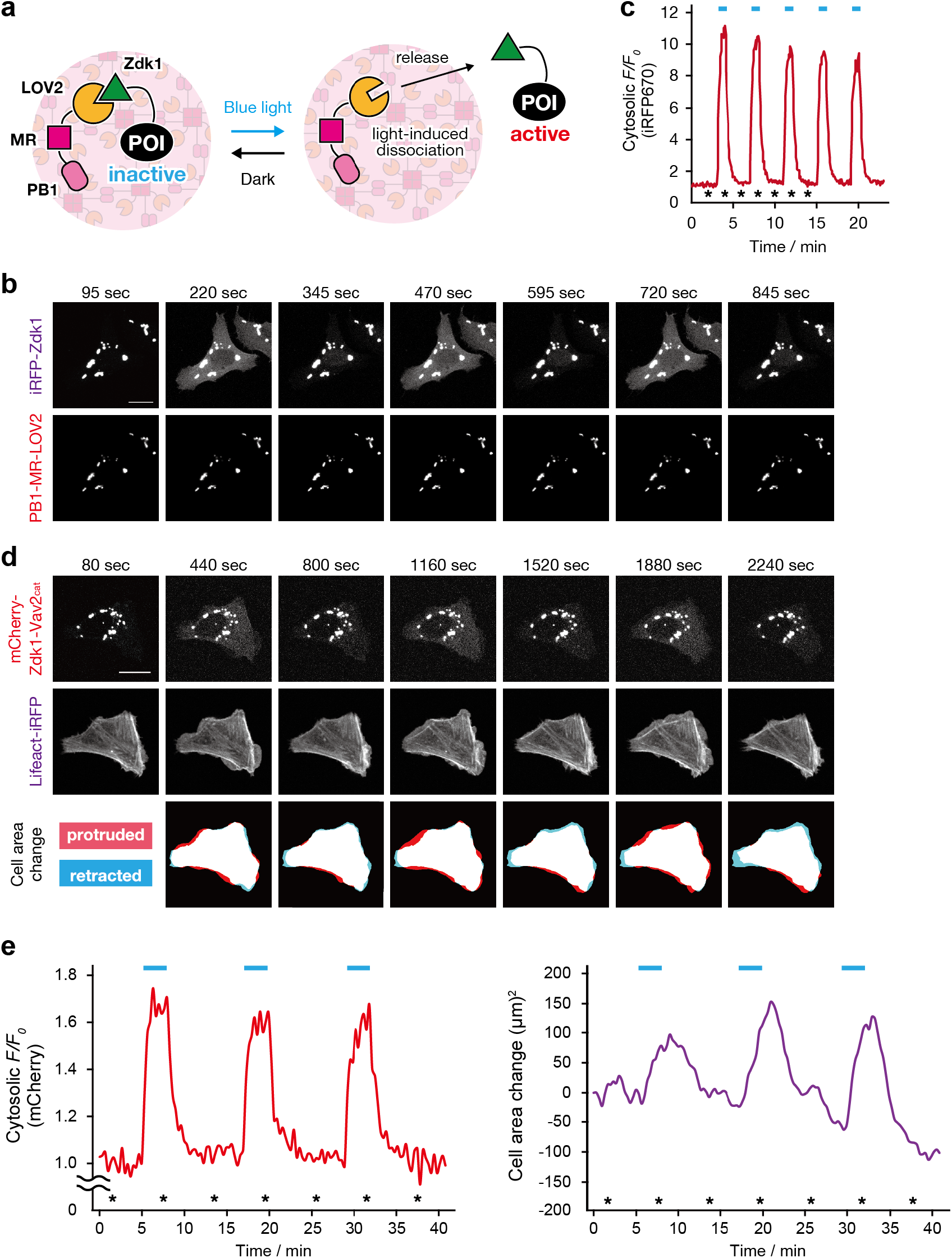
Engineering of the LOVTRAP-based optoSPREC system. **a** Schematic of the concept and design. In this system, a Zdk1-fused protein of interest (Zdk1-POI) is sequestered in synthetic ^LOV2^PAC (formed by assembly of PB1-MR-LOV2) by binding to the LOV2 domain in the dark but is released to the cytoplasm upon illumination with blue light. The Zdk1-POI is returned to the ^LOV2^PAC by switching off the light, enabling optical reversible control of protein activity. **b** Time-lapse confocal fluorescence images of HeLa cells coexpressing ^LOV2^PAC (PB1-MR-LOV2) and miRFP-Zdk1 under alternating periods of darkness and blue light illumination (440 nm). Images were taken at time points indicated by asterisks shown in panel **c**. Scale bar, 20 μm. **c** Time course of light-controlled miRFP-Zdk1 release from the ^LOV^PAC to the cytoplasm. The blue bars indicate periods of blue light illumination. Normalized fluorescence intensities (*F*/*F*0) of miRFP-Zdk1 in the cytoplasm were plotted against time. **d** Light-mediated reversible control of Vav2 activity. Time-lapse confocal fluorescent images of a HeLa cell coexpressing non-fluorescent ^LOV2^PAC [PB1-AG-LOV2 containing a Y63L mutation in AG], mCherry-Zdk1-Vav2_cat_ and Lifeact-iRFP under alternating periods of darkness and blue light illumination. Images were taken at time points indicated by asterisks shown in panel **e**. Scale bar, 20 μm. **e** Time course of Vav2_cat_ release and cell area change. (Left) Normalized fluorescence intensities (*F*/*F*0) of mCherry-Zdk1-Vav2_cat_ in the cytoplasm were plotted against time. (Right) Cell areas were calculated from Lifeact-iRFP images, and cell area changes were plotted against time. The repeated Vav2 activity release was observed at least three times.

## Discussion

Biological phase separation is an emerging concept in cell biology. A growing number of studies have shown that compartmentation of proteins and other biomolecules such as RNA into biomolecular condensates (or membraneless organelles) formed by phase separation play critical roles in the regulation of diverse cellular processes ranging from RNA processing and translation regulation to cell signaling^18–21^. This has spurred development of a variety of synthetic biology tools for investigating phase separation in living cells. Currently, most efforts have focused on designing protein- or protein-RNA-based condensates comprising intrinsically disordered proteins with an aim to understand the fundamental principles underlying the formation, physical properties and biological function of membraneless organelles found in nature^47–50^. There is also significant interest in engineering molecular tools that allow researchers to control the assembly and disassembly of condensates in living cells using external inputs such as light^24,31–33^ or small molecules^29,30^. However, attempts to create and harness synthetic condensates as a platform for manipulating intracellular protein function has so far been limited.

As a pioneering work, Lee and coworkers developed a light-inducible protein clustering system, termed LARIAT (light-activated reversible inhibition by assembly trap), which sequesters target proteins into condensates (clusters) formed by multimeric proteins and a blue light-mediated heterodimerization module^51^. The LARIAT system has been demonstrated to inactivate various signaling proteins rapidly^51,52^ or even mRNA translation^53^. However, the LARIAT technique requires continuous illumination to maintain clustering (i.e., inhibition) of target proteins and is not suitable for controlled protein activation. Recently, Schuster et al. reported synthetic membraneless organelles capable of incorporating cargo proteins^24^. Controlled release of a cargo protein from synthetic organelles in living cells was also achieved by expressing a protease. However, the system lacks temporal control over cargo recruitment and release, and its applicability as a tool for regulating protein function has not been demonstrated.

In this work, we designed and engineered three types of synthetic protein condensates, SPREC-In, SPREC-Out and optoSPREC, that are capable of recruiting and/or releasing specific target proteins in living mammalian cells in response to a small molecule or light. In the chemogenetic SPREC-In system, FKBP-tagged proteins are rapidly recruited from the cytoplasm to FRB-containing condensates (^FRB^PAC) within minutes following addition of the small-molecule rapamycin, resulting in sequestration of protein activity. The SPREC-Out system, which combines the SPREC-In scheme with a proximity-dependent protease, triggers protein release from condensates to the cytoplasm within approximately 10–20 minutes following addition of rapamycin. The chemogenetic SPREC-In and -Out systems achieve only single-cycle protein recruitment and release induction, respectively, and lack reversibility because of the irreversible nature of rapamycin-mediated heterodimerization of FKBP and FRB^45^. In contrast, the LOVTRAP-based optoSPREC system allows reversible and repeatable protein release and recruitment on the timescale of seconds by switching blue light on and off. We further demonstrated the applicability of these SPREC tools for temporally controlling protein activity and cellular processes such as membrane ruffling and ERK signaling. Notably, the SPREC systems are easy to use because they only require the genetic fusion of a target protein to the module component including FKBP (for SPREC-In), PB1-AG-FRB-TCV (for SPREC-Out), or Zdk1 (for optoSPREC). The FKBP (12 kDa) and Zdk1 (6.5 kDa) protein tags used in the SPREC-In and optoSPREC systems, respectively, are small, which minimizes potential perturbation to the function of target proteins in the cytoplasmically diffused (i.e., active) state. The SPREC-Out system releases the target protein in an almost traceless manner; only the Leu residue is in principle added to the N-terminus of the target protein after TEVcs cleavage. Thus, we believe that the present SPREC platforms will offer powerful new tools and strategies for chemogenetic and optogenetic control of signaling proteins in mammalian cells.

Another important feature of our SPREC approach is that phase-separated protein condensates are generated in cells by self-assembly of genetically encoded protein polymers consisting of multiple modular domains. This modular design allows engineering of tailor-made protein condensates with diverse chemical, physical and biological properties^23–28^. We envision that the construction of SPREC using various protein modules, such as ligand-binding domains, oligomerizing domains, nanobodies, enzymes and photoreceptors (optogenetic modules), will provide a variety of new functional synthetic condensates or soft materials that can incorporate and manipulate proteins (and even other biomolecules such as RNA) in living cells in unprecedented ways. Such efforts not only expand the potential of the SPREC concept, but also may contribute to the fundamental understanding of biomolecular phase separation in biology.

## Methods

### Plasmid construction

All constructs and their cDNA and amino acid sequences are listed in **Supplementary Fig. 16** and **17**. The following plasmids were purchased or kindly provided from the following companies or researchers: phAG-MCL (encoding Azami-Green), pMonti-Red-MCL and pER-mAG1 from Medical and Biological Laboratories (MBL); pC_4_-RHE [encoding FRB(T2098L)] and pC4EN-F1 (encoding FKBP12) from ARIAD Pharmaceuticals; pEGFP-Vav2 from W. D. Heo (KAIST)^51^; pmTagBFP2-Lifeact-7 (Addgene plasmid #54602) from M. W. Davidson (Florida State University)^54^; pAL197 (encoding PakGBD) (Addgene plasmid #22280) from C. A. Voigt (MIT)^39^; pCAG-H2B-tdiRFP-IP (encoding tdiRFP670) (Addgene plasmid #47884) from Y. Miyanari (Kanazawa University)^55^; pHA-Sos1 (Addgene plasmid #32920) from D. Bar-Sagi (NYU Langone Health)^56^; pcDNA3.1 TEV (full length) (Addgene plasmid #64276) from X. Shu (University of California, San Francisco)^57^; pTriEx-NTOM2-LOV2 (Addgene plasmid #81009) and pTriEx-mCherry-Zdk1-VAV2 DH/PH/C1 (Addgene plasmid #81060) from K. Hahn (University of North Carolina at Chapel Hill)^46^; and pmiRFP670-N1 (Addgene plasmid #79987) from V. Verkhusha (Albert Einstein College of Medicine)^58^. We used pAsh-MCL (encoding the PB1 domain) (MBL), pmCherry-C1 (Clontech) and pCAGGS [kindly provided by J. Miyazaki (Osaka University)]^59^ as vectors for constructing expression plasmids. We also used pPBbsr (blasticidin S resistance)^60^ as a *piggyBac* donor vector for establishing stable cell lines. Monomeric PB1 (mPB1) was generated by introducing D69A and D71R mutations in PB1^61^. Non-fluorescent colorless Azami-Green was generated by mutating the chromophore-forming Tyr 63 with leucine (Y63L)^62^. Light-insensitive LOV2, LOV2(dark), was generated by introducing C450A, L514K, G528A, L531E and N538E mutations in LOV2^46^. All expression plasmids were generated using standard cloning techniques or Gibson assembly (NEB). All PCR-amplified sequences were verified by DNA sequencing.

### Cell culture and transfection

HeLa, NIH3T3 and Cos-7 cells were cultured in Dulbecco’s modified Eagle’s medium (DMEM) (Wako) supplemented with 10% heat-inactivated fetal bovine serum (Biowest), penicillin (100 U/mL) and streptomycin (100 μg/mL) at 37 °C under a humidified 5% CO_2_ atmosphere. For transient expression experiments, cells were transfected using Polyethylenimine MAX (Polysciences Inc.) or 293fectin (Invitrogen) according to the manufacturer’s instructions.

### Establishment of stable cell lines

A *piggyBac* transposon system^63^ was employed to establish HeLa cells stably expressing the indicated constructs. HeLa cells were cotransfected with a *piggyBac* donor vector (pPBbsr) encoding a desired protein and pCMV-mPBase encoding the *piggyBac* transposase (provided by Dr. Allan Bradley, Wellcome Trust Sanger Institute)^64^ using Polyethylenimine MAX. Cells were selected with 2 μg/mL puromycin or 10 μg/mL blasticidin S for at least 10 days. Bulk populations of selected cells were used.

### Live-cell imaging

Confocal fluorescence imaging was performed on an IX83/FV3000 confocal laser-scanning microscope (Olympus) equipped with a PlanApo N 60×/1.42 NA oil objective (Olympus), a Z drift compensator system (IX3-ZDC2, Olympus) and stage top incubator (Tokai Hit). Lasers used for excitation were: 405 nm for mTagBFP2; 488 nm for Azami-Green and EGFP; 561 nm for mCherry and Monti-Red; and 640 nm for tdiRFP670 and miRFP670. Live-cell imaging was performed at 37 °C under a humidified 5% CO_2_ atmosphere. Fluorescence and differential interference contrast (DIC) images were analyzed using the Fiji distribution of ImageJ^65^.

### SPREC-In assay

To conduct the rapamycin-induced protein recruitment assay, 1 × 10^5^ HeLa cells were plated on 35 mm glass-bottomed dishes (Iwaki Glass) and cultured for 24 h at 37 °C in 5% CO_2_. The cells were cotransfected with pCMV-PB1-AG-FRB and pCMV-FKBP-mCherry using 293fectin. After incubation for 24 h, the medium was changed to serum-free DMEM supplemented with penicillin (100 U/mL) and streptomycin (100 μg/mL) [DMEM(−)], and cells were observed by time-lapse imaging before and after the addition of rapamycin (200 nM) dissolved in DMSO (final DMSO concentration <0.1% v/v). Control experiments were performed in the same manner.

### SPREC-Out assay

To conduct the rapamycin-induced protein release assay, 1 × 10^5^ HeLa cells were plated on 35 mm glass-bottomed dishes (Iwaki Glass) and cultured for 24 h at 37 °C in 5% CO_2_. The cells were cotransfected with pCMV-PB1-AG, pCMV-PB1-AG-FRB-TEVCS-BFP-HA and pCMV-mCherry-FKBP-TEVp using 293fectin. After incubation for 24 h, the medium was changed to DMEM(–), and cells were observed by time-lapse imaging before and after the addition of rapamycin (200 nM). Control experiments were performed in the same manner.

### Western blotting

Rapamycin-induced TEVcs cleavage in SPREC-Out systems was evaluated by western blotting as follows. HeLa cells were plated at 2 × 10^5^ cells on 40 mm plastic dishes (TPP) and cultured for 24 h at 37 °C in 5% CO_2_. The cells were cotransfected with the indicated plasmids using 293fectin and treated as described for SPREC-Out experiments. After rapamycin (or mock) treatment for 30 min, the cells were washed twice with cold phosphate-buffered saline and lysed at 4 °C (on ice) in RIPA buffer (Wako) containing 5 mM EDTA and protease inhibitor cocktail (Thermo Fisher Scientific). The cell lysates were cleared by centrifugation, mixed with 2× SDS-PAGE sample buffer and heated at 98 °C for 5 min. The samples were resolved on 7.5% SDS-PAGE gels and transferred to nitrocellulose membranes (Bio-Rad Laboratories). After blocking with 5% skimmed milk in Tris-buffered saline containing 0.1% Tween-20 (TBST), the membranes were probed with appropriate antibodies. TBST was used for all washing steps. The immunoblots were developed with SuperSignal West Femto Maximum Sensitivity Substrate (Thermo Fisher Scientific) and detected using a ChemiDoc MP system (Bio-Rad Laboratories). Antibodies used were: primary antibodies, anti-HA-Tag rabbit mAb (C29F4) (Cell Signaling Technology), anti-monomeric Azami-Green 1 rabbit pAb (MBL), anti-RFP mouse mAb (MBL) and anti-GAPDH rabbit mAb (14C10) (Cell Signaling Technology); secondary antibodies, anti-rabbit IgG HRP-linked antibody and anti-mouse IgG HRP-linked antibody (Cell Signaling Technology).

### Chemogenetic control of Vav2 activity

For chemogenetic Vav2 activity control experiments, 0.6 × 10^5^ HeLa cells stably expressing Lifeact-mTagBFP2 (for SPREC-In) or Lifeact-tdiRFP670 (for SPREC-Out) were plated on 35 mm glass-bottomed dishes coated with 50 μg/mL fibronectin (Corning) and cultured for 24 h at 37 °C in 5% CO_2_. The cells were cotransfected with the following plasmids using 293fectin: for SPREC-In experiments, pCMV-PB1-AG-FRB and pCMV-FKBP-mCherry-Vav2_cat_; and for SPREC-Out experiments, pCMV-PB1-AG, pCMV-PB1-AG-FRB-TEVCS-BFP-HA-Vav2_cat_ and pCMV-mCherry-FKBP-TEVp. After incubation for 24 h, the medium was changed to DMEM(−), and cells were observed by time-lapse imaging before and after the addition of rapamycin (200 nM). Morphological changes were monitored by fluorescence imaging of Lifeact-mTagBFP2 or Lifeact-tdiRFP670 and quantified using QuimP^66^, an ImageJ plugin developed at the University of Warwick. Control experiments were performed in the same manner.

### Chemogenetic control of ERK activity

For chemogenetic ERK activity control experiments, 1.2 × 10^5^ HeLa cells stably expressing MEK1-P2A-mCherry-ERK2 were plated on 35 mm glass-bottomed dishes coated with 0.3 mg/mL collagen (Cell matrix Type I-C, Nitta Gelatin) and cultured for 24 h at 37 °C in 5% CO_2_. The cells were cotransfected with the following plasmids using 293fectin: for SPREC-In experiments, pCMV-PB1-AG-FRB and pCMV-FKBP-mCherry-SOScat; and for SPREC-Out experiments, pCMV-PB1-AG, pCMV-PB1-AG-FRB-TEVCS-BFP-HA-SOS_cat_ and pCMV-mCherry-FKBP-TEVp. After incubation for 24 h, the medium was changed to DMEM(–), and cells were observed by time-lapse imaging before and after the addition of rapamycin (200 nM). Control experiments were performed in the same manner. For MEK inhibition experiments, cells were preincubated with PD184352 (50 μM) (AdooQ BioScience) for 1 h before rapamycin addition.

### optoSPREC assay

To conduct the light-induced protein release assay, 1 × 10^5^ HeLa cells were plated on 35 mm glass-bottomed dishes and cultured for 24 h at 37 °C in 5% CO_2_. The cells were cotransfected with pCAGGS-PB1-MontiRed-LOV2 and pCMV-miRFP670-Zdk1 using 293fectin. After incubation for 24 h, the medium was changed to DMEM(–), and cells were observed by time-lapse imaging under alternating periods of darkness and blue light illumination. Blue light was illuminated from above the dish using a Xenon light source MAX-303 (Asahi Spectra) with 440 nm band-pass filter (LX0440, Asahi Spectra). Control experiments were performed in the same manner.

### Optogenetic control of Vav2 activity

For optogenetic Vav2 activity control using the optoSPREC system, 0.6 × 10^5^ HeLa cells stably expressing Lifeact-tdiRFP670 were plated on 35 mm glass-bottomed dishes coated with 50 μg/mL fibronectin (Corning) and cultured for 24 h at 37 °C in 5% CO_2_. The cells were cotransfected with pCAGGS-PB1-MontiRed-LOV2 and pCMV-miRFP670-Zdk1-Vav2_cat_. After incubation for 24 h, the medium was changed to DMEM(–), and cells were observed by time-lapse imaging under alternating periods of darkness and blue light illumination, as described above. Control experiments were performed in the same manner.

## Acknowledgments

We thank W. D. Heo (KAIST) for the Vav2 plasmid. This work was supported by JSPS Grants-in-Aid for Scientific Research (KAKENHI) (Grant Nos. 15H05459 “Resonance Bio”, 18H04546 and 20H04706 “Chemistry for Multimolecular Crowding Biosystems”, and 20K21250) and the Takeda Science Foundation (to S.T.). This work was also supported by MEXT Leading Initiative for Excellent Young Researchers, JST PRESTO (JPMJPR178B), JSPS KAKENHI (Grant Nos. 17H06759 and 19K15697), the Asahi Glass Foundation, the Nakatani Foundation and the Takeda Science Foundation (to T.Y.).

## Author Contributions

S.T. directed the project. T.Y., M.Y. and S.T. designed the experiments. T.Y., M.Y. and M.I. performed the experiments. All authors analyzed the data. T.Y., M.Y. and S.T. wrote the manuscript.

