## Supplementary Information for "Synthetic protein condensates that recruit and release protein activity in living cells"

### Supplementary Figures

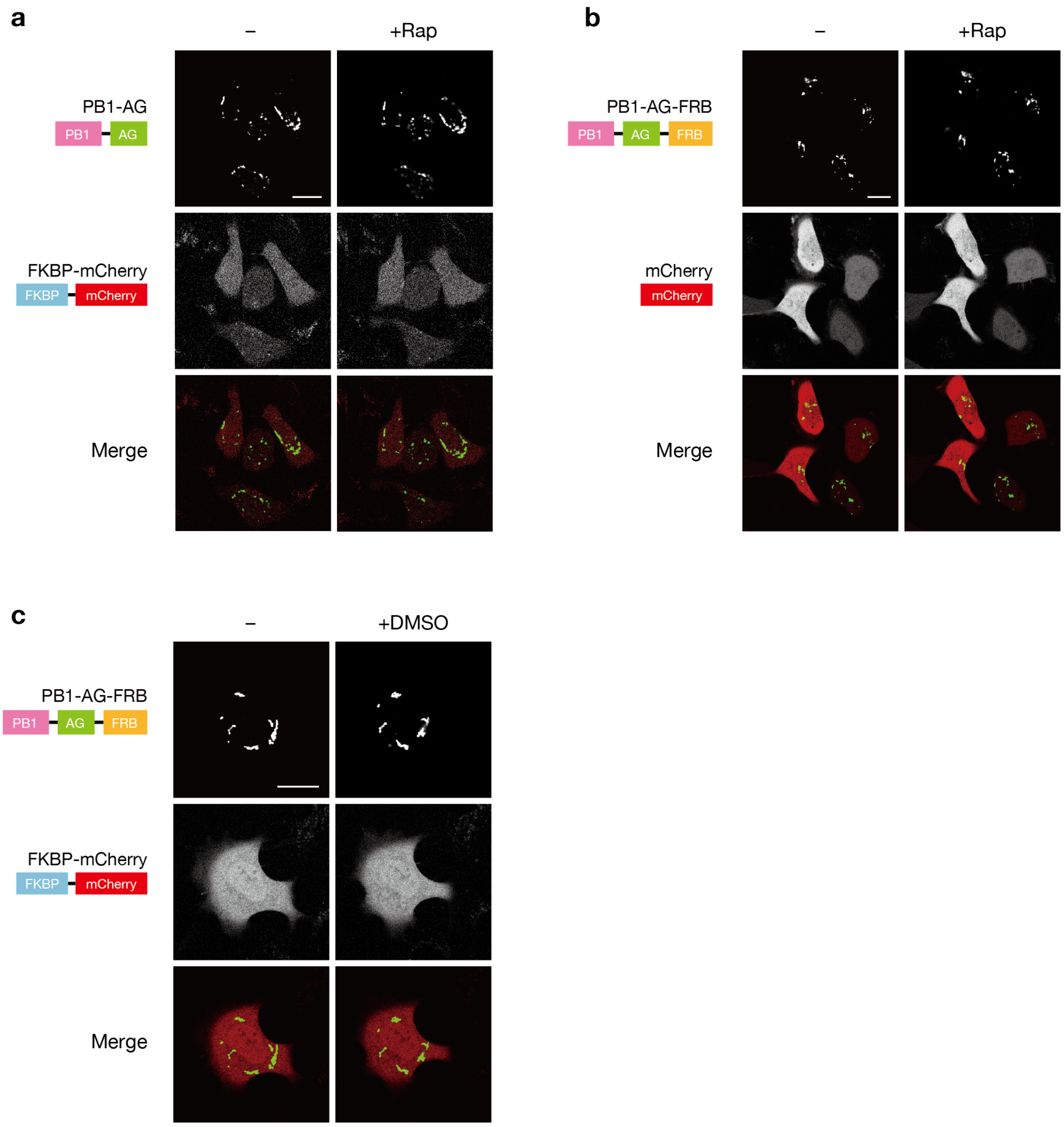


**Supplementary Fig. 1 Control experiments for rapamycin-triggered FKBP-mCherry recruitment to $^{FRB}$PAC (Fig. 1b,c).** **a** Without FRB. **b** Without FKBP. **c** DMSO treatment. Confocal fluorescence images of HeLa cells coexpressing the indicated constructs before and 20 min after incubation with 200 nM rapamycin or DMSO. Scale bars, 20 µm.

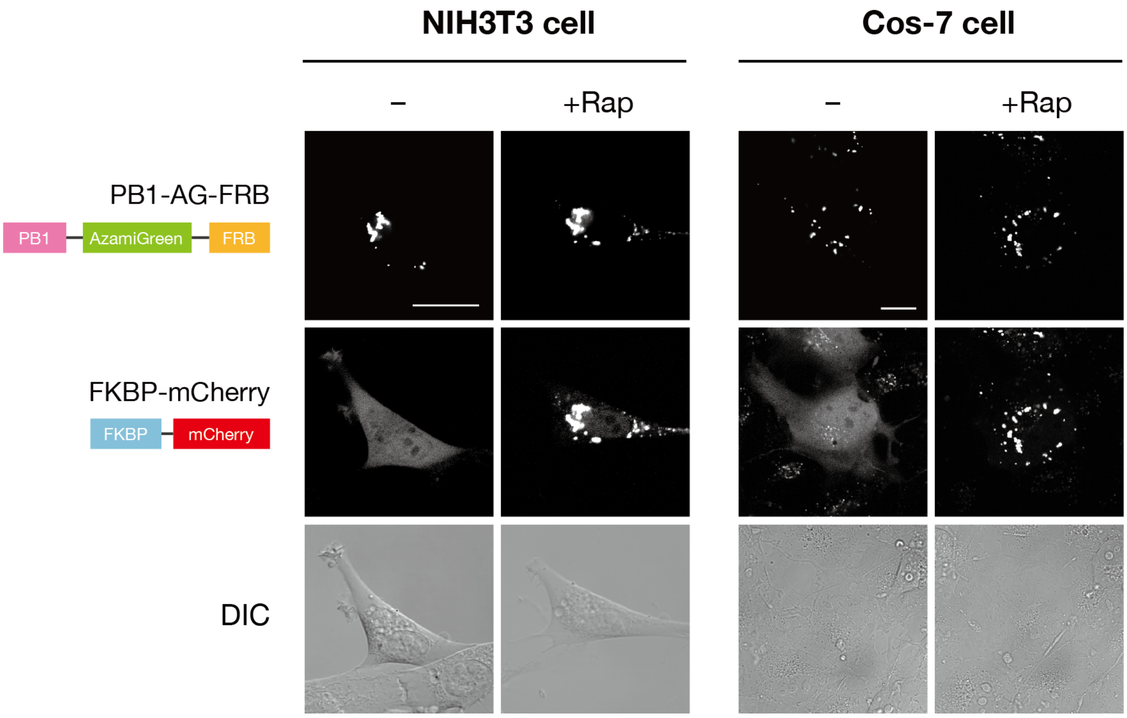


**Supplementary Fig. 2 The use of the SPREC-In system in other cell lines. a** NIH3T3 cells. **b** Cos-7 cells. Confocal fluorescence images of cells coexpressing $^{FRB}$PAC (PB1-AG-FRB) and FKBP-mCherry before and 30 min after incubation with 200 nM rapamycin. Scale bars, 20 µm.

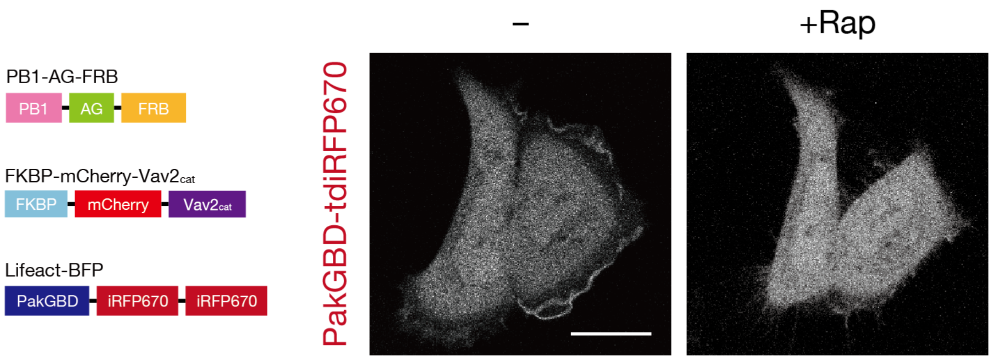


**Supplementary Fig. 3 Monitoring of endogenous Rac activity upon Vav2$_{cat}$ sequestration using the SPREC-In system.** Confocal fluorescence (iRFP) images of HeLa cells coexpressing $^{FRB}$PAC (PB1-AG-FRB), FKBP-mCherry-Vav2$_{cat}$ and PakRBD-tdiRFP670 (a fluorescent reporter for active Rac in a GTP-bound form) before and 20 min after incubation with rapamycin (200 nM). Localized iRFP signals observed at the cell periphery were attenuated after rapamycin treatment, indicating suppression of the endogenous Rac activity. Scale bar, 20 µm.

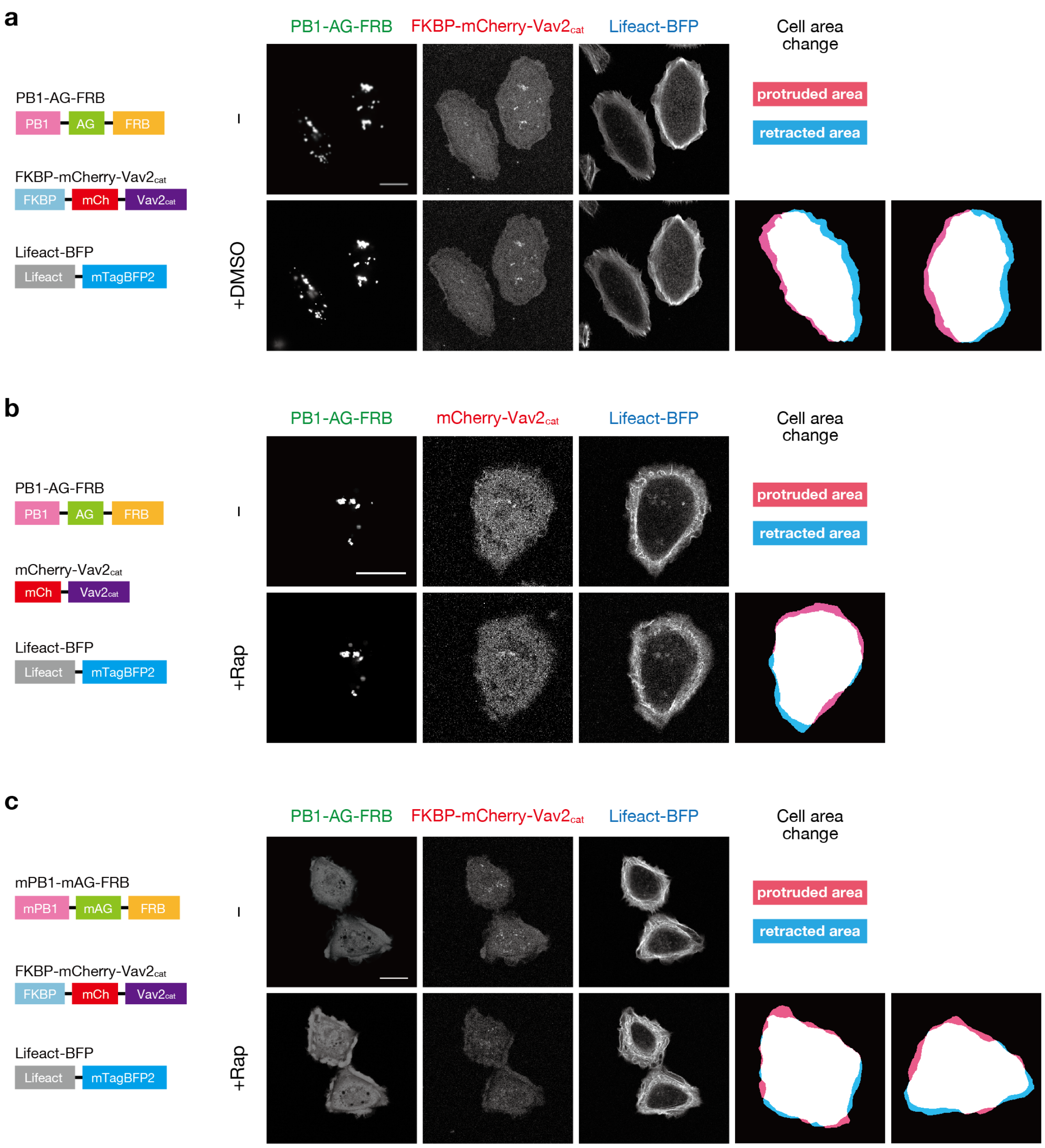


**Supplementary Fig. 4 Control experiments for Vav2 activity sequestration using the SPREC-In system (Fig. 2a–c)**. **a** DMSO treatment. **b** Without FKBP. **c** Use of a monomeric PB1-AG-FRB mutant (mPB1-mAG-FRB). Confocal fluorescence images of HeLa cells coexpressing the indicated constructs before and 20 min after incubation with 200 nM rapamycin or DMSO. Scale bars, 20 µm.

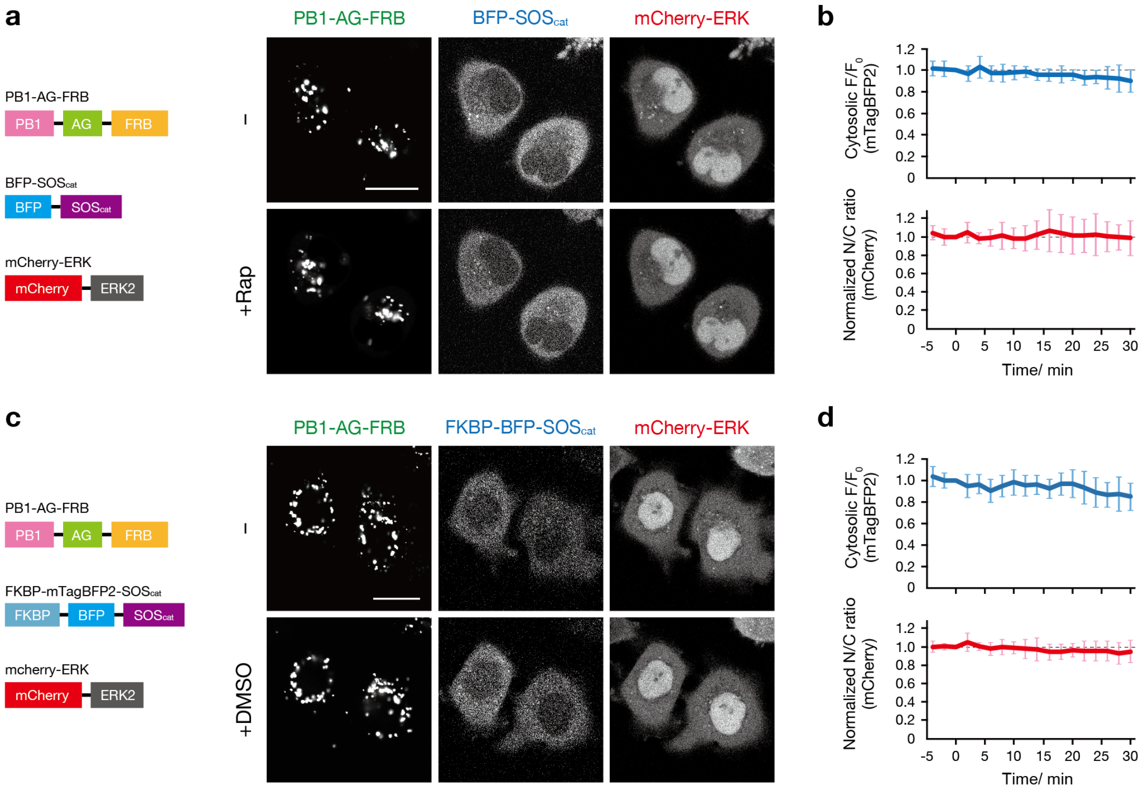


**Supplementary Fig. 5 Control experiments for SOS activity sequestration using the SPREC-In system** (**Fig. 2d,e**). **a** Without FKBP. **b** DMSO treatment. Confocal fluorescence images of HeLa cells coexpressing the indicated constructs before and 30 min after incubation with 200 nM rapamycin or DMSO. $SOS_{cat}$ recruitment and ERK activity were evaluated as described in the legend of **Fig. 2e**. Scale bars, 20 μm.

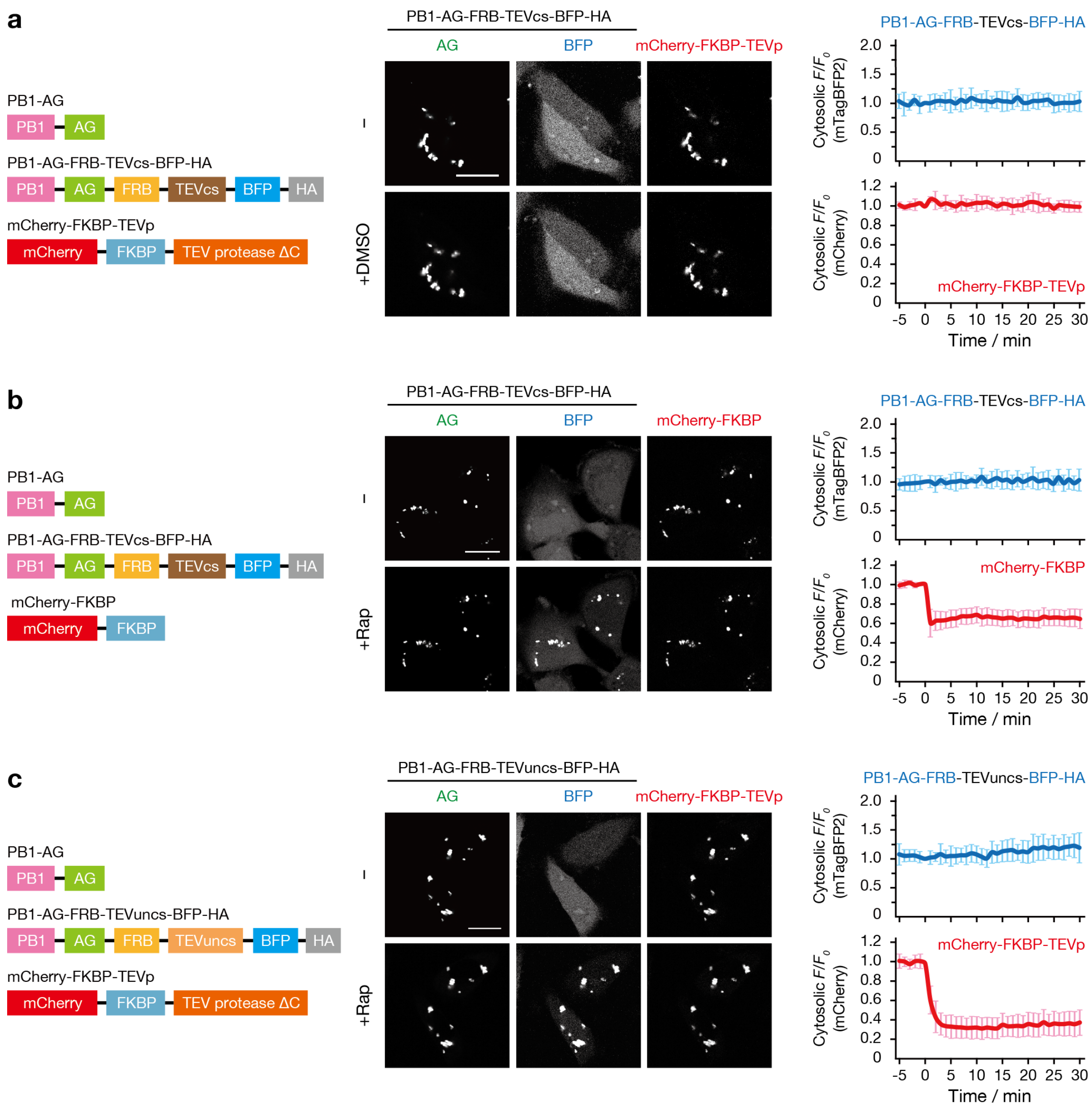


**Supplementary Fig. 6 Control experiments for rapamycin-triggered BFP-HA release using the SPREC-Out system (Fig. 3b,c). a** DMSO treatment. **b** Without TEVp. **c** Use of the uncleavable TEVcs (TEVuncs). Confocal fluorescence images of HeLa cells coexpressing the indicated constructs before and 30 min after incubation with 200 nM rapamycin or DMSO. TEVp recruitment and BFP release were evaluated as described in the legend of **Fig. 3c**. Scale bars, 20 µm.

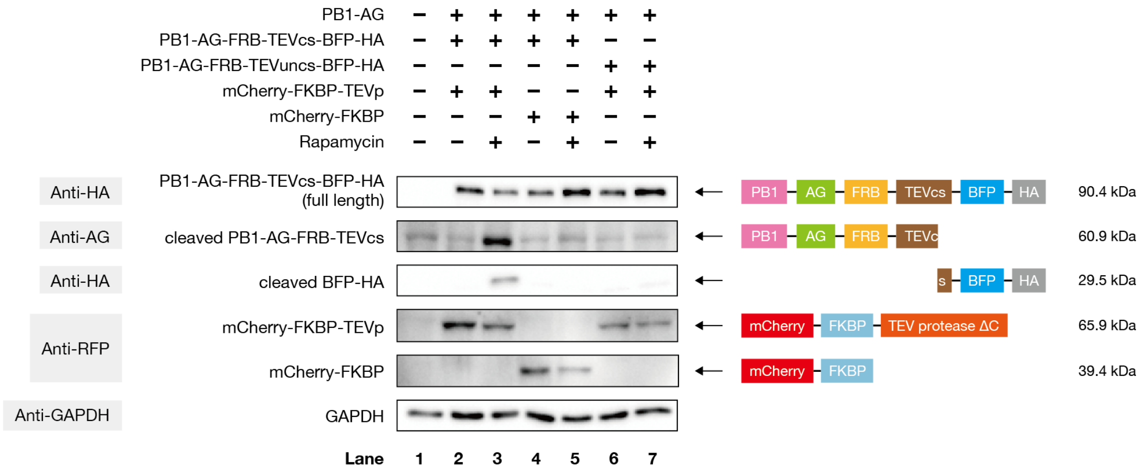


**Supplementary Fig. 7 Western blotting analysis of BFP-HA release using the SPREC-Out system.** HeLa cells coexpressing the indicated constructs were lysed after 30 min incubation with 200 nM rapamycin or without rapamycin treatment. The cell lysates were immunoblotted with the indicated antibodies. Rapamycin- and TEVp-dependent cleavage of PB1-AG-FRB-TEVcs-BFP-HA (90.4 kDa), producing cleavage fragments PB1-AG-FRB-TEVcs (60.9 kDa) and BFP-HA (29.5 kDa), was observed in lane 3, which contained all the components required for the SPREC-Out system.

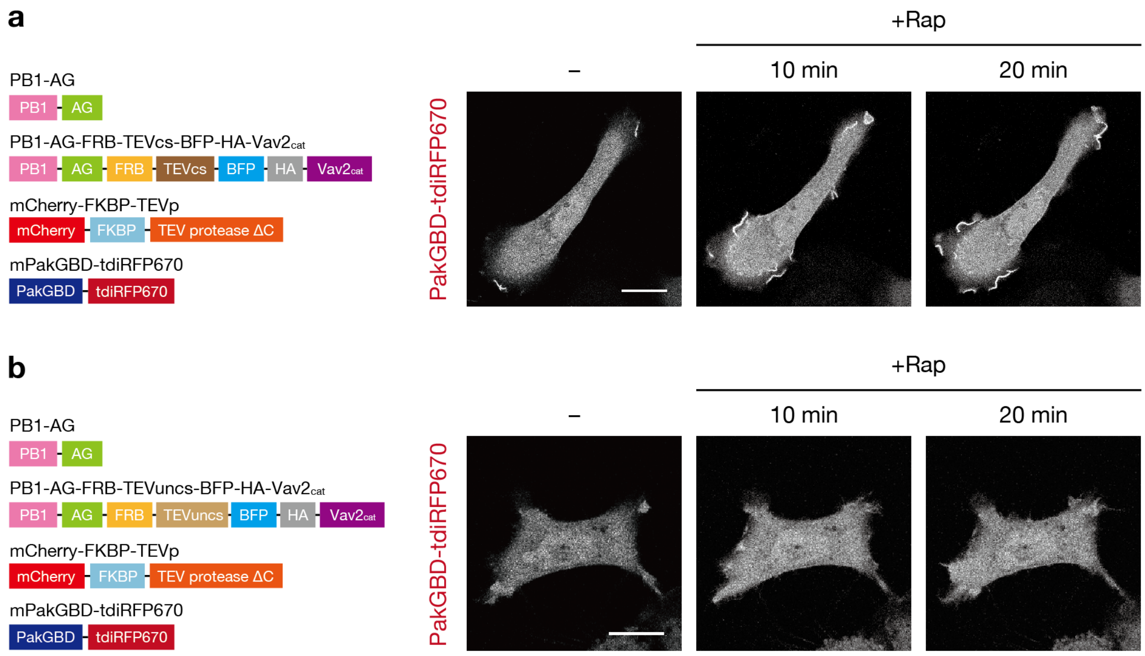


**Supplementary Fig. 8 Monitoring of endogenous Rac activity upon Vav2$_{cat}$ release using the SPREC-Out system.** **a** Time-lapse confocal fluorescence (iRFP) images of HeLa cells coexpressing $^{\text{FRB-TCS-BFP-HA-Vav2}}$PAC (formed by coassembly of PB1-AG and PB1-AG-FRB-TEVcs-BFP-HA-Vav2$_{cat}$), mCherry-FKBP-TEVp and PakGBD-tdiRFP670 (a fluorescent reporter for active Rac) before and 10 and 20 min after incubation with rapamycin (200 nM). Localized iRFP signals at the cell periphery were observed after rapamycin treatment, indicating endogenous Rac activation. **b** Negative control using the uncleavable TEVuncs. Scale bars, 20 µm.

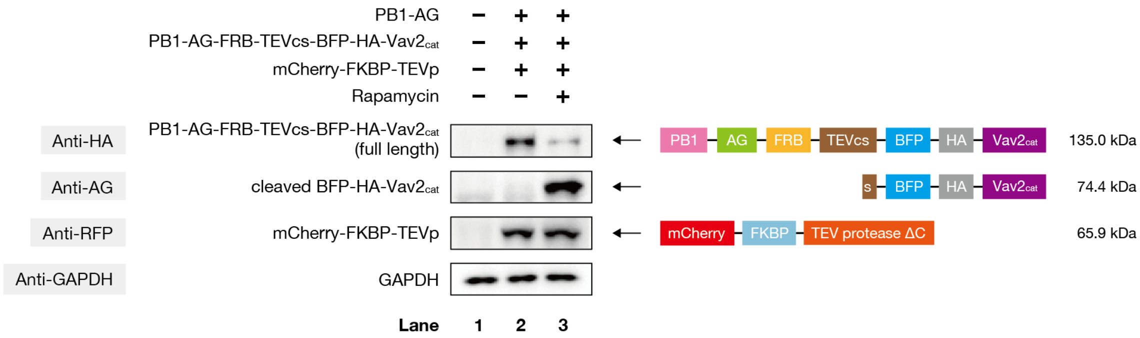


**Supplementary Fig. 9 Western blotting analysis of the Vav2$_{cat}$-releasing SPREC-Out system.** HeLa cells coexpressing $^{\text{FRB-TCS-BFP-HA-Vav2}}$PAC (PB1-AG and PB1-AG-FRB-TEVcs-BFP-HA-Vav2$_{cat}$) and mCherry-FKBP-TEVp were lysed after 30 min incubation with 200 nM rapamycin or without rapamycin treatment. The cell lysates were immunoblotted with the indicated antibodies. The cleaved product BFP-HA-Vav2$_{cat}$ (74.4 kDa) was observed in lane 3.

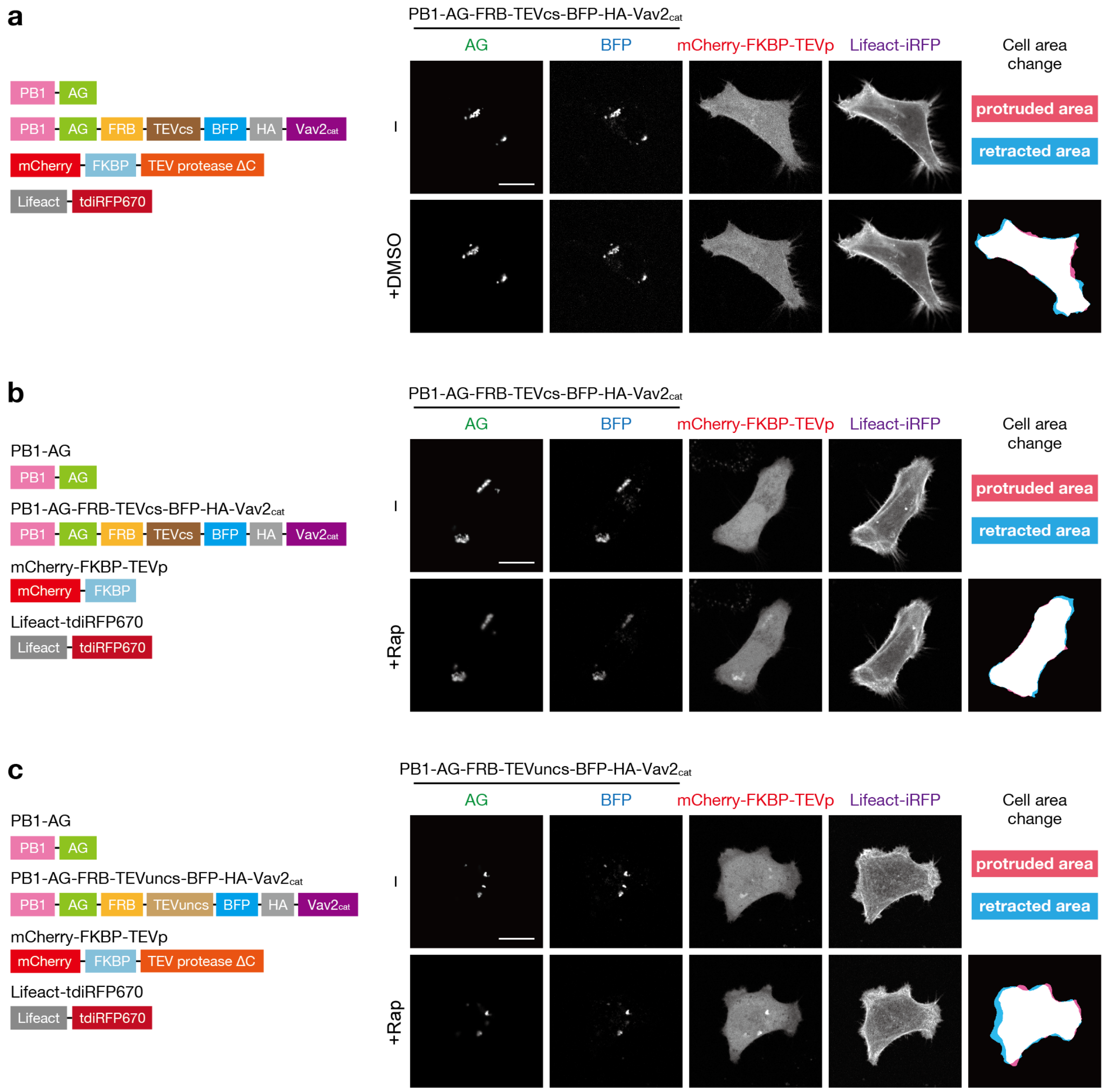


**Supplementary Fig. 10 Control experiments for Vav2 activity release using the SPREC-Out system (Fig. 4a–c).** **a** DMSO treatment. **b** Without TEVp. **c** Use of the uncleavable TEVuncs. Confocal fluorescent images of HeLa cells coexpressing the indicated constructs before and 20 min after incubation with 200 nM rapamycin. Scale bars, 20 µm.

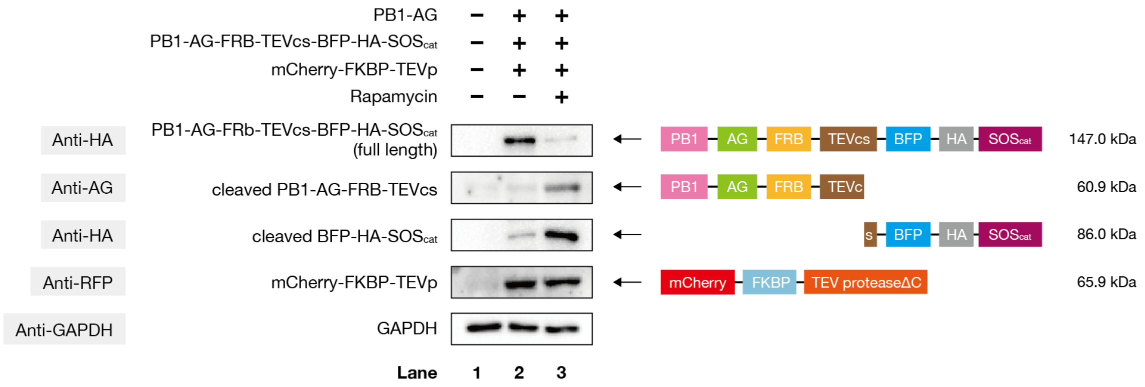


**Supplementary Fig. 11 Western blotting analysis of the SOS$_{cat}$-releasing SPREC-Out system.** HeLa cells coexpressing $^{FRB-TCS-BFP-HA-SOS}$PAC (PB1-AG and PB1-AG-FRB-TEVcs-BFP-HA-SOS$_{cat}$) and mCherry-FKBP-TEVp were lysed after 30 min incubation with 200 nM rapamycin or without rapamycin treatment. The cell lysates were immunoblotted with the indicated antibodies. The cleaved product BFP-HA-SOS$_{cat}$ (86.0 kDa) was observed in lane 3.

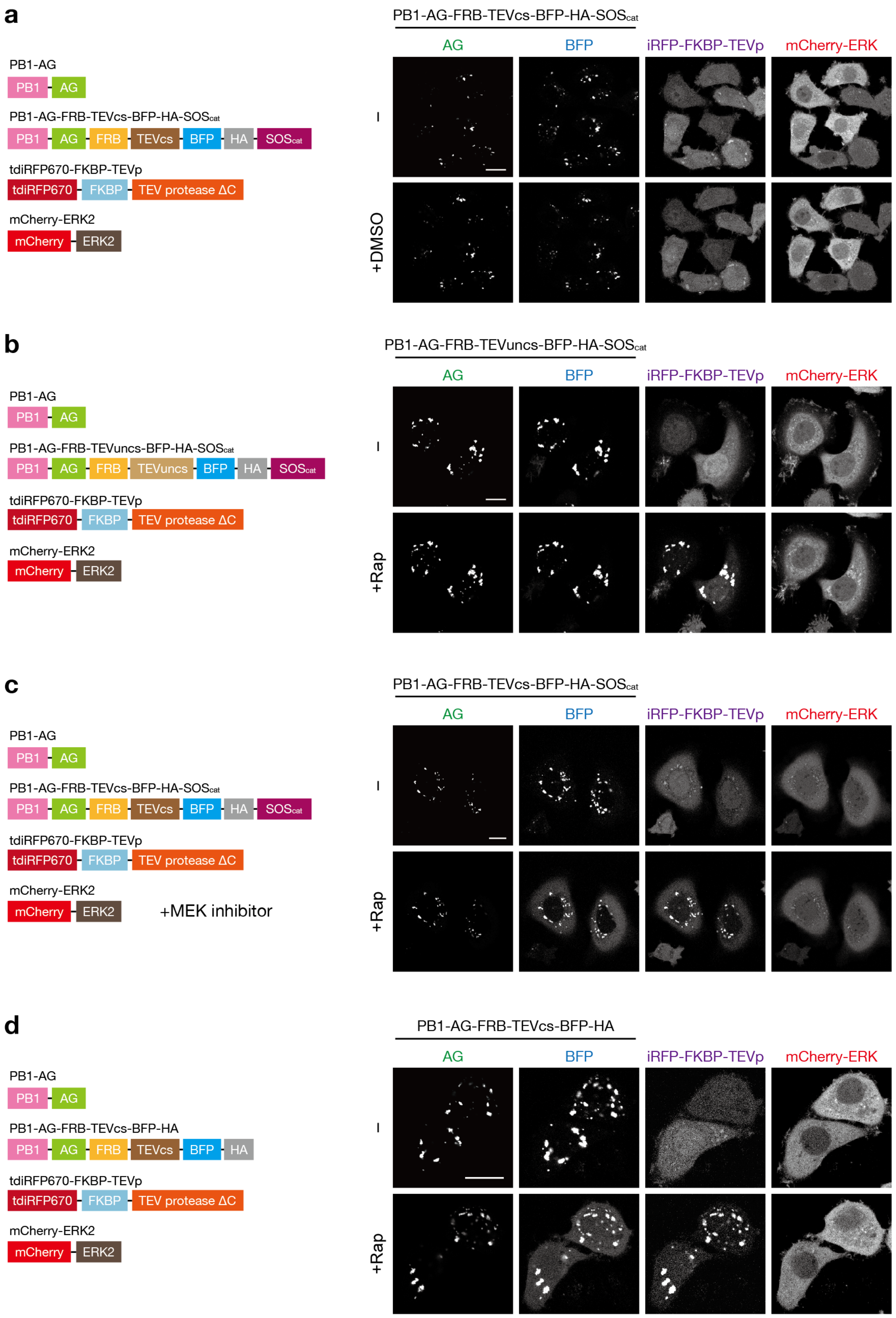


**Supplementary Fig. 12 Control experiments for SOS activity release using the SPREC-Out**

**system** (**Fig. 4d,e**). **a** DMSO treatment. **b** Use of the uncleavable TEVuncs. **c** BFP-HA-$SOS_{cat}$ release in the presence of the MEK inhibitor, PD184352 (50 μM). **d** Release of BFP-HA. Confocal fluorescent images of HeLa cells coexpressing the indicated constructs before and 20 min after incubation with 200 nM rapamycin or DMSO. $SOS_{cat}$ release and ERK activity were evaluated as described in the legend of **Fig. 4e**. Scale bars, 20 μm.

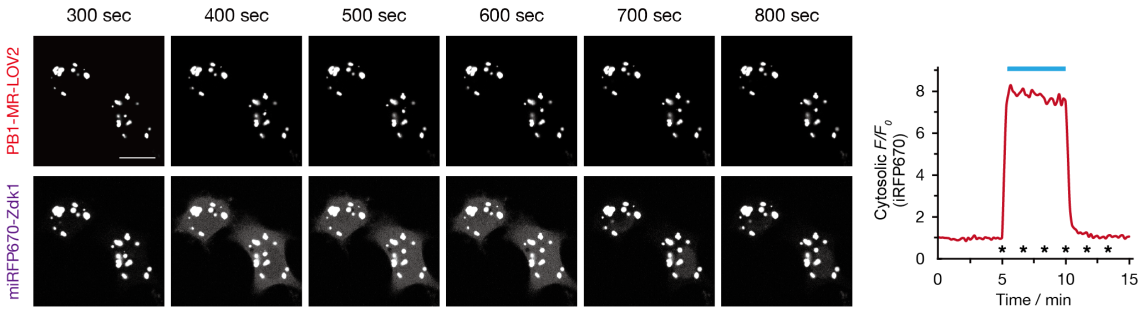


**Supplementary Fig. 13 Prolonged light illumination-induced miRFP-Zdk release.** Time-lapse confocal fluorescence images of HeLa cells coexpressing $^{LOV2}$PAC (PB1-MR-LOV2) and miRFP-Zdk under a cycle of dark and blue-light illumination. Images were taken at time points indicated by asterisks shown in the right graph. Light-induced miRFP-Zdk release was evaluated as described in the legend of **Fig. 5c**. The blue bar indicates the period of blue light illumination. Scale bar, 20 µm.

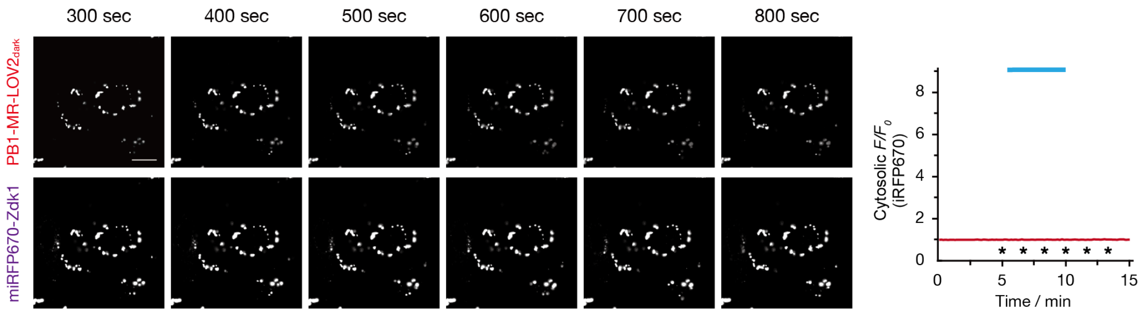


**Supplementary Fig. 14 Control experiments for light-induced miRFP-Zdk release (Fig. 5b)**. Time-lapse confocal fluorescence images of HeLa cells coexpressing PB1-MR-LOV2$_{dark}$ (PB1-AG-LOV2 containing C450A/L514K/G528A/L531E/N538E mutations in LOV2) and miRFP-Zdk under a cycle of dark and blue-light illumination. Images were taken at time points indicated by asterisks shown in the right graph. Light-induced miRFP-Zdk release was evaluated as described in the legend of **Fig. 5c**. The blue bar indicates the period of blue light illumination. Scale bar, 20 µm.

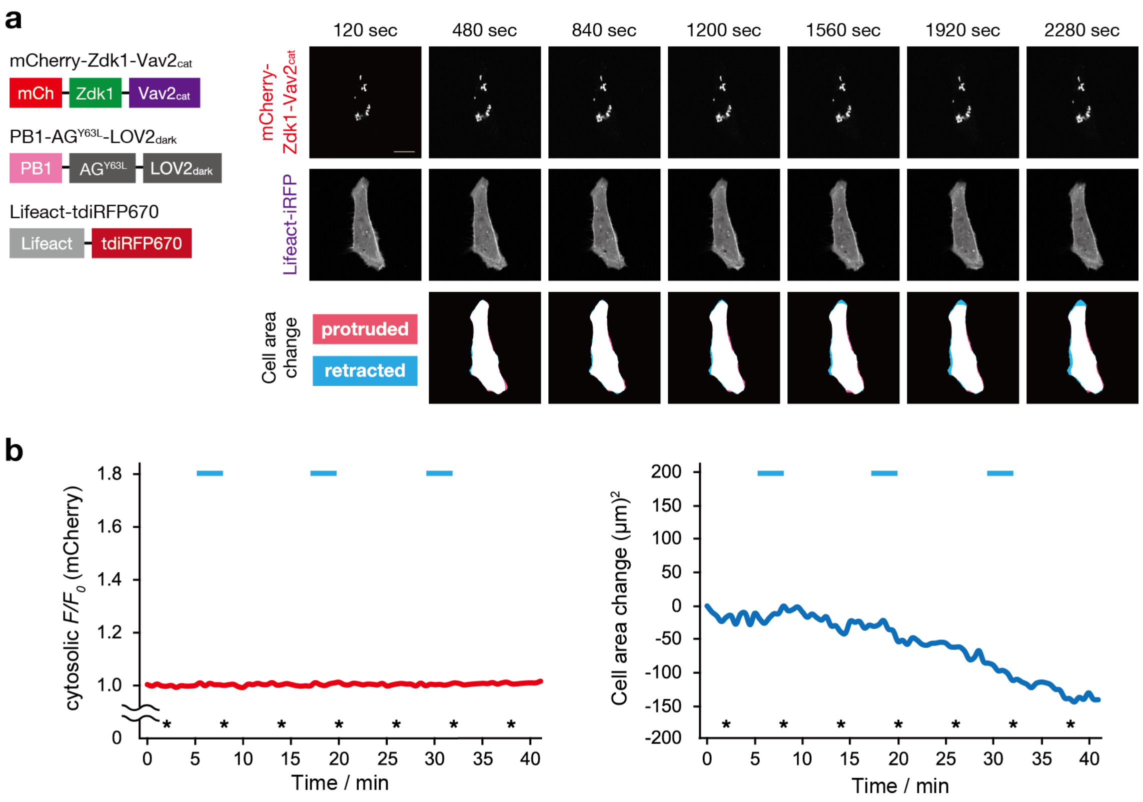


**Supplementary Fig. 15 Control experiments for light-induced release of Vav2 activity using the optoSPREC system** (**Fig. 5e,f**). **a** Time-lapse confocal fluorescent images of a HeLa cell coexpressing non-fluorescent, light-insensitive $^{\mathrm{LOV2(dark)}}$PAC [PB1-AG(Y63L)-LOV2$_{\mathrm{dark}}$], mCherry-Zdk-Vav2$_{\mathrm{cat}}$ and Lifeact-iRFP under alternating periods of darkness and blue light illumination. Images were taken at time points indicated by asterisks shown in **b**. **b** Vav2$_{\mathrm{cat}}$ release and cell area changes were evaluated as described in the legend of **Fig. 5e**. The blue bars indicate the periods of light illumination. Scale bar, 20 μm.

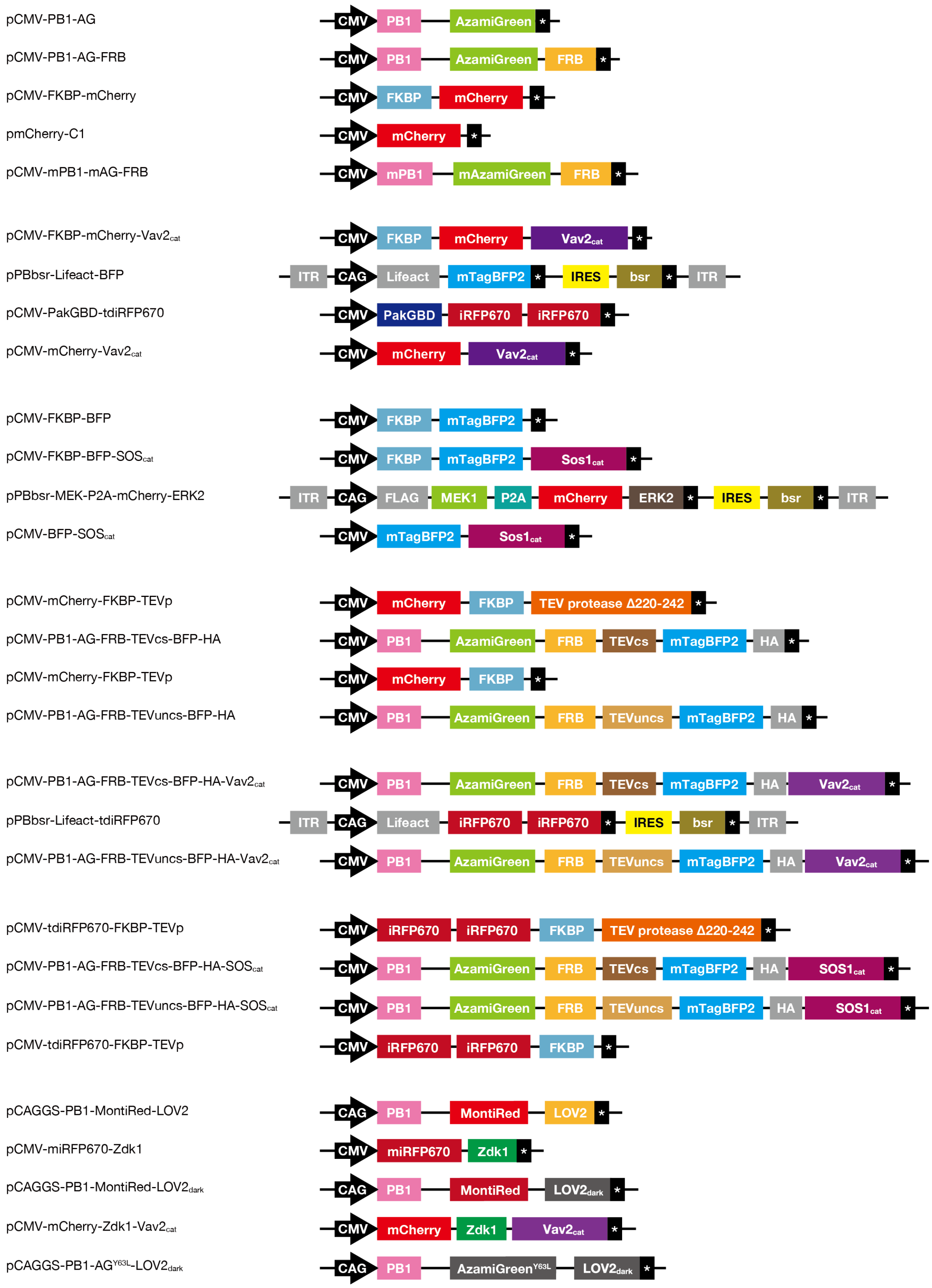


**Supplementary Fig. 16 Schematic illustration of the domain structures of constructs used in this study.**

**a**

**pCMV-PB1-AG**

>Amino acid sequence
MASLTVKAYLLGKEDAAREIRRFSFCCSPEPEAEAEAAAGPGPCERLLSRVAALFPALRPGGFQAHYRDE
DGDLVAFSSDEELTMAMSYVKDDIFRIYIKEKTGSGSGGSGAGGSAGSGAGGSAGSGAGGSAGSGAGGSA
GSGAGGSASGSSPGSGSGGSGAGGSAGSGAGGSAGSGAGGSAGSGAGGSAGSGAGGSASGSSPGNSADGG
GGSGGSGGSGGGSTQGGSMVSVIKPEMKIKLCMRGTVNGHNFVIEGEGKGNPYEGTQILDLNVTEGAPLP
FAYDILTTVFQYGNRAFTKYPADIQDYFKQTFPEGYHWERSMTYEDQGICTATSNISMRGDCFFYDIRFD
GVNFPPNGPVMQKKTLKWEPSTEKMYVRDGVLKGDVNMALLLEGGGHYRCDFKTTYKAKKDVRLPDYHFV
DHRIEILKHDKDYNKVKLYENAVARYSMLPSQAKGTGTAAENSGNSRTKLE*

>Nucleic acid sequence
ATGGCGTCGCTCACCGTGAAGGCCTACCTTCTGGGCAAGGAGGACGCGGCGCGCGAGATTCGCCGCTTCA
GCTTCTGTTGCAGCCCCGAGCCTGAGGCGGAAGCCGAGGCTGCGGCGGGTCCGGGACCCTGCGAGCGGCT
GCTGAGCCGGGTGGCCGCCCTGTTCCCCGCGCTGCGGCCTGGCGGCTTCCAGGCGCACTACCGCGATGAG
GACGGGGACTTGGTTGCCTTTTCCAGTGACGAGGAATTGACAATGGCCATGTCCTACGTGAAGGATGACA
TCTTCCGAATCTACATTAAAGAGAAAACCGGTTCTGGGAGTGGCGGTAGTGGTGCTGGAGGCAGCGCAGG
CTCTGGAGCAGGCGGCAGTGCCGGAAGTGGCGCTGGAGGGTCTGCCGGATCTGGAGCCGGTGGTTCTGCC
GGCTCCGGCGCTGGTGGGAGCGCTTCCGGAAGTAGTCCCGGTTCTGGGAGTGGCGGTAGTGGTGCTGGAG
GCAGCGCAGGCTCTGGAGCAGGCGGCAGTGCCGGAAGTGGCGCTGGAGGGTCTGCCGGATCTGGAGCCGG
TGGTTCTGCCGGCTCCGGCGCTGGTGGGAGCGCTTCCGGAAGTAGTCCCGGTAATTCCGCTGACGGCGGC
GGAGGATCGGGTGGTAGTGGTGGTTCAGGAGGAGGATCGACCCAAGGAGGATCCATGGTGAGCGTGATCA
AGCCCGAGATGAAGATCAAGCTGTGCATGAGGGGCACCGTGAACGGCCACAACTTCGTGATCGAGGGCGA
GGGCAAGGGCAACCCCTACGAGGGCACCCAGATCCTGGACCTGAACGTGACCGAGGGCGCCCCCCTGCCC
TTCGCCTACGACATCCTGACCACCGTGTTCCAGTACGGCAACAGGGCCTTCACCAAGTACCCCGCCGACA
TCCAGGACTACTTCAAGCAGACCTTCCCCGAGGGCTACCACTGGGAGAGGAGCATGACCTACGAGGACCA
GGGCATCTGCACCGCCACCAGCAACATCAGCATGAGGGGCGACTGCTTCTTCTACGACATCAGGTTCGAC
GGCGTGAACTTCCCCCCCAACGGCCCCGTGATGCAGAAGAAGACCCTGAAGTGGGAGCCCAGCACCGAGA
AGATGTACGTGAGGGACGGCGTGCTGAAGGGCGACGTGAACATGGCCCTGCTGCTGGAGGGCGGCGGCCA
CTACAGGTGCGACTTCAAGACCACCTACAAGGCCAAGAAGGACGTGAGGCTGCCCGACTACCACTTCGTG
GACCACAGGATCGAGATCCTGAAGCACGACAAGGACTACAACAAGGTGAAGCTGTACGAGAACGCCGTGG
CCAGGTACAGCATGCTGCCCAGCCAGGCCAAGGGTACCGGAACTGCAGCAGAGAATTCGGGAAACTCGAG
AACAAAGCTTGAATAA

**Annotation**
PB1: PB1 domain from human p62 (residues 1-102)
AG: green fluorescent protein Azami-Green

**b**

**pCMV-PB1-AG-FRB**

>Amino acid sequence
MASLTVKAYLLGKEDAAREIRRFSFCCSPEPEAEAEAAAGPGPCERLLSRVAALFPALRPGGFQAHYRDE
DGDLVAFSSDEELTMAMSYVKDDIFRIYIKEKTGSGSGGSGAGGSAGSGAGGSAGSGAGGSAGSGAGGSA
GSGAGGSASGSSPGSGSGGSGAGGSAGSGAGGSAGSGAGGSAGSGAGGSAGSGAGGSASGSSPGNSADGG
GGSGGSGGSGGGSTQGGSMVSVIKPEMKIKLCMRGTVNGHNFVIEGEGKGNPYEGTQILDLNVTEGAPLP
FAYDILTTVFQYGNRAFTKYPADIQDYFKQTFPEGYHWERSMTYEDQGICTATSNISMRGDCFFYDIRFD
GVNFPPNGPVMQKKTLKWEPSTEKMYVRDGVLKGDVNMALLLEGGGHYRCDFKTTYKAKKDVRLPDYHFV
DHRIEILKHDKDYNKVKLYENAVARYSMLPSQAKGTGTAAENSGNSRTKLMILWHEMWHEGLEEASRLYF
GERNVKGMFEVLEPLHAMMERGPQTLKETSFNQAYGRDLMEAQEWCRKYMKSGNVKDLLQAWDLYYHVFR
RISKPG*

>DNA sequence
ATGGCGTCGCTCACCGTGAAGGCCTACCTTCTGGGCAAGGAGGACGCGGCGCGCGAGATTCGCCGCTTCA
GCTTCTGTTGCAGCCCCGAGCCTGAGGCGGAAGCCGAGGCTGCGGCGGGTCCGGGACCCTGCGAGCGGCT
GCTGAGCCGGGTGGCCGCCCTGTTCCCCGCGCTGCGGCCTGGCGGCTTCCAGGCGCACTACCGCGATGAG
GACGGGGACTTGGTTGCCTTTTCCAGTGACGAGGAATTGACAATGGCCATGTCCTACGTGAAGGATGACA
TCTTCCGAATCTACATTAAAGAGAAAACCGGTTCTGGGAGTGGCGGTAGTGGTGCTGGAGGCAGCGCAGG
CTCTGGAGCAGGCGGCAGTGCCGGAAGTGGCGCTGGAGGGTCTGCCGGATCTGGAGCCGGTGGTTCTGCC
GGCTCCGGCGCTGGTGGGAGCGCTTCCGGAAGTAGTCCCGGTTCTGGGAGTGGCGGTAGTGGTGCTGGAG
GCAGCGCAGGCTCTGGAGCAGGCGGCAGTGCCGGAAGTGGCGCTGGAGGGTCTGCCGGATCTGGAGCCGG
TGGTTCTGCCGGCTCCGGCGCTGGTGGGAGCGCTTCCGGAAGTAGTCCCGGTAATTCCGCTGACGGCGGC
GGAGGATCGGGTGGTAGTGGTGGTTCAGGAGGAGGATCGACCCAAGGAGGATCCATGGTGAGCGTGATCA
AGCCCGAGATGAAGATCAAGCTGTGCATGAGGGGCACCGTGAACGGCCACAACTTCGTGATCGAGGGCGA
GGGCAAGGGCAACCCCTACGAGGGCACCCAGATCCTGGACCTGAACGTGACCGAGGGCGCCCCCCTGCCC
TTCGCCTACGACATCCTGACCACCGTGTTCCAGTACGGCAACAGGGCCTTCACCAAGTACCCCGCCGACA
TCCAGGACTACTTCAAGCAGACCTTCCCCGAGGGCTACCACTGGGAGAGGAGCATGACCTACGAGGACCA
GGGCATCTGCACCGCCACCAGCAACATCAGCATGAGGGGCGACTGCTTCTTCTACGACATCAGGTTCGAC
GGCGTGAACTTCCCCCCCAACGGCCCCGTGATGCAGAAGAAGACCCTGAAGTGGGAGCCCAGCACCGAGA
AGATGTACGTGAGGGACGGCGTGCTGAAGGGCGACGTGAACATGGCCCTGCTGCTGGAGGGCGGCGGCCA
CTACAGGTGCGACTTCAAGACCACCTACAAGGCCAAGAAGGACGTGAGGCTGCCCGACTACCACTTCGTG
GACCACAGGATCGAGATCCTGAAGCACGACAAGGACTACAACAAGGTGAAGCTGTACGAGAACGCCGTGG
CCAGGTACAGCATGCTGCCCAGCCAGGCCAAGGGTACCGGAACTGCAGCAGAGAATTCGGGAAACTCGAG
AACAAAGCTTATGATCCTCTGGCATGAGATGTGGCATGAAGGCCTGGAAGAGGCATCTCGTTTGTACTTT
GGGGAAAGGAACGTGAAAGGCATGTTTGAGGTGCTGGAGCCCTTGCATGCTATGATGGAACGGGGCCCCC
AGACTCTGAAGGAAACATCCTTTAATCAGGCCTATGGTCGAGATTTAATGGAGGCCCAAGAGTGGTGCAG
GAAGTACATGAAATCAGGGAATGTCAAGGACCTCCTCCAAGCCTGGGACCTCTATTATCATGTGTTCCGA
CGAATCTCAAAGCCCGGGTAA

**Annotation**
PB1: PB1 domain from human p62 (residues 1-102)
AG: green fluorescent protein Azami-Green
FRB: FRB domain from human mammalian target of rapamycin (mTOR) (residues 2021-2113 with a T2098L mutation)

**c**

**pCMV-FKBP-mCherry**

```
>Amino acid sequence
MGVQVETISPGDGRTFPKRGQTCVVHYTGMLEDGKKFDSSRDRNKPFKFMLGKQEVIRGWEEGVAQMSVG
QRAKLTISPDYAYGATGHPGIIPPHATLVFDVELLKLEGSGASAPVATMVSKGEEDNMAIIKEFMRFKVH
MEGSVNGHEFEIEGEGEGRPYEGTQTAKLKVTKGGPLPFAWDILSPQFMYGSKAYVKHPADIPDYLKLSF
PEGFKWERVMNFEDGGVVTVTQDSSLQDGEFIYKVKLRGTNFPSDGPVMQKKTMGWEASSERMYPEDGAL
KGEIKQRLKLKDGGHYDAEVKTTYKAKKPVQLPGAYNVNIKLDITSHNEDYTIVEQYERAEGRHSTGGMD
ELYKSGLRSRAQASNSAVDGTAGPGSTGSR*

>DNA sequence
ATGGGAGTGCAGGTGGAAACCATCTCCCCAGGAGACGGGCGCACCTTCCCCAAGCGCGGCCAGACCTGCG
TGGTGCACTACACCGGGATGCTTGAAGATGGAAAGAAATTTGATTCCTCCCGGGACAGAAACAAGCCCTT
TAAGTTTATGCTAGGCAAGCAGGAGGTGATCCGAGGCTGGGAAGAAGGGGTTGCCCAGATGAGTGTGGGT
CAGAGAGCCAAACTGACTATATCTCCAGATTATGCCTATGGTGCCACTGGGCACCCAGGCATCATCCCAC
CACATGCCACTCTCGTCTTCGATGTGGAGCTTCTAAAACTGGAAGGCTCCGGTGCCAGTGCACCGGTCGC
CACCATGGTGAGCAAGGGCGAGGAGGATAACATGGCCATCATCAAGGAGTTCATGCGCTTCAAGGTGCAC
ATGGAGGGCTCCGTGAACGGCCACGAGTTCGAGATCGAGGGCGAGGGCGAGGGCCGCCCCTACGAGGGCA
CCCAGACCGCCAAGCTGAAGGTGACCAAGGGTGGCCCCCTGCCCTTCGCCTGGGACATCCTGTCCCCTCA
GTTCATGTACGGCTCCAAGGCCTACGTGAAGCACCCCGCCGACATCCCCGACTACTTGAAGCTGTCCTTC
CCCGAGGGCTTCAAGTGGGAGCGCGTGATGAACTTCGAGGACGGCGGCGTGGTGACCGTGACCCAGGACT
CCTCCCTGCAGGACGGCGAGTTCATCTACAAGGTGAAGCTGCGCGGCACCAACTTCCCCTCCGACGGCCC
CGTAATGCAGAAGAAGACCATGGGCTGGGAGGCCTCCTCCGAGCGGATGTACCCCGAGGACGGCGCCCTG
AAGGGCGAGATCAAGCAGAGGCTGAAGCTGAAGGACGGCGGCCACTACGACGCTGAGGTCAAGACCACCT
ACAAGGCCAAGAAGCCCGTGCAGCTGCCCGGCGCCTACAACGTCAACATCAAGTTGGACATCACCTCCCA
CAACGAGGACTACACCATCGTGGAACAGTACGAACGCGCCGAGGGCCGCCACTCCACCGGCGGCATGGAC
GAGCTGTACAAGTCCGGACTCAGATCTCGAGCTCAAGCTTCGAATTCTGCAGTCGACGGTACCGCGGGCC
CGGGATCCACCGGATCTAGATAA
```

**Annotation**

FKBP: FKBP12

mCherry: red fluorescent protein mCherry

**d**

**pCMV-mCherry**

>Amino acid sequence
MVSKGEEDNMAIIKEFMRFKVHMEGSVNGHEFEIEGEGEGRPYEGTQTAKLKVTKGGPLPFAWDILSPQF
MYGSKAYVKHPADIPDYLKLSFPEGFKWERVMNFEDGGVVTVTQDSSLQDGEFIYKVKLRGTNFPSDGPV
MQKKTMGWEASSERMYPEDGALKGEIKQRLKLKDGGHYDAEVKTTYKAKKPVQLPGAYNVNIKLDITSHN
EDYTIVEQYERAEGRHSTGGMDELYKSGLRSRAQASNSAVDGTAGPGSTGSR*

>DNA sequence
ATGGTGAGCAAGGGCGAGGAGGATAACATGGCCATCATCAAGGAGTTCATGCGCTTCAAGGTGCACATGG
AGGGCTCCGTGAACGGCCACGAGTTCGAGATCGAGGGCGAGGGCGAGGGCCGCCCCTACGAGGGCACCCA
GACCGCCAAGCTGAAGGTGACCAAGGGTGGCCCCCTGCCCTTCGCCTGGGACATCCTGTCCCCTCAGTTC
ATGTACGGCTCCAAGGCCTACGTGAAGCACCCCGCCGACATCCCCGACTACTTGAAGCTGTCCTTCCCCG
AGGGCTTCAAGTGGGAGCGCGTGATGAACTTCGAGGACGGCGGCGTGGTGACCGTGACCCAGGACTCCTC
CCTGCAGGACGGCGAGTTCATCTACAAGGTGAAGCTGCGCGGCACCAACTTCCCCTCCGACGGCCCCGTA
ATGCAGAAGAAGACCATGGGCTGGGAGGCCTCCTCCGAGCGGATGTACCCCGAGGACGGCGCCCTGAAGG
GCGAGATCAAGCAGAGGCTGAAGCTGAAGGACGGCGGCCACTACGACGCTGAGGTCAAGACCACCTACAA
GGCCAAGAAGCCCGTGCAGCTGCCCGGCGCCTACAACGTCAACATCAAGTTGGACATCACCTCCCACAAC
GAGGACTACACCATCGTGGAACAGTACGAACGCGCCGAGGGCCGCCACTCCACCGGCGGCATGGACGAGC
TGTACAAGTCCGGACTCAGATCTCGAGCTCAAGCTTCGAATTCTGCAGTCGACGGTACCGCGGGCCCGGG
ATCCACCGGATCTAGATAA

**Annotation**
mCherry: red fluorescent protein mCherry

**e**

**pCMV-mPB1-mAG-FRB**

>Amino acid sequence
MASLTVKAYLLGKEDAAREIRRFSFCCSPEPEAEAEAAAGPGPCERLLSRVAALFPALRPGGFQAHYRAE
RGDLVAFSSDEELTMAMSYVKDDIFRIYIKEKTGSGSGGSGAGGSAGSGAGGSAGSGAGGSAGSGAGGSA
GSGAGGSASGSSPGSGSGGSGAGGSAGSGAGGSAGSGAGGSAGSGAGGSAGSGAGGSASGSSPGNSADGG
GGSGGSGGSGGGSTQGGSMVSVIKPEMKIKLCMRGTVNGHNFVIEGEGKGNPYEGTQILDLNVTEGAPLP
FAYDILTTVFQYGNRAFTKYPADIQDYFKQTFPEGYHWERSMTYEDQGICTATSNISMRGDCFFYDIRFD
GTNFPPNGPVMQKKTLKWEPSTEKMYVEDGVLKGDVNMRLLLEGGGHYRCDFKTTYKAKKEVRLPDAHKI
DHRIEILKHDKDYNKVKLYENAVARYSMLPSQAKGTGTAAENSGNSRTKLMILWHEMWHEGLEEASRLYF
GERNVKGMFEVLEPLHAMMERGPQTLKETSFNQAYGRDLMEAQEWCRKYMKSGNVKDLLQAWDLYYHVFR
RISKPG*

>DNA sequence
ATGGCGTCGCTCACCGTGAAGGCCTACCTTCTGGGCAAGGAGGACGCGGCGCGCGAGATTCGCCGCTTCA
GCTTCTGTTGCAGCCCCGAGCCTGAGGCGGAAGCCGAGGCTGCGGCGGGTCCGGGACCCTGCGAGCGGCT
GCTGAGCCGGGTGGCCGCCCTGTTCCCCGCGCTGCGGCCTGGCGGCTTCCAGGCGCACTACCGCGCTGAG
CGCGGGGACTTGGTTGCCTTTTCCAGTGACGAGGAATTGACAATGGCCATGTCCTACGTGAAGGATGACA
TCTTCCGAATCTACATTAAAGAGAAAACCGGTTCTGGGAGTGGCGGTAGTGGTGCTGGAGGCAGCGCAGG
CTCTGGAGCAGGCGGCAGTGCCGGAAGTGGCGCTGGAGGGTCTGCCGGATCTGGAGCCGGTGGTTCTGCC
GGCTCCGGCGCTGGTGGGAGCGCTTCCGGAAGTAGTCCCGGTTCTGGGAGTGGCGGTAGTGGTGCTGGAG
GCAGCGCAGGCTCTGGAGCAGGCGGCAGTGCCGGAAGTGGCGCTGGAGGGTCTGCCGGATCTGGAGCCGG
TGGTTCTGCCGGCTCCGGCGCTGGTGGGAGCGCTTCCGGAAGTAGTCCCGGTAATTCCGCTGACGGCGGC
GGAGGATCGGGTGGTAGTGGTGGTTCAGGAGGAGGATCGACCCAAGGAGGATCCATGGTGAGTGTGATTA
AACCAGAGATGAAGATCAAGCTGTGTATGAGAGGCACTGTAAACGGGCATAATTTCGTGATTGAAGGAGA
AGGAAAAGGAAATCCTTACGAGGGAACGCAGATTTTAGACCTGAACGTCACTGAAGGCGCACCTCTGCCT
TTCGCTTACGATATCTTGACAACAGTGTTCCAGTACGGCAACAGGGCATTCACCAAGTACCCAGCAGATA
TTCAGGACTATTTCAAGCAGACTTTTCCTGAGGGGTATCACTGGGAAAGAAGCATGACTTATGAAGACCA
GGGCATTTGCACCGCCACAAGCAACATAAGCATGAGGGGCGACTGTTTTTTCTATGACATTCGTTTTGAT
GGCACCAACTTTCCTCCCAATGGTCCGGTTATGCAGAAGAAGACTCTTAAATGGGAGCCATCCACTGAGA
AAATGTACGTAGAGGATGGAGTGCTGAAGGGTGATGTTAACATGCGCCTGTTGCTTGAAGGAGGTGGCCA
TTATCGATGTGATTTCAAAACTACTTACAAAGCAAAGAAGGAGGTCCGTTTGCCAGACGCGCACAAAATT
GACCACCGCATTGAGATTTTGAAGCATGACAAAGATTACAACAAGGTCAAGCTCTATGAGAATGCCGTTG
CTCGCTATTCTATGCTGCCGAGTCAGGCCAAGGGTACCGGAACTGCAGCAGAGAATTCGGGAAACTCGAG
AACAAAGCTTATGATCCTCTGGCATGAGATGTGGCATGAAGGCCTGGAAGAGGCATCTCGTTTGTACTTT
GGGGAAAGGAACGTGAAAGGCATGTTTGAGGTGCTGGAGCCCTTGCATGCTATGATGGAACGGGGCCCCC
AGACTCTGAAGGAAACATCCTTTAATCAGGCCTATGGTCGAGATTTAATGGAGGCCCAAGAGTGGTGCAG
GAAGTACATGAAATCAGGGAATGTCAAGGACCTCCTCCAAGCCTGGGACCTCTATTATCATGTGTTCCGA
CGAATCTCAAAGCCCGGGTAA

**Annotation**
mPB1: monomeric PB1 domain (with D69A/D71R mutations)
mAG: monomeric Azami-Green
FRB: FRB domain from human mammalian target of rapamycin (mTOR) (residues 2021-2113 with a T2098L mutation)

**f**

**pCMV-FKBP-mCherry-Vav2cat**

>Amino acid sequence
MGVQVETISPGDGRTFPKRGQTCVVHYTGMLEDGKKFDSSRDRNKPFKFMLGKQEVIRGWEEGVAQMSVG
QRAKLTISPDYAYGATGHPGIIPPHATLVFDVELLKLEGSGASAPVATMVSKGEEDNMAIIKEFMRFKVH
MEGSVNGHEFEIEGEGEGRPYEGTQTAKLKVTKGGPLPFAWDILSPQFMYGSKAYVKHPADIPDYLKLSF
PEGFKWERVMNFEDGGVVTVTQDSSLQDGEFIYKVKLRGTNFPSDGPVMQKKTMGWEASSERMYPEDGAL
KGEIKQRLKLKDGGHYDAEVKTTYKAKKPVQLPGAYNVNIKLDITSHNEDYTIVEQYERAEGRHSTGGMD
ELYKSGLRSRGPMKMGMTEDDKRSCCLLEIQETEAKYYRTLEDIEKNYMGPLRLVLSPADMAAVFINLED
LIKVHHSFLRAIDVSMMAGGSTLAKVFLEFKERLLIYGEYCSHMEHAQSTLNQLLASREDFRQKVEECTL
RVQDGKFKLQDLLVVPMQRVLKYHLLLKELLSHSADRPERQQLKEALEAMQDLAMYINEVKRDKETLKKI
SEFQCSIENLQVKLEEFGRPKIDGELKVRSIVNHTKQDRYLFLFDKVVIVCKRKGYSYELKEVIELLFHK
MTDDPMHNKDIKKWSYGFYLIHLQGKQGFQFFCKTEDMKRKWMEQFEMAMSNIKPDKANANHHSFQMYTF
DKTTNCKACKMFLRGTFYQGYLCTRCGVGAHKECLEVIPPCK*

>DNA sequence
ATGGGAGTGCAGGTGGAAACCATCTCCCCAGGAGACGGGCGCACCTTCCCCAAGCGCGGCCAGACCTGCG
TGGTGCACTACACCGGGATGCTTGAAGATGGAAAGAAATTTGATTCCTCCCGGGACAGAAACAAGCCCTT
TAAGTTTATGCTAGGCAAGCAGGAGGTGATCCGAGGCTGGGAAGAAGGGGTTGCCCAGATGAGTGTGGGT
CAGAGAGCCAAACTGACTATATCTCCAGATTATGCCTATGGTGCCACTGGGCACCCAGGCATCATCCCAC
CACATGCCACTCTCGTCTTCGATGTGGAGCTTCTAAAACTGGAAGGCTCCGGTGCCAGTGCACCGGTCGC
CACCATGGTGAGCAAGGGCGAGGAGGATAACATGGCCATCATCAAGGAGTTCATGCGCTTCAAGGTGCAC
ATGGAGGGCTCCGTGAACGGCCACGAGTTCGAGATCGAGGGCGAGGGCGAGGGCCGCCCCTACGAGGGCA
CCCAGACCGCCAAGCTGAAGGTGACCAAGGGTGGCCCCCTGCCCTTCGCCTGGGACATCCTGTCCCCTCA
GTTCATGTACGGCTCCAAGGCCTACGTGAAGCACCCCGCCGACATCCCCGACTACTTGAAGCTGTCCTTC
CCCGAGGGCTTCAAGTGGGAGCGCGTGATGAACTTCGAGGACGGCGGCGTGGTGACCGTGACCCAGGACT
CCTCCCTGCAGGACGGCGAGTTCATCTACAAGGTGAAGCTGCGCGGCACCAACTTCCCCTCCGACGGCCC
CGTAATGCAGAAGAAGACCATGGGCTGGGAGGCCTCCTCCGAGCGGATGTACCCCGAGGACGGCGCCCTG
AAGGGCGAGATCAAGCAGAGGCTGAAGCTGAAGGACGGCGGCCACTACGACGCTGAGGTCAAGACCACCT
ACAAGGCCAAGAAGCCCGTGCAGCTGCCCGGCGCCTACAACGTCAACATCAAGTTGGACATCACCTCCCA
CAACGAGGACTACACCATCGTGGAACAGTACGAACGCGCCGAGGGCCGCCACTCCACCGGCGGCATGGAC
GAGCTGTACAAGTCCGGACTCAGATCTCGAGGGCCCATGAAAATGGGCATGACTGAGGACGACAAGAGAA
GCTGCTGCTTGTTAGAGATTCAGGAGACCGAGGCCAAGTACTACCGCACCCTGGAGGACATTGAGAAGAA
CTACATGGGTCCCTTGCGGCTGGTGCTGAGCCCGGCGGATATGGCTGCTGTCTTCATCAACCTGGAGGAC
CTCATCAAGGTGCATCACAGCTTTCTGCGAGCCATCGATGTGTCCATGATGGCTGGTGGCAGTACCCTGG
CTAAGGTCTTTCTGGAGTTTAAGGAAAGGCTCCTGATCTATGGAGAGTACTGTAGCCACATGGAACACGC
TCAGAGTACACTGAACCAGCTCCTCGCCAGCCGAGAGGACTTCAGGCAGAAAGTGGAGGAGTGCACACTC
AGGGTTCAGGATGGCAAGTTCAAGCTGCAAGACCTGCTGGTGGTGCCCATGCAACGGGTGCTGAAGTACC
ACCTGCTGCTCAAGGAGCTCCTGAGCCATTCTGCAGACCGACCAGAAAGACAACAGCTCAAAGAAGCCCT
GGAAGCCATGCAGGACTTGGCCATGTACATCAATGAAGTGAAGCGGGACAAGGAGACCTTGAAGAAGATT
AGCGAGTTCCAGTGCTCCATAGAAAACCTGCAAGTGAAGCTGGAGGAATTTGGGAGGCCAAAGATTGACG
GGGAGCTTAAAGTCCGGTCCATAGTCAACCACACCAAGCAAGACAGGTACCTGTTCCTATTTGACAAGGT
GGTCATCGTGTGCAAGAGGAAGGGCTACAGCTATGAGCTGAAGGAGGTCATTGAGCTGCTCTTCCACAAG
ATGACCGATGACCCGATGCACAACAAGGACATCAAGAAGTGGTCCTATGGCTTCTACCTGATTCACCTCC
AAGGAAAGCAAGGCTTTCAGTTCTTCTGCAAGACGGAAGACATGAAGCGGAAGTGGATGGAGCAGTTCGA
GATGGCCATGTCAAACATCAAGCCAGATAAGGCCAATGCCAACCATCATAGCTTCCAGATGTACACATTC
GACAAGACTACCAACTGCAAAGCCTGCAAGATGTTTCTCAGGGGTACCTTCTACCAGGGATACCTGTGTA
CCAGATGTGGCGTCGGGGCACACAAGGAATGCCTGGAGGTGATCCCCCCCTGCAAGTAA

**Annotation**
FKBP: FKBP12
mCherry: red fluorescent protein mCherry
$Vav2_{cat}$: DH-PH-CR domain from mouse Vav2 (residues 183-563)

**g**

**pPBbsr-Lifeact-mTagBFP2**

>Amino acid sequence
MGVADLIKKFESISKEEGDPPVATMVSKGEELIKENMHMKLYMEGTVDNHHFKCTSEGEGKPYEGTQTMR
IKVVEGGPLPFAFDILATSFLYGSKTFINHTQGIPDFFKQSFPEGFTWERVTTYEDGGVLTATQDTSLQD
GCLIYNVKIRGVNFTSNGPVMQKKTLGWEAFTETLYPADGGLEGRNDMALKLVGGSHLIANAKTTYRSKK
PAKNLKMPGVYYVDYRLERIKEANNETYVEQHEVAVARYCDLPSKLGHKLN*[EMCV_IRES]MLYEDNK
HHVGAAIRTKTGEIISAVHIEAYIGRVTVCAEAIAIGSAVSNGQKDFDTIVAVRHPYSDEVDRSIRVVSP
CGMCRELISDYAPDCFVLIEMNGKLVKTTIEELIPLKYTRN*

>DNA sequence
ATGGGCGTGGCCGACTTGATCAAGAAGTTCGAGTCCATCTCCAAGGAGGAGGGGGATCCACCGGTCGCCA
CCATGGTGTCTAAGGGCGAAGAGCTGATTAAGGAGAACATGCACATGAAGCTGTACATGGAGGGCACCGT
GGACAACCATCACTTCAAGTGCACATCCGAGGGCGAAGGCAAGCCCTACGAGGGCACCCAGACCATGAGA
ATCAAGGTGGTCGAGGGCGGCCCTCTCCCCTTCGCCTTCGACATCCTGGCTACTAGCTTCCTCTACGGCA
GCAAGACCTTCATCAACCACACCCAGGGCATCCCCGACTTCTTCAAGCAGTCCTTCCCTGAGGGCTTCAC
ATGGGAGAGAGTCACCACATACGAAGACGGGGGCGTGCTGACCGCTACCCAGGACACCAGCCTCCAGGAC
GGCTGCCTCATCTACAACGTCAAGATCAGAGGGGTGAACTTCACATCCAACGGCCCTGTGATGCAGAAGA
AAACACTCGGCTGGGAGGCCTTCACCGAGACGCTGTACCCCGCTGACGGCGGCCTGGAAGGCAGAAACGA
CATGGCCCTGAAGCTCGTGGGCGGGAGCCATCTGATCGCAAACGCCAAGACCACATATAGATCCAAGAAA
CCCGCTAAGAACCTCAAGATGCCTGGCGTCTACTATGTGGACTACAGACTGGAAAGAATCAAGGAGGCCA
ACAACGAGACCTACGTCGAGCAGCACGAGGTGGCAGTGGCCAGATACTGCGACCTCCCTAGCAAACTGGG
GCACAAGCTTAATTAAGCGGCCGCTCTAGAGTCGACGGGCCGCGGTAACAATTGTTAACTAACTTAAGCT
AGCAACGGTTTCCCTCTAGCGGGATCAATTCCGCCCCCCCCCCCTAACGTTACTGGCCGAAGCCGCTTGG
AATAAGGCCGGTGTGCGTTTGTCTATATGTTATTTTCCACCATATTGCCGTCTTTTGGCAATGTGAGGGC
CCGGAAACCTGGCCCTGTCTTCTTGACGAGCATTCCTAGGGGTCTTTCCCCTCTCGCCAAAGGAATGCAA
GGTCTGTTGAATGTCGTGAAGGAAGCAGTTCCTCTGGAAGCTTCTTGAAGACAAACAACGTCTGTAGCGA
CCCTTTGCAGGCAGCGGAACCCCCCACCTGGCGACAGGTGCCTCTGCGGCCAAAAGCCACGTGTATAAGA
TACACCTGCAAAGGCGGCACAACCCCAGTGCCACGTTGTGAGTTGGATAGTTGTGGAAAGAGTCAAATGG
CTCTCCTCAAGCGTATTCAACAAGGGGCTGAAGGATGCCCAGAAGGTACCCCATTGTATGGGATCTGATC
TGGGGCCTCGGTGCACATGCTTTACATGTGTTTAGTCGAGGTTAAAAAACGTCTAGGCCCCCCGAACCAC
GGGGACGTGGTTTTCCTTTGAAAAACACGATAATACCATGGTCATGAAAACATTTAACATTTCTCAACAA
GATCTAGAATTAGTAGAAGTAGCGACAGAGAAGATTACAATGCTTTATGAGGATAATAAACATCATGTGG
GAGCGGCAATTCGTACGAAAACAGGAGAAATCATTTCGGCAGTACATATTGAAGCGTATATAGGACGAGT
AACTGTTTGTGCAGAAGCCATTGCGATTGGTAGTGCAGTTTCGAATGGACAAAAGGATTTTGACACGATT
GTAGCTGTTAGACACCCTTATTCTGACGAAGTAGATAGAAGTATTCGAGTGGTAAGTCCTTGTGGTATGT
GTAGGGAGTTGATTTCAGACTATGCACCAGATTGTTTTGTGTTAATAGAAATGAATGGCAAGTTAGTCAA
AACTACGATTGAAGAACTCATTCCACTCAAATATACCCGAAATTAA

**Annotation**
Lifeact: actin-binding peptide Lifeact
mTagBFP2: blue fluorescent protein mTagBFP2
EMCV IRES: internal ribosomal entry site from *Encephalomyocarditis* virus
Bsr: blasticidin S-deaminase

**h**

**pCMV-PakGBD-tdiRFP670**

>Amino acid sequence
MKKEKERPEISLPSDFEHTIHVGFDAVTGEFTGMPEQWARLLQTSNITKSEQKKNPQAVLDVLEFYNSKK
TSNSQKYMSFTDKSPPVATMARKVDLTSCDREPIHIPGSIQPCGCLLACDAQAVRITRITENAGAFFGRE
TPRVGELLADYFGETEAHALRNALAQSSDPKRPALIFGWRDGLTGRTFDISLHRHDGTSIIEFEPAAAEQ
ADNPLRLTRQIIARTKELKSLEEMAARVPRYLQAMLGYHRVMLYRFADDGSGMVIGEAKRSDLESFLGQH
FPASLVPQQARLLYLKNAIRVVSDSRGISSRIVPEHDASGAALDLSFAHLRSISPCHLEFLRNMGVSASM
SLSIIIDGTLWGLIICHHYEPRAVPMAQRVAAEMFADFLSLHFTAAHHQRGHATGSTGSGSAEGGTASSE
DNMARKVDLTSCDREPIHIPGSIQPCGCLLACDAQAVRITRITENAGAFFGRETPRVGELLADYFGETEA
HALRNALAQSSDPKRPALIFGWRDGLTGRTFDISLHRHDGTSIIEFEPAAAEQADNPLRLTRQIIARTKE
LKSLEEMAARVPRYLQAMLGYHRVMLYRFADDGSGMVIGEAKRSDLESFLGQHFPASLVPQQARLLYLKN
AIRVVSDSRGISSRIVPEHDASGAALDLSFAHLRSISPCHLEFLRNMGVSASMSLSIIIDGTLWGLIICH
HYEPRAVPMAQRVAAEMFADFLSLHFTAAHHQRMYK*

>DNA sequence
ATGAAGAAAGAGAAAGAGCGGCCAGAGATTTCTCTCCCTTCAGATTTTGAACACACAATTCATGTCGGTT
TTGATGCTGTCACAGGGGAGTTTACGGGAATGCCAGAGCAGTGGGCCCGCTTGCTTCAGACATCAAATAT
CACTAAGTCGGAGCAGAAGAAAAACCCGCAGGCTGTTCTGGATGTGTTGGAGTTTTACAACTCGAAGAAG
ACATCCAACAGCCAGAAATACATGAGCTTTACAGATAAGTCACCACCGGTCGCCACCATGGCTCGCAAGG
TGGACCTGACCAGCTGCGACCGCGAGCCCATCCACATCCCCGGCAGCATCCAGCCCTGCGGCTGCCTGCT
GGCCTGCGACGCCCAGGCCGTGCGCATCACCCGCATCACCGAGAACGCCGGCGCCTTCTTCGGCCGCGAG
ACCCCCCGCGTGGGCGAGCTGCTGGCCGACTACTTCGGCGAGACCGAGGCCCACGCCCTGCGCAACGCCC
TGGCCCAGAGCAGCGACCCCAAGCGCCCCGCCCTGATCTTCGGCTGGCGCGACGGCCTGACCGGCCGCAC
CTTCGACATCAGCCTGCACCGCCACGACGGCACCAGCATCATCGAGTTCGAGCCCGCCGCCGCCGAGCAG
GCCGACAACCCCCTGCGCCTGACCCGCCAGATCATCGCCCGCACCAAGGAGCTGAAGAGCCTGGAGGAGA
TGGCCGCCCGCGTGCCCCGCTACCTGCAGGCCATGCTGGGCTACCACCGCGTGATGCTGTACCGCTTCGC
CGACGACGGCAGCGGCATGGTGATCGGCGAGGCCAAGCGCAGCGACCTGGAGAGCTTCCTGGGCCAGCAC
TTCCCCGCCAGCCTGGTGCCCCAGCAGGCCCGCCTGCTGTACCTGAAGAACGCCATCCGCGTGGTGAGCG
ACAGCCGCGGCATCAGCAGCCGCATCGTGCCCGAGCACGACGCCAGCGGCGCCGCCCTGGACCTGAGCTT
CGCCCACCTGCGCAGCATCAGCCCCTGCCACCTGGAGTTCCTGCGCAACATGGGCGTGAGCGCCAGCATG
AGCCTGAGCATCATCATCGACGGCACCCTGTGGGGCCTGATCATCTGCCACCACTACGAGCCCCGCGCCG
TGCCCATGGCCCAGCGCGTGGCCGCCGAGATGTTCGCCGACTTCCTGAGCCTGCACTTCACCGCCGCTCA
CCATCAGAGAGGACATGCTACTGGAAGCACTGGAAGCGGCAGCGCTGAGGGAGGCACAGCTTCTAGCGAA
GATAATATGGCTCGCAAGGTGGACCTGACCAGCTGCGACCGCGAGCCCATCCACATCCCCGGCAGCATCC
AGCCCTGCGGCTGCCTGCTGGCCTGCGACGCCCAGGCCGTGCGCATCACCCGCATCACCGAGAACGCCGG
CGCCTTCTTCGGCCGCGAGACCCCCCGCGTGGGCGAGCTGCTGGCCGACTACTTCGGCGAGACCGAGGCC
CACGCCCTGCGCAACGCCCTGGCCCAGAGCAGCGACCCCAAGCGCCCCGCCCTGATCTTCGGCTGGCGCG
ACGGCCTGACCGGCCGCACCTTCGACATCAGCCTGCACCGCCACGACGGCACCAGCATCATCGAGTTCGA
GCCCGCCGCCGCCGAGCAGGCCGACAACCCCCTGCGCCTGACCCGCCAGATCATCGCCCGCACCAAGGAG
CTGAAGAGCCTGGAGGAGATGGCCGCCCGCGTGCCCCGCTACCTGCAGGCCATGCTGGGCTACCACCGCG
TGATGCTGTACCGCTTCGCCGACGACGGCAGCGGCATGGTGATCGGCGAGGCCAAGCGCAGCGACCTGGA
GAGCTTCCTGGGCCAGCACTTCCCCGCCAGCCTGGTGCCCCAGCAGGCCCGCCTGCTGTACCTGAAGAAC
GCCATCCGCGTGGTGAGCGACAGCCGCGGCATCAGCAGCCGCATCGTGCCCGAGCACGACGCCAGCGGCG
CCGCCCTGGACCTGAGCTTCGCCCACCTGCGCAGCATCAGCCCCTGCCACCTGGAGTTCCTGCGCAACAT
GGGCGTGAGCGCCAGCATGAGCCTGAGCATCATCATCGACGGCACCCTGTGGGGCCTGATCATCTGCCAC
CACTACGAGCCCCGCGCCGTGCCCATGGCCCAGCGCGTGGCCGCCGAGATGTTCGCCGACTTCCTGAGCC
TGCACTTCACCGCCGCTCACCATCAGAGAATGTACAAGTAA

**Annotation**
PakGBD: Rac(GTP)-binding domain from Pak
iRFP670: near-infrared fluorescent protein iRFP670

**i**

**pCMV-mCherry-Vav2cat**

>Amino acid sequence
MVSKGEEDNMAIIKEFMRFKVHMEGSVNGHEFEIEGEGEGRPYEGTQTAKLKVTKGGPLPFAWDILSPQF
MYGSKAYVKHPADIPDYLKLSFPEGFKWERVMNFEDGGVVTVTQDSSLQDGEFIYKVKLRGTNFPSDGPV
MQKKTMGWEASSERMYPEDGALKGEIKQRLKLKDGGHYDAEVKTTYKAKKPVQLPGAYNVNIKLDITSHN
EDYTIVEQYERAEGRHSTGGMDELYKSGLRSRGPMKMGMTEDDKRSCCLLEIQETEAKYYRTLEDIEKNY
MGPLRLVLSPADMAAVFINLEDLIKVHHSFLRAIDVSMMAGGSTLAKVFLEFKERLLIYGEYCSHMEHAQ
STLNQLLASREDFRQKVEECTLRVQDGKFKLQDLLVVPMQRVLKYHLLLKELLSHSADRPERQQLKEALE
AMQDLAMYINEVKRDKETLKKISEFQCSIENLQVKLEEFGRPKIDGELKVRSIVNHTKQDRYLFLFDKVV
IVCKRKGYSYELKEVIELLFHKMTDDPMHNKDIKKWSYGFYLIHLQGKQGFQFFCKTEDMKRKWMEQFEM
AMSNIKPDKANANHHSFQMYTFDKTTNCKACKMFLRGTFYQGYLCTRCGVGAHKECLEVIPPCK*

>DNA sequence
ATGGTGAGCAAGGGCGAGGAGGATAACATGGCCATCATCAAGGAGTTCATGCGCTTCAAGGTGCACATGG
AGGGCTCCGTGAACGGCCACGAGTTCGAGATCGAGGGCGAGGGCGAGGGCCGCCCCTACGAGGGCACCCA
GACCGCCAAGCTGAAGGTGACCAAGGGTGGCCCCCTGCCCTTCGCCTGGGACATCCTGTCCCCTCAGTTC
ATGTACGGCTCCAAGGCCTACGTGAAGCACCCCGCCGACATCCCCGACTACTTGAAGCTGTCCTTCCCCG
AGGGCTTCAAGTGGGAGCGCGTGATGAACTTCGAGGACGGCGGCGTGGTGACCGTGACCCAGGACTCCTC
CCTGCAGGACGGCGAGTTCATCTACAAGGTGAAGCTGCGCGGCACCAACTTCCCCTCCGACGGCCCCGTA
ATGCAGAAGAAGACCATGGGCTGGGAGGCCTCCTCCGAGCGGATGTACCCCGAGGACGGCGCCCTGAAGG
GCGAGATCAAGCAGAGGCTGAAGCTGAAGGACGGCGGCCACTACGACGCTGAGGTCAAGACCACCTACAA
GGCCAAGAAGCCCGTGCAGCTGCCCGGCGCCTACAACGTCAACATCAAGTTGGACATCACCTCCCACAAC
GAGGACTACACCATCGTGGAACAGTACGAACGCGCCGAGGGCCGCCACTCCACCGGCGGCATGGACGAGC
TGTACAAGTCCGGACTCAGATCTCGAGGGCCCATGAAAATGGGCATGACTGAGGACGACAAGAGAAGCTG
CTGCTTGTTAGAGATTCAGGAGACCGAGGCCAAGTACTACCGCACCCTGGAGGACATTGAGAAGAACTAC
ATGGGTCCCTTGCGGCTGGTGCTGAGCCCGGCGGATATGGCTGCTGTCTTCATCAACCTGGAGGACCTCA
TCAAGGTGCATCACAGCTTTCTGCGAGCCATCGATGTGTCCATGATGGCTGGTGGCAGTACCCTGGCTAA
GGTCTTTCTGGAGTTTAAGGAAAGGCTCCTGATCTATGGAGAGTACTGTAGCCACATGGAACACGCTCAG
AGTACACTGAACCAGCTCCTCGCCAGCCGAGAGGACTTCAGGCAGAAAGTGGAGGAGTGCACACTCAGGG
TTCAGGATGGCAAGTTCAAGCTGCAAGACCTGCTGGTGGTGCCCATGCAACGGGTGCTGAAGTACCACCT
GCTGCTCAAGGAGCTCCTGAGCCATTCTGCAGACCGACCAGAAAGACAACAGCTCAAAGAAGCCCTGGAA
GCCATGCAGGACTTGGCCATGTACATCAATGAAGTGAAGCGGGACAAGGAGACCTTGAAGAAGATTAGCG
AGTTCCAGTGCTCCATAGAAAACCTGCAAGTGAAGCTGGAGGAATTTGGGAGGCCAAAGATTGACGGGGA
GCTTAAAGTCCGGTCCATAGTCAACCACACCAAGCAAGACAGGTACCTGTTCCTATTTGACAAGGTGGTC
ATCGTGTGCAAGAGGAAGGGCTACAGCTATGAGCTGAAGGAGGTCATTGAGCTGCTCTTCCACAAGATGA
CCGATGACCCGATGCACAACAAGGACATCAAGAAGTGGTCCTATGGCTTCTACCTGATTCACCTCCAAGG
AAAGCAAGGCTTTCAGTTCTTCTGCAAGACGGAAGACATGAAGCGGAAGTGGATGGAGCAGTTCGAGATG
GCCATGTCAAACATCAAGCCAGATAAGGCCAATGCCAACCATCATAGCTTCCAGATGTACACATTCGACA
AGACTACCAACTGCAAAGCCTGCAAGATGTTTCTCAGGGGTACCTTCTACCAGGGATACCTGTGTACCAG
ATGTGGCGTCGGGGCACACAAGGAATGCCTGGAGGTGATCCCCCCCTGCAAGTAA

**Annotation**
mCherry: red fluorescent protein mCherry
$Vav2_{cat}$: DH-PH-CR domain from mouse Vav2 (residues 183-563)

**j**

**pCMV-FKBP-mTagBFP2**

>Amino acid sequence
MGVQVETISPGDGRTFPKRGQTCVVHYTGMLEDGKKFDSSRDRNKPFKFMLGKQEVIRGWEEGVAQMSVG
QRAKLTISPDYAYGATGHPGIIPPHATLVFDVELLKLEGSGASAPVATMVSKGEELIKENMHMKLYMEGT
VDNHHFKCTSEGEGKPYEGTQTMRIKVVEGGPLPFAFDILATSFLYGSKTFINHTQGIPDFFKQSFPEGF
TWERVTTYEDGGVLTATQDTSLQDGCLIYNVKIRGVNFTSNGPVMQKKTLGWEAFTETLYPADGGLEGRN
DMALKLVGGSHLIANAKTTYRSKKPAKNLKMPGVYYVDYRLERIKEANNETYVEQHEVAVARYCDLPSKL
GHKLNRSRAQASNSAVDGTAGPGSTGSR*

>DNA sequence
ATGGGAGTGCAGGTGGAAACCATCTCCCCAGGAGACGGGCGCACCTTCCCCAAGCGCGGCCAGACCTGCG
TGGTGCACTACACCGGGATGCTTGAAGATGGAAAGAAATTTGATTCCTCCCGGGACAGAAACAAGCCCTT
TAAGTTTATGCTAGGCAAGCAGGAGGTGATCCGAGGCTGGGAAGAAGGGGTTGCCCAGATGAGTGTGGGT
CAGAGAGCCAAACTGACTATATCTCCAGATTATGCCTATGGTGCCACTGGGCACCCAGGCATCATCCCAC
CACATGCCACTCTCGTCTTCGATGTGGAGCTTCTAAAACTGGAAGGCTCCGGTGCCAGTGCACCGGTCGC
CACCATGGTGTCTAAGGGCGAAGAGCTGATTAAGGAGAACATGCACATGAAGCTGTACATGGAGGGCACC
GTGGACAACCATCACTTCAAGTGCACATCCGAGGGCGAAGGCAAGCCCTACGAGGGCACCCAGACCATGA
GAATCAAGGTGGTCGAGGGCGGCCCTCTCCCCTTCGCCTTCGACATCCTGGCTACTAGCTTCCTCTACGG
CAGCAAGACCTTCATCAACCACACCCAGGGCATCCCCGACTTCTTCAAGCAGTCCTTCCCTGAGGGCTTC
ACATGGGAGAGAGTCACCACATACGAAGACGGGGGCGTGCTGACCGCTACCCAGGACACCAGCCTCCAGG
ACGGCTGCCTCATCTACAACGTCAAGATCAGAGGGGTGAACTTCACATCCAACGGCCCTGTGATGCAGAA
GAAAACACTCGGCTGGGAGGCCTTCACCGAGACGCTGTACCCCGCTGACGGCGGCCTGGAAGGCAGAAAC
GACATGGCCCTGAAGCTCGTGGGCGGGAGCCATCTGATCGCAAACGCCAAGACCACATATAGATCCAAGA
AACCCGCTAAGAACCTCAAGATGCCTGGCGTCTACTATGTGGACTACAGACTGGAAAGAATCAAGGAGGC
CAACAACGAGACCTACGTCGAGCAGCACGAGGTGGCAGTGGCCAGATACTGCGACCTCCCTAGCAAACTG
GGGCACAAGCTTAATAGATCTCGAGCTCAAGCTTCGAATTCTGCAGTCGACGGTACCGCGGGCCCGGGAT
CCACCGGATCTAGATAA

**Annotation**
FKBP: FKBP12
mTagBFP2: blue fluorescent protein mTagBFP2

**k**

**pCMV-FKBP-mTagBFP2-SOS1cat**

>Amino acid sequence
MGVQVETISPGDGRTFPKRGQTCVVHYTGMLEDGKKFDSSRDRNKPFKFMLGKQEVIRGWEEGVAQMSVG
QRAKLTISPDYAYGATGHPGIIPPHATLVFDVELLKLEGSGASAPVATMVSKGEELIKENMHMKLYMEGT
VDNHHFKCTSEGEGKPYEGTQTMRIKVVEGGPLPFAFDILATSFLYGSKTFINHTQGIPDFFKQSFPEGF
TWERVTTYEDGGVLTATQDTSLQDGCLIYNVKIRGVNFTSNGPVMQKKTLGWEAFTETLYPADGGLEGRN
DMALKLVGGSHLIANAKTTYRSKKPAKNLKMPGVYYVDYRLERIKEANNETYVEQHEVAVARYCDLPSKL
GHKLNRSRAQASQMRLPSADVYRFAEPDSEENIIFEENMQPKAGIPIIKAGTVIKLIERLTYHMYADPNF
VRTFLTTYRSFCKPQELLSLIIERFEIPEPEPTEADRIAIENGDQPLSAELKRFRKEYIQPVQLRVLNVC
RHWVEHHFYDFERDAYLLQRMEEFIGTVRGKAMKKWVESITKIIQRKKIARDNGPGHNITFQSSPPTVEW
HISRPGHIETFDLLTLHPIEIARQLTLLESDLYRAVQPSELVGSVWTKEDKEINSPNLLKMIRHTTNLTL
WFEKCIVETENLEERVAVVSRIIEILQVFQELNNFNGVLEVVSAMNSSPVYRLDHTFEQIPSRQKKILEE
AHELSEDHYKKYLAKLRSINPPCVPFFGIYLTNILKTEEGNPEVLKRHGKELINFSKRRKVAEITGEIQQ
YQNQPYCLRVESDIKRFFENLNPMGNSMEKEFTDYLFNKSLEIEPRNPKPLPRFPKKYSYPLKSPGVRPS
NPRPGT*

>DNA sequence
ATGGGAGTGCAGGTGGAAACCATCTCCCCAGGAGACGGGCGCACCTTCCCCAAGCGCGGCCAGACCTGCG
TGGTGCACTACACCGGGATGCTTGAAGATGGAAAGAAATTTGATTCCTCCCGGGACAGAAACAAGCCCTT
TAAGTTTATGCTAGGCAAGCAGGAGGTGATCCGAGGCTGGGAAGAAGGGGTTGCCCAGATGAGTGTGGGT
CAGAGAGCCAAACTGACTATATCTCCAGATTATGCCTATGGTGCCACTGGGCACCCAGGCATCATCCCAC
CACATGCCACTCTCGTCTTCGATGTGGAGCTTCTAAAACTGGAAGGCTCCGGTGCCAGTGCACCGGTCGC
CACCATGGTGTCTAAGGGCGAAGAGCTGATTAAGGAGAACATGCACATGAAGCTGTACATGGAGGGCACC
GTGGACAACCATCACTTCAAGTGCACATCCGAGGGCGAAGGCAAGCCCTACGAGGGCACCCAGACCATGA
GAATCAAGGTGGTCGAGGGCGGCCCTCTCCCCTTCGCCTTCGACATCCTGGCTACTAGCTTCCTCTACGG
CAGCAAGACCTTCATCAACCACACCCAGGGCATCCCCGACTTCTTCAAGCAGTCCTTCCCTGAGGGCTTC
ACATGGGAGAGAGTCACCACATACGAAGACGGGGGCGTGCTGACCGCTACCCAGGACACCAGCCTCCAGG
ACGGCTGCCTCATCTACAACGTCAAGATCAGAGGGGTGAACTTCACATCCAACGGCCCTGTGATGCAGAA
GAAAACACTCGGCTGGGAGGCCTTCACCGAGACGCTGTACCCCGCTGACGGCGGCCTGGAAGGCAGAAAC
GACATGGCCCTGAAGCTCGTGGGCGGGAGCCATCTGATCGCAAACGCCAAGACCACATATAGATCCAAGA
AACCCGCTAAGAACCTCAAGATGCCTGGCGTCTACTATGTGGACTACAGACTGGAAAGAATCAAGGAGGC
CAACAACGAGACCTACGTCGAGCAGCACGAGGTGGCAGTGGCCAGATACTGCGACCTCCCTAGCAAACTG
GGGCACAAGCTTAATAGATCTCGAGCTCAAGCTTCGCAGATGAGGCTGCCTAGTGCTGATGTTTATAGAT
TTGCAGAGCCTGACTCTGAAGAGAATATTATATTTGAAGAGAACATGCAGCCCAAGGCTGGAATTCCAAT
TATCAAAGCAGGAACTGTTATTAAACTTATAGAGAGGCTTACGTACCATATGTACGCAGATCCCAATTTT
GTTCGGACATTTCTTACAACATACAGATCCTTTTGCAAACCTCAAGAACTACTGAGTCTTATAATAGAAA
GGTTTGAAATTCCAGAGCCTGAGCCAACAGAAGCTGATCGCATAGCTATAGAGAATGGAGATCAACCCTT
GAGTGCAGAACTGAAAAGATTTAGAAAAGAATATATACAGCCTGTGCAACTGCGAGTATTAAATGTATGT
CGGCACTGGGTAGAGCACCACTTCTATGATTTTGAAAGAGATGCATATCTTTTGCAACGAATGGAAGAAT
TTATTGGAACAGTAAGAGGTAAAGCAATGAAAAAATGGGTTGAATCCATCACTAAAATAATCCAAAGGAA
AAAAATTGCAAGAGACAATGGACCAGGTCATAATATTACATTTCAGAGTTCACCTCCCACAGTTGAGTGG
CATATAAGCAGACCTGGGCACATAGAGACTTTTGACCTGCTCACCTTACACCCAATAGAAATTGCTCGAC
AACTCACTTTACTTGAATCAGATCTATACCGAGCTGTACAGCCATCAGAATTAGTTGGAAGTGTGTGGAC
AAAAGAAGACAAAGAAATTAACTCTCCTAATCTTCTGAAAATGATTCGACATACCACCAACCTCACTCTG
TGGTTTGAGAAATGTATTGTAGAAACTGAAAATTTAGAAGAAAGAGTAGCTGTGGTGAGTCGAATTATTG
AGATTCTACAAGTCTTTCAAGAGTTGAACAACTTTAATGGTGTCCTTGAGGTTGTCAGTGCTATGAATTC
ATCACCTGTTTACAGACTAGACCACACATTTGAGCAAATACCAAGTCGCCAGAAGAAAATTTTAGAAGAA
GCTCATGAATTGAGTGAAGATCACTATAAGAAATATTTGGCAAAACTCAGGTCTATTAATCCACCATGTG
TGCCTTTCTTTGGAATTTATCTCACTAATATCTTGAAAACAGAAGAAGGCAACCCTGAGGTCCTAAAAAG
ACATGGAAAAGAGCTTATAAACTTTAGCAAAAGGAGGAAAGTAGCAGAAATAACAGGAGAGATCCAGCAG
TACCAAAATCAGCCTTACTGTTTACGAGTAGAATCAGATATCAAAAGGTTCTTTGAAAACTTGAATCCGA
TGGGAAATAGCATGGAGAAGGAATTTACAGATTATCTTTTCAACAAATCCCTAGAAATAGAACCACGAAA

CCCTAAGCCTCTCCCAAGATTTCCAAAAAAATATAGCTATCCCCTAAAATCTCCTGGTGTTCGTCCATCA
AACCCAAGACCAGGTACCTAA

**Annotation**

FKBP: FKBP12

mTagBFP2: blue fluorescent protein mTagBFP2

$SOS1_{cat}$: Rem-Cdc25 (RasGEF) domain from human SOS1 (residues 566-1049)

**pPBbsr-MEK1-P2A-mCherry-ERK2**

>Amino acid sequence
MDYKDDDDKARLEMPKKKPTPIQLNPNPEGTAVNGTPTAETNLEALQKKLEELELDEQQRKRLEAFLTQK
QKVGELKDDDFEKVSELGAGNGGVVFKVSHKPTSLIMARKLIHLEIKPAIRNQIIRELQVLHECNSPYIV
GFYGAFYSDGEISICMEHMDGGSLDQVLKKAGKIPEKILGKVSIAVIKGLTYLREKHKIMHRDVKPSNIL
VNSRGEIKLCDFGVSGQLIDSMANSFVGTRSYMSPERLQGTHYSVQSDIWSMGLSLVEMAIGRYPIPPPD
AKELELIFGCSVERDPASSELAPRPRPPGRPISSYGPDSRPPMAIFELLDYIVNEPPPKLPSGVFGAEFQ
DFVNKCLVKNPAERADLKQLMVHSFIKQSELEEVDFAGWLCSTMGLKQPSTPTHAAGVGGRGTSGSGATN
FSLLKQAGDVEENPGPQLIKGAMVSKGEEDNMAIIKEFMRFKVHMEGSVNGHEFEIEGEGEGRPYEGTQT
AKLKVTKGGPLPFAWDILSPQFMYGSKAYVKHPADIPDYLKLSFPEGFKWERVMNFEDGGVVTVTQDSSL
QDGEFIYKVKLRGTNFPSDGPVMQKKTMGWEASSERMYPEDGALKGEIKQRLKLKDGGHYDAEVKTTYKA
KKPVQLPGAYNVNIKLDITSHNEDYTIVEQYDRAEGRHSTGGMDELYLEMAAAGAASNPGGGPEMVRGQA
FDVGPRYINLAYIGEGAYGMVCSAHDNVNKVRVAIKKISPFEHQTYCQRTLREIKILLRFKHENIIGIND
IIRAPTIEQMKDVYIVQDLMETDLYKLLKTQHLSNDHICYFLYQILRGLKYIHSANVLHRDLKPSNLLLN
TTCDLKICDFGLARVADPDHDHTGFLTEYVATRWYRAPEIMLNSKGYTKSIDIWSVGCILAEMLSNRPIF
PGKHYLDQLNHILGILGSPSQEDLNCIINLKARNYLLSLPHKNKVPWNRLFPNADPKALDLLDKMLTFNP
HKRIEVEAALAHPYLEQYYDPSDEPVAEAPFKFEMELDDLPKETLKELIFEETARFQPGY*[EMCV_IRE
S]MLYEDNKHHVGAAIRTKTGEIISAVHIEAYIGRVTVCAEAIAIGSAVSNGQKDFDTIVAVRHPYSDEV
DRSIRVVSPCGMCRELISDYAPDCFVLIEMNGKLVKTTIEELIPLKYTRN*

>DNA sequence
ATGGACTACAAAGACGATGACGATAAAGCAAGGCTCGAGATGCCTAAAAAGAAGCCTACGCCCATACAGC
TGAATCCCAACCCCGAAGGGACTGCTGTGAACGGGACCCCTACAGCCGAGACAAACCTTGAAGCTCTGCA
GAAAAAGTTGGAAGAGCTTGAGCTGGATGAGCAGCAGAGGAAGCGTCTGGAGGCTTTTCTCACCCAGAAG
CAGAAAGTTGGGGAACTGAAGGATGACGACTTTGAAAAAGTTTCAGAGCTTGGAGCAGGCAACGGAGGAG
TGGTGTTTAAGGTGTCCCACAAGCCAACCAGCTTGATTATGGCCAGGAAGTTGATTCATCTGGAGATTAA
GCCTGCAATCCGAAACCAGATTATCCGAGAGTTGCAGGTTCTGCATGAATGTAACTCCCCATACATTGTG
GGGTTCTATGGGGCCTTCTACAGTGATGGAGAGATCAGCATTTGCATGGAACACATGGATGGAGGCTCCC
TTGATCAGGTTCTGAAGAAAGCTGGCAAAATCCCAGAAAAGATTTTGGGAAAAGTCAGCATTGCAGTGAT
AAAAGGTCTAACCTACCTGAGAGAAAAGCATAAGATAATGCACAGAGATGTGAAACCTTCTAACATCCTG
GTCAACTCTAGAGGAGAGATAAAACTCTGCGACTTTGGGGTCAGCGGGCAACTCATAGACTCCATGGCAA
ATTCCTTTGTTGGGACAAGATCCTATATGTCACCGGAGCGACTACAGGGCACTCATTATTCTGTGCAATC
AGACATCTGGAGCATGGGGCTGTCGCTGGTGGAAATGGCCATTGGAAGGTATCCCATTCCACCCCCTGAT
GCCAAAGAGCTGGAACTTATCTTTGGGTGTTCTGTAGAAAGGGATCCAGCGTCTTCTGAACTGGCACCTC
GCCCCCGGCCACCCGGACGTCCAATAAGCTCATACGGTCCTGATAGTCGACCACCCATGGCTATTTTTGA
ACTTCTGGATTATATCGTGAACGAGCCGCCTCCAAAATTGCCCAGTGGAGTATTTGGAGCTGAGTTCCAG
GACTTTGTGAATAAATGTCTTGTGAAGAATCCGGCAGAGAGAGCAGACCTTAAACAGCTAATGGTTCACA
GCTTCATTAAGCAGTCAGAGTTGGAGGAAGTGGATTTTGCTGGATGGCTCTGTTCCACTATGGGCCTTAA
GCAGCCCAGTACCCCAACCCATGCCGCCGGAGTGGGCGGCCGCGGCACTAGTGGAAGCGGAGCTACTAAC
TTCAGCCTGCTGAAGCAGGCTGGAGACGTGGAGGAGAACCCTGGACCTCAATTAATTAAGGGCGCAATGG
TGAGCAAGGGCGAGGAGGATAACATGGCCATCATCAAGGAGTTCATGCGCTTCAAGGTGCACATGGAGGG
CTCCGTGAACGGCCACGAGTTCGAGATCGAGGGCGAGGGCGAGGGCCGCCCCTACGAGGGCACCCAGACC
GCCAAGCTGAAGGTGACCAAGGGTGGCCCCCTGCCCTTCGCCTGGGACATCCTGTCCCCTCAGTTCATGT
ACGGCTCCAAGGCCTACGTGAAGCACCCCGCCGACATCCCCGACTACTTGAAGCTGTCCTTCCCCGAGGG
CTTCAAGTGGGAGCGCGTGATGAACTTCGAGGACGGCGGCGTGGTGACCGTGACCCAGGACTCCTCCCTG
CAGGACGGCGAGTTCATCTACAAGGTGAAGCTGCGCGGCACCAACTTCCCCTCCGACGGCCCCGTAATGC
AGAAGAAGACCATGGGCTGGGAGGCCTCCTCCGAGCGGATGTACCCCGAGGACGGCGCCCTGAAGGGCGA
GATCAAGCAGAGGCTGAAGCTGAAGGACGGCGGCCACTACGACGCTGAGGTCAAGACCACCTACAAGGCC
AAGAAGCCCGTGCAGCTGCCCGGCGCCTACAACGTCAACATCAAGTTGGACATCACCTCCCACAACGAGG
ACTACACCATCGTGGAACAGTACGACCGCGCCGAGGGCCGCCACTCCACCGGCGGCATGGACGAGCTGTA
CCTCGAGATGGCAGCGGCAGGAGCTGCGTCTAACCCCGGCGGGGGTCCGGAGATGGTGCGGGGCCAGGCG
TTCGACGTAGGCCCTCGATACATCAATCTGGCTTATATCGGCGAGGGAGCGTACGGCATGGTGTGTTCTG

```
CCCATGACAATGTTAACAAAGTTCGAGTTGCTATCAAGAAAATCAGCCCATTTGAGCATCAGACATACTG
CCAGCGAACATTGCGGGAGATCAAAATCTTGCTACGTTTTAAACATGAAAACATCATTGGGATAAACGAC
ATTATTCGCGCTCCAACCATTGAGCAGATGAAAGATGTGTACATTGTGCAGGACCTCATGGAGACAGACC
TCTATAAGCTCCTGAAGACTCAGCATCTTAGCAATGACCATATCTGCTATTTCTTGTACCAGATTCTGAG
AGGATTAAAGTACATCCATTCAGCCAATGTTCTACATCGTGATCTTAAGCCTTCAAATTTGCTGCTTAAC
ACTACCTGTGATCTCAAGATCTGTGATTTTGGATTGGCTCGTGTTGCAGACCCAGATCATGATCACACTG
GCTTTCTCACAGAATATGTAGCCACTCGCTGGTACAGAGCTCCTGAGATCATGCTGAATTCCAAGGGCTA
TACCAAATCAATTGACATCTGGTCTGTTGGCTGCATTCTTGCTGAGATGCTTTCTAATAGACCCATATTT
CCTGGGAAACATTATCTTGACCAGCTTAATCACATACTTGGTATTCTTGGATCTCCATCTCAAGAGGACC
TAAACTGTATAATCAATTTAAAAGCTAGGAATTACTTGCTTTCCCTTCCTCACAAAAATAAGGTGCCATG
GAACAGACTTTTCCCCAATGCAGATCCCAAAGCTCTAGACTTACTGGACAAGATGCTGACTTTCAACCCC
CATAAAAGAATTGAAGTAGAGGCAGCTTTGGCTCATCCTTATCTGGAGCAGTATTATGACCCAAGTGATG
AGCCTGTAGCTGAAGCTCCCTTTAAATTTGAAATGGAGCTTGATGATTTGCCCAAGGAGACTCTTAAGGA
GCTAATTTTTGAAGAAACCGCTAGATTCCAGCCAGGGTACTAATCGCGCCTCTAGAGGATCCGTTAACTA
ACTTAAGCTAGCGTCGACGGGCCGCGGTAACAATTGTTAACTAACTTAAGCTAGCAACGGTTTCCCTCTA
GCGGGATCAATTCCGCCCCCCCCCCCTAACGTTACTGGCCGAAGCCGCTTGGAATAAGGCCGGTGTGCGT
TTGTCTATATGTTATTTTCCACCATATTGCCGTCTTTTGGCAATGTGAGGGCCCGGAAACCTGGCCCTGT
CTTCTTGACGAGCATTCCTAGGGGTCTTTCCCCTCTCGCCAAAGGAATGCAAGGTCTGTTGAATGTCGTG
AAGGAAGCAGTTCCTCTGGAAGCTTCTTGAAGACAAACAACGTCTGTAGCGACCCTTTGCAGGCAGCGGA
ACCCCCCACCTGGCGACAGGTGCCTCTGCGGCCAAAAGCCACGTGTATAAGATACACCTGCAAAGGCGGC
ACAACCCCAGTGCCACGTTGTGAGTTGGATAGTTGTGGAAAGAGTCAAATGGCTCTCCTCAAGCGTATTC
AACAAGGGGCTGAAGGATGCCCAGAAGGTACCCCATTGTATGGGATCTGATCTGGGGCCTCGGTGCACAT
GCTTTACATGTGTTTAGTCGAGGTTAAAAAACGTCTAGGCCCCCCGAACCACGGGGACGTGGTTTTCCTT
TGAAAAACACGATAATACCATGGTCATGAAAACATTTAACATTTCTCAACAAGATCTAGAATTAGTAGAA
GTAGCGACAGAGAAGATTACAATGCTTTATGAGGATAATAAACATCATGTGGGAGCGGCAATTCGTACGA
AAACAGGAGAAATCATTTCGGCAGTACATATTGAAGCGTATATAGGACGAGTAACTGTTTGTGCAGAAGC
CATTGCGATTGGTAGTGCAGTTTCGAATGGACAAAAGGATTTTGACACGATTGTAGCTGTTAGACACCCT
TATTCTGACGAAGTAGATAGAAGTATTCGAGTGGTAAGTCCTTGTGGTATGTGTAGGGAGTTGATTTCAG
ACTATGCACCAGATTGTTTTGTGTTAATAGAAATGAATGGCAAGTTAGTCAAAACTACGATTGAAGAACT
CATTCCACTCAAATATACCCGAAATTAA
```

**Annotation**

MEK: MEK1

P2A: 2A self-cleaving peptide

mCherry: red fluorescent protein mCherry

ERK2: ERK2

ECMV IRES: internal ribosomal entry site from *Encephalomyocarditis* virus

Bsr: blasticidin S-deaminase

**m**

**pCMV-mTagBFP2-SOS1cat**

```
>Amino acid sequence
MVSKGEELIKENMHMKLYMEGTVDNHHFKCTSEGEGKPYEGTQTMRIKVVEGGPLPFAFDILATSFLYGS
KTFINHTQGIPDFFKQSFPEGFTWERVTTYEDGGVLTATQDTSLQDGCLIYNVKIRGVNFTSNGPVMQKK
TLGWEAFTETLYPADGGLEGRNDMALKLVGGSHLIANAKTTYRSKKPAKNLKMPGVYYVDYRLERIKEAN
NETYVEQHEVAVARYCDLPSKLGHKLNRSRAQASQMRLPSADVYRFAEPDSEENIIFEENMQPKAGIPII
KAGTVIKLIERLTYHMYADPNFVRTFLTTYRSFCKPQELLSLIIERFEIPEPEPTEADRIAIENGDQPLS
AELKRFRKEYIQPVQLRVLNVCRHWVEHHFYDFERDAYLLQRMEEFIGTVRGKAMKKWVESITKIIQRKK
IARDNGPGHNITFQSSPPTVEWHISRPGHIETFDLLTLHPIEIARQLTLLESDLYRAVQPSELVGSVWTK
EDKEINSPNLLKMIRHTTNLTLWFEKCIVETENLEERVAVVSRIIEILQVFQELNNFNGVLEVVSAMNSS
PVYRLDHTFEQIPSRQKKILEEAHELSEDHYKKYLAKLRSINPPCVPFFGIYLTNILKTEEGNPEVLKRH
GKELINFSKRRKVAEITGEIQQYQNQPYCLRVESDIKRFFENLNPMGNSMEKEFTDYLFNKSLEIEPRNP
KPLPRFPKKYSYPLKSPGVRPSNPRPGT*

>DNA sequence
ATGGTGTCTAAGGGCGAAGAGCTGATTAAGGAGAACATGCACATGAAGCTGTACATGGAGGGCACCGTGG
ACAACCATCACTTCAAGTGCACATCCGAGGGCGAAGGCAAGCCCTACGAGGGCACCCAGACCATGAGAAT
CAAGGTGGTCGAGGGCGGCCCTCTCCCCTTCGCCTTCGACATCCTGGCTACTAGCTTCCTCTACGGCAGC
AAGACCTTCATCAACCACACCCAGGGCATCCCCGACTTCTTCAAGCAGTCCTTCCCTGAGGGCTTCACAT
GGGAGAGAGTCACCACATACGAAGACGGGGGCGTGCTGACCGCTACCCAGGACACCAGCCTCCAGGACGG
CTGCCTCATCTACAACGTCAAGATCAGAGGGGTGAACTTCACATCCAACGGCCCTGTGATGCAGAAGAAA
ACACTCGGCTGGGAGGCCTTCACCGAGACGCTGTACCCCGCTGACGGCGGCCTGGAAGGCAGAAACGACA
TGGCCCTGAAGCTCGTGGGCGGGAGCCATCTGATCGCAAACGCCAAGACCACATATAGATCCAAGAAACC
CGCTAAGAACCTCAAGATGCCTGGCGTCTACTATGTGGACTACAGACTGGAAAGAATCAAGGAGGCCAAC
AACGAGACCTACGTCGAGCAGCACGAGGTGGCAGTGGCCAGATACTGCGACCTCCCTAGCAAACTGGGGC
ACAAGCTTAATAGATCTCGAGCTCAAGCTTCGCAGATGAGGCTGCCTAGTGCTGATGTTTATAGATTTGC
AGAGCCTGACTCTGAAGAGAATATTATATTTGAAGAGAACATGCAGCCCAAGGCTGGAATTCCAATTATC
AAAGCAGGAACTGTTATTAAACTTATAGAGAGGCTTACGTACCATATGTACGCAGATCCCAATTTTGTTC
GGACATTTCTTACAACATACAGATCCTTTTGCAAACCTCAAGAACTACTGAGTCTTATAATAGAAAGGTT
TGAAATTCCAGAGCCTGAGCCAACAGAAGCTGATCGCATAGCTATAGAGAATGGAGATCAACCCTTGAGT
GCAGAACTGAAAAGATTTAGAAAAGAATATATACAGCCTGTGCAACTGCGAGTATTAAATGTATGTCGGC
ACTGGGTAGAGCACCACTTCTATGATTTTGAAAGAGATGCATATCTTTTGCAACGAATGGAAGAATTTAT
TGGAACAGTAAGAGGTAAAGCAATGAAAAAATGGGTTGAATCCATCACTAAAATAATCCAAAGGAAAAAA
ATTGCAAGAGACAATGGACCAGGTCATAATATTACATTTCAGAGTTCACCTCCCACAGTTGAGTGGCATA
TAAGCAGACCTGGGCACATAGAGACTTTTGACCTGCTCACCTTACACCCAATAGAAATTGCTCGACAACT
CACTTTACTTGAATCAGATCTATACCGAGCTGTACAGCCATCAGAATTAGTTGGAAGTGTGTGGACAAAA
GAAGACAAAGAAATTAACTCTCCTAATCTTCTGAAAATGATTCGACATACCACCAACCTCACTCTGTGGT
TTGAGAAATGTATTGTAGAAACTGAAAATTTAGAAGAAAGAGTAGCTGTGGTGAGTCGAATTATTGAGAT
TCTACAAGTCTTTCAAGAGTTGAACAACTTTAATGGTGTCCTTGAGGTTGTCAGTGCTATGAATTCATCA
CCTGTTTACAGACTAGACCACACATTTGAGCAAATACCAAGTCGCCAGAAGAAAATTTTAGAAGAAGCTC
ATGAATTGAGTGAAGATCACTATAAGAAATATTTGGCAAAACTCAGGTCTATTAATCCACCATGTGTGCC
TTTCTTTGGAATTTATCTCACTAATATCTTGAAAACAGAAGAAGGCAACCCTGAGGTCCTAAAAAGACAT
GGAAAAGAGCTTATAAACTTTAGCAAAAGGAGGAAAGTAGCAGAAATAACAGGAGAGATCCAGCAGTACC
AAAATCAGCCTTACTGTTTACGAGTAGAATCAGATATCAAAAGGTTCTTTGAAAACTTGAATCCGATGGG
AAATAGCATGGAGAAGGAATTTACAGATTATCTTTTCAACAAATCCCTAGAAATAGAACCACGAAACCCT
AAGCCTCTCCCAAGATTTCCAAAAAAATATAGCTATCCCCTAAAATCTCCTGGTGTTCGTCCATCAAACC
CAAGACCAGGTACCTAA
```

**Annotation**

mTagBFP2: blue fluorescent protein mTagBFP2

$SOS1_{cat}$: Rem-Cdc25 (RasGEF) domain from human SOS1 (residues 566-1049)

**n**

**pCMV-mCherry-FKBP-TEVp**

```
>Amino acid sequence
MVSKGEEDNMAIIKEFMRFKVHMEGSVNGHEFEIEGEGEGRPYEGTQTAKLKVTKGGPLPFAWDILSPQF
MYGSKAYVKHPADIPDYLKLSFPEGFKWERVMNFEDGGVVTVTQDSSLQDGEFIYKVKLRGTNFPSDGPV
MQKKTMGWEASSERMYPEDGALKGEIKQRLKLKDGGHYDAEVKTTYKAKKPVQLPGAYNVNIKLDITSHN
EDYTIVEQYERAEGRHSTGGMDELYKSGLRSRAMGVQVETISPGDGRTFPKRGQTCVVHYTGMLEDGKKF
DSSRDRNKPFKFMLGKQEVIRGWEEGVAQMSVGQRAKLTISPDYAYGATGHPGIIPPHATLVFDVELLKL
EEFCSRRYRMGESLFKGPRDYNPISSTICHLTNESDGHTTSLYGIGFGPFIITNKHLFRRNNGTLLVQSL
HGVFKVKNTTTLQQHLIDGRDMIIIRMPKDFPPFPQKLKFREPQREERICLVTTNFQTKSMSSMVSDTSC
TFPSSDGIFWKHWIQTKDGQCGSPLVSTRDGFIVGIHSASNFTNTNNYFTSVPKNFMELLTNQEAQQWVS
GWRLNADSVLWGGHKVFMVGSTGSR*

>DNA sequence
ATGGTGAGCAAGGGCGAGGAGGATAACATGGCCATCATCAAGGAGTTCATGCGCTTCAAGGTGCACATGG
AGGGCTCCGTGAACGGCCACGAGTTCGAGATCGAGGGCGAGGGCGAGGGCCGCCCCTACGAGGGCACCCA
GACCGCCAAGCTGAAGGTGACCAAGGGTGGCCCCCTGCCCTTCGCCTGGGACATCCTGTCCCCTCAGTTC
ATGTACGGCTCCAAGGCCTACGTGAAGCACCCCGCCGACATCCCCGACTACTTGAAGCTGTCCTTCCCCG
AGGGCTTCAAGTGGGAGCGCGTGATGAACTTCGAGGACGGCGGCGTGGTGACCGTGACCCAGGACTCCTC
CCTGCAGGACGGCGAGTTCATCTACAAGGTGAAGCTGCGCGGCACCAACTTCCCCTCCGACGGCCCCGTA
ATGCAGAAGAAGACCATGGGCTGGGAGGCCTCCTCCGAGCGGATGTACCCCGAGGACGGCGCCCTGAAGG
GCGAGATCAAGCAGAGGCTGAAGCTGAAGGACGGCGGCCACTACGACGCTGAGGTCAAGACCACCTACAA
GGCCAAGAAGCCCGTGCAGCTGCCCGGCGCCTACAACGTCAACATCAAGTTGGACATCACCTCCCACAAC
GAGGACTACACCATCGTGGAACAGTACGAACGCGCCGAGGGCCGCCACTCCACCGGCGGCATGGACGAGC
TGTACAAGTCCGGACTCAGATCTCGAGCTATGGGAGTGCAGGTGGAAACCATCTCCCCAGGAGACGGGCG
CACCTTCCCCAAGCGCGGCCAGACCTGCGTGGTGCACTACACCGGGATGCTTGAAGATGGAAAGAAATTT
GATTCCTCCCGGGACAGAAACAAGCCCTTTAAGTTTATGCTAGGCAAGCAGGAGGTGATCCGAGGCTGGG
AAGAAGGGGTTGCCCAGATGAGTGTGGGTCAGAGAGCCAAACTGACTATATCTCCAGATTATGCCTATGG
TGCCACTGGGCACCCAGGCATCATCCCACCACATGCCACTCTCGTCTTCGATGTGGAGCTTCTAAAACTG
GAAGAATTCTGCAGTCGACGGTACCGCATGGGCGAGAGCCTTTTCAAGGGCCCGAGGGACTACAACCCGA
TCTCCAGCACCATCTGTCACCTGACCAACGAGAGCGACGGTCACACCACTAGTCTGTACGGCATCGGCTT
CGGCCCCTTCATCATCACCAACAAGCATCTGTTCAGGAGGAATAACGGCACACTGCTGGTGCAAAGCCTG
CACGGCGTGTTCAAAGTGAAGAACACAACCACCCTGCAACAGCACCTGATCGACGGCAGGGACATGATTA
TCATCAGGATGCCCAAGGACTTCCCCCCCTTTCCCCAGAAACTGAAGTTCAGGGAGCCACAAAGGGAGGA
GCGAATCTGCCTGGTGACCACCAACTTCCAGACCaagTCCATGAGCAGCATGGTCTCTGATACCAGCTGC
ACCTTCCCCAGCAGCGACGGCATCTTCTGGAAGCACTGGATTCAGACGAAGGATGGCCAATGCGGCAGCC
CATTGGTGAGCACTAGGGACGGCTTCATCGTGGGCATCCACAGCGCCAGCAATTTTACCAATACCAACAA
CTACTTCACGAGCGTGCCGAAAAACTTCATGGAGCTGTTGACCAATCAAGAGGCGCAGCAGTGGGTGAGC
GGCTGGAGGCTGAACGCCGACAGCGTTCTTTGGGGCGGACATAAGGTGTTCATGGTCGGATCCACCGGAT
CTAGATAA
```

**Annotation**

mCherry: red fluorescent protein mCherry

FKBP: FKBP12

TEVp: TEV protease lacking the C-terminal residues 220-242

**o**

**pCMV-PB1-AG-FRB-TEVcs-BFP-HA**

>Amino acid sequence
MASLTVKAYLLGKEDAAREIRRFSFCCSPEPEAEAEAAAGPGPCERLLSRVAALFPALRPGGFQAHYRDE
DGDLVAFSSDEELTMAMSYVKDDIFRIYIKEKTGSGSGGSGAGGSAGSGAGGSAGSGAGGSAGSGAGGSA
GSGAGGSASGSSPGSGSGGSGAGGSAGSGAGGSAGSGAGGSAGSGAGGSAGSGAGGSASGSSPGNSADGG
GGSGGSGGSGGGSTQGGSMVSVIKPEMKIKLCMRGTVNGHNFVIEGEGKGNPYEGTQILDLNVTEGAPLP
FAYDILTTVFQYGNRAFTKYPADIQDYFKQTFPEGYHWERSMTYEDQGICTATSNISMRGDCFFYDIRFD
GVNFPPNGPVMQKKTLKWEPSTEKMYVRDGVLKGDVNMALLLEGGGHYRCDFKTTYKAKKDVRLPDYHFV
DHRIEILKHDKDYNKVKLYENAVARYSMLPSQAKGTGTAAENSGNSRTKLMILWHEMWHEGLEEASRLYF
GERNVKGMFEVLEPLHAMMERGPQTLKETSFNQAYGRDLMEAQEWCRKYMKSGNVKDLLQAWDLYYHVFR
RISKPGGSGAGSGAGSGAGENLYFQLVDGGSAGGSGGVDMVSKGEELIKENMHMKLYMEGTVDNHHFKCT
SEGEGKPYEGTQTMRIKVVEGGPLPFAFDILATSFLYGSKTFINHTQGIPDFFKQSFPEGFTWERVTTYE
DGGVLTATQDTSLQDGCLIYNVKIRGVNFTSNGPVMQKKTLGWEAFTETLYPADGGLEGRNDMALKLVGG
SHLIANAKTTYRSKKPAKNLKMPGVYYVDYRLERIKEANNETYVEQHEVAVARYCDLPSKLGHKLNRSGS
YPYDVPDYARS*

>DNA sequence
ATGGCGTCGCTCACCGTGAAGGCCTACCTTCTGGGCAAGGAGGACGCGGCGCGCGAGATTCGCCGCTTCA
GCTTCTGTTGCAGCCCCGAGCCTGAGGCGGAAGCCGAGGCTGCGGCGGGTCCGGGACCCTGCGAGCGGCT
GCTGAGCCGGGTGGCCGCCCTGTTCCCCGCGCTGCGGCCTGGCGGCTTCCAGGCGCACTACCGCGATGAG
GACGGGGACTTGGTTGCCTTTTCCAGTGACGAGGAATTGACAATGGCCATGTCCTACGTGAAGGATGACA
TCTTCCGAATCTACATTAAAGAGAAAACCGGTTCTGGGAGTGGCGGTAGTGGTGCTGGAGGCAGCGCAGG
CTCTGGAGCAGGCGGCAGTGCCGGAAGTGGCGCTGGAGGGTCTGCCGGATCTGGAGCCGGTGGTTCTGCC
GGCTCCGGCGCTGGTGGGAGCGCTTCCGGAAGTAGTCCCGGTTCTGGGAGTGGCGGTAGTGGTGCTGGAG
GCAGCGCAGGCTCTGGAGCAGGCGGCAGTGCCGGAAGTGGCGCTGGAGGGTCTGCCGGATCTGGAGCCGG
TGGTTCTGCCGGCTCCGGCGCTGGTGGGAGCGCTTCCGGAAGTAGTCCCGGTAATTCCGCTGACGGCGGC
GGAGGATCGGGTGGTAGTGGTGGTTCAGGAGGAGGATCGACCCAAGGAGGATCCATGGTGAGCGTGATCA
AGCCCGAGATGAAGATCAAGCTGTGCATGAGGGGCACCGTGAACGGCCACAACTTCGTGATCGAGGGCGA
GGGCAAGGGCAACCCCTACGAGGGCACCCAGATCCTGGACCTGAACGTGACCGAGGGCGCCCCCCTGCCC
TTCGCCTACGACATCCTGACCACCGTGTTCCAGTACGGCAACAGGGCCTTCACCAAGTACCCCGCCGACA
TCCAGGACTACTTCAAGCAGACCTTCCCCGAGGGCTACCACTGGGAGAGGAGCATGACCTACGAGGACCA
GGGCATCTGCACCGCCACCAGCAACATCAGCATGAGGGGCGACTGCTTCTTCTACGACATCAGGTTCGAC
GGCGTGAACTTCCCCCCCAACGGCCCCGTGATGCAGAAGAAGACCCTGAAGTGGGAGCCCAGCACCGAGA
AGATGTACGTGAGGGACGGCGTGCTGAAGGGCGACGTGAACATGGCCCTGCTGCTGGAGGGCGGCGGCCA
CTACAGGTGCGACTTCAAGACCACCTACAAGGCCAAGAAGGACGTGAGGCTGCCCGACTACCACTTCGTG
GACCACAGGATCGAGATCCTGAAGCACGACAAGGACTACAACAAGGTGAAGCTGTACGAGAACGCCGTGG
CCAGGTACAGCATGCTGCCCAGCCAGGCCAAGGGTACCGGAACTGCAGCAGAGAATTCGGGAAACTCGAG
AACAAAGCTTATGATCCTCTGGCATGAGATGTGGCATGAAGGCCTGGAAGAGGCATCTCGTTTGTACTTT
GGGGAAAGGAACGTGAAAGGCATGTTTGAGGTGCTGGAGCCCTTGCATGCTATGATGGAACGGGGCCCCC
AGACTCTGAAGGAAACATCCTTTAATCAGGCCTATGGTCGAGATTTAATGGAGGCCCAAGAGTGGTGCAG
GAAGTACATGAAATCAGGGAATGTCAAGGACCTCCTCCAAGCCTGGGACCTCTATTATCATGTGTTCCGA
CGAATCTCAAAGCCCGGGGGTAGTGGTGCTGGCTCTGGTGCTGGTAGTGGCGCTGGTGAAAACCTGTATT
TTCAGCTGGTCGATGGTGGTAGTGCTGGTGGTTCCGGAGGTGTCGACATGGTGTCTAAGGGCGAAGAGCT
GATTAAGGAGAACATGCACATGAAGCTGTACATGGAGGGCACCGTGGACAACCATCACTTCAAGTGCACA
TCCGAGGGCGAAGGCAAGCCCTACGAGGGCACCCAGACCATGAGAATCAAGGTGGTCGAGGGCGGCCCTC
TCCCCTTCGCCTTCGACATCCTGGCTACTAGCTTCCTCTACGGCAGCAAGACCTTCATCAACCACACCCA
GGGCATCCCCGACTTCTTCAAGCAGTCCTTCCCTGAGGGCTTCACATGGGAGAGAGTCACCACATACGAA
GACGGGGGCGTGCTGACCGCTACCCAGGACACCAGCCTCCAGGACGGCTGCCTCATCTACAACGTCAAGA
TCAGAGGGGTGAACTTCACATCCAACGGCCCTGTGATGCAGAAGAAAACACTCGGCTGGGAGGCCTTCAC
CGAGACGCTGTACCCCGCTGACGGCGGCCTGGAAGGCAGAAACGACATGGCCCTGAAGCTCGTGGGCGGG
AGCCATCTGATCGCAAACGCCAAGACCACATATAGATCCAAGAAACCCGCTAAGAACCTCAAGATGCCTG
GCGTCTACTATGTGGACTACAGACTGGAAAGAATCAAGGAGGCCAACAACGAGACCTACGTCGAGCAGCA

CGAGGTGGCAGTGGCCAGATACTGCGACCTCCCTAGCAAACTGGGGCACAAGCTTAATAGATCCGGCTCTTACCCATACGATGTTCCAGATTACGCTAGATCTTAA

**Annotation**

PB1: PB1 domain from human p62 (residues 1-102)

AG: green fluorescent protein Azami-Green

FRB: FRB domain from human mammalian target of rapamycin (mTOR) (residues 2021-2113 with a T2098L mutation)

TEV$_{cs}$: TEV protease cleavage sequence (ENLYFQ/L)

mTagBFP2: blue fluorescent protein mTagBFP2

HA: HA epitope tag

**p**

**pCMV-mCherry-FKBP**

>Amino acid sequence
MVSKGEEDNMAIIKEFMRFKVHMEGSVNGHEFEIEGEGEGRPYEGTQTAKLKVTKGGPLPFAWDILSPQF
MYGSKAYVKHPADIPDYLKLSFPEGFKWERVMNFEDGGVVTVTQDSSLQDGEFIYKVKLRGTNFPSDGPV
MQKKTMGWEASSERMYPEDGALKGEIKQRLKLKDGGHYDAEVKTTYKAKKPVQLPGAYNVNIKLDITSHN
EDYTIVEQYERAEGRHSTGGMDELYKGSGASAGGGSMGVQVETISPGDGRTFPKRGQTCVVHYTGMLEDG
KKFDSSRDRNKPFKFMLGKQEVIRGWEEGVAQMSVGQRAKLTISPDYAYGATGHPGIIPPHATLVFDVEL
LKLE*

>DNA sequence
ATGGTGAGCAAGGGCGAGGAGGATAACATGGCCATCATCAAGGAGTTCATGCGCTTCAAGGTGCACATGG
AGGGCTCCGTGAACGGCCACGAGTTCGAGATCGAGGGCGAGGGCGAGGGCCGCCCCTACGAGGGCACCCA
GACCGCCAAGCTGAAGGTGACCAAGGGTGGCCCCCTGCCCTTCGCCTGGGACATCCTGTCCCCTCAGTTC
ATGTACGGCTCCAAGGCCTACGTGAAGCACCCCGCCGACATCCCCGACTACTTGAAGCTGTCCTTCCCCG
AGGGCTTCAAGTGGGAGCGCGTGATGAACTTCGAGGACGGCGGCGTGGTGACCGTGACCCAGGACTCCTC
CCTGCAGGACGGCGAGTTCATCTACAAGGTGAAGCTGCGCGGCACCAACTTCCCCTCCGACGGCCCCGTA
ATGCAGAAGAAGACCATGGGCTGGGAGGCCTCCTCCGAGCGGATGTACCCCGAGGACGGCGCCCTGAAGG
GCGAGATCAAGCAGAGGCTGAAGCTGAAGGACGGCGGCCACTACGACGCTGAGGTCAAGACCACCTACAA
GGCCAAGAAGCCCGTGCAGCTGCCCGGCGCCTACAACGTCAACATCAAGTTGGACATCACCTCCCACAAC
GAGGACTACACCATCGTGGAACAGTACGAACGCGCCGAGGGCCGCCACTCCACCGGCGGCATGGACGAGC
TGTACAAGGGCTCCGGTGCCAGTGCTGGTGGTGGCAGCATGGGAGTGCAGGTGGAAACCATCTCCCCAGG
AGACGGGCGCACCTTCCCCAAGCGCGGCCAGACCTGCGTGGTGCACTACACCGGGATGCTTGAAGATGGA
AAGAAATTTGATTCCTCCCGGGACAGAAACAAGCCCTTTAAGTTTATGCTAGGCAAGCAGGAGGTGATCC
GAGGCTGGGAAGAAGGGGTTGCCCAGATGAGTGTGGGTCAGAGAGCCAAACTGACTATATCTCCAGATTA
TGCCTATGGTGCCACTGGGCACCCAGGCATCATCCCACCACATGCCACTCTCGTCTTCGATGTGGAGCTT
CTAAAACTGGAATGA

**Annotation**
mCherry: red fluorescent protein mCherry
FKBP: FKBP12

**q**

**pCMV-PB1-AG-FRB-TEVuncs-BFP-HA**

>Amino acid sequence
MASLTVKAYLLGKEDAAREIRRFSFCCSPEPEAEAEAAAGPGPCERLLSRVAALFPALRPGGFQAHYRDE
DGDLVAFSSDEELTMAMSYVKDDIFRIYIKEKTGSGSGGSGAGGSAGSGAGGSAGSGAGGSAGSGAGGSA
GSGAGGSASGSSPGSGSGGSGAGGSAGSGAGGSAGSGAGGSAGSGAGGSAGSGAGGSASGSSPGNSADGG
GGSGGSGGSGGGSTQGGSMVSVIKPEMKIKLCMRGTVNGHNFVIEGEGKGNPYEGTQILDLNVTEGAPLP
FAYDILTTVFQYGNRAFTKYPADIQDYFKQTFPEGYHWERSMTYEDQGICTATSNISMRGDCFFYDIRFD
GVNFPPNGPVMQKKTLKWEPSTEKMYVRDGVLKGDVNMALLLEGGGHYRCDFKTTYKAKKDVRLPDYHFV
DHRIEILKHDKDYNKVKLYENAVARYSMLPSQAKGTGTAAENSGNSRTKLMILWHEMWHEGLEEASRLYF
GERNVKGMFEVLEPLHAMMERGPQTLKETSFNQAYGRDLMEAQEWCRKYMKSGNVKDLLQAWDLYYHVFR
RISKPGGSGAGSGAGSGAGNEYLQFLVDGGSAGGSGGVDMVSKGEELIKENMHMKLYMEGTVDNHHFKCT
SEGEGKPYEGTQTMRIKVVEGGPLPFAFDILATSFLYGSKTFINHTQGIPDFFKQSFPEGFTWERVTTYE
DGGVLTATQDTSLQDGCLIYNVKIRGVNFTSNGPVMQKKTLGWEAFTETLYPADGGLEGRNDMALKLVGG
SHLIANAKTTYRSKKPAKNLKMPGVYYVDYRLERIKEANNETYVEQHEVAVARYCDLPSKLGHKLNRSGS
YPYDVPDYARS*

>DNA sequence
ATGGCGTCGCTCACCGTGAAGGCCTACCTTCTGGGCAAGGAGGACGCGGCGCGCGAGATTCGCCGCTTCA
GCTTCTGTTGCAGCCCCGAGCCTGAGGCGGAAGCCGAGGCTGCGGCGGGTCCGGGACCCTGCGAGCGGCT
GCTGAGCCGGGTGGCCGCCCTGTTCCCCGCGCTGCGGCCTGGCGGCTTCCAGGCGCACTACCGCGATGAG
GACGGGGACTTGGTTGCCTTTTCCAGTGACGAGGAATTGACAATGGCCATGTCCTACGTGAAGGATGACA
TCTTCCGAATCTACATTAAAGAGAAAACCGGTTCTGGGAGTGGCGGTAGTGGTGCTGGAGGCAGCGCAGG
CTCTGGAGCAGGCGGCAGTGCCGGAAGTGGCGCTGGAGGGTCTGCCGGATCTGGAGCCGGTGGTTCTGCC
GGCTCCGGCGCTGGTGGGAGCGCTTCCGGAAGTAGTCCCGGTTCTGGGAGTGGCGGTAGTGGTGCTGGAG
GCAGCGCAGGCTCTGGAGCAGGCGGCAGTGCCGGAAGTGGCGCTGGAGGGTCTGCCGGATCTGGAGCCGG
TGGTTCTGCCGGCTCCGGCGCTGGTGGGAGCGCTTCCGGAAGTAGTCCCGGTAATTCCGCTGACGGCGGC
GGAGGATCGGGTGGTAGTGGTGGTTCAGGAGGAGGATCGACCCAAGGAGGATCCATGGTGAGCGTGATCA
AGCCCGAGATGAAGATCAAGCTGTGCATGAGGGGCACCGTGAACGGCCACAACTTCGTGATCGAGGGCGA
GGGCAAGGGCAACCCCTACGAGGGCACCCAGATCCTGGACCTGAACGTGACCGAGGGCGCCCCCCTGCCC
TTCGCCTACGACATCCTGACCACCGTGTTCCAGTACGGCAACAGGGCCTTCACCAAGTACCCCGCCGACA
TCCAGGACTACTTCAAGCAGACCTTCCCCGAGGGCTACCACTGGGAGAGGAGCATGACCTACGAGGACCA
GGGCATCTGCACCGCCACCAGCAACATCAGCATGAGGGGCGACTGCTTCTTCTACGACATCAGGTTCGAC
GGCGTGAACTTCCCCCCCAACGGCCCCGTGATGCAGAAGAAGACCCTGAAGTGGGAGCCCAGCACCGAGA
AGATGTACGTGAGGGACGGCGTGCTGAAGGGCGACGTGAACATGGCCCTGCTGCTGGAGGGCGGCGGCCA
CTACAGGTGCGACTTCAAGACCACCTACAAGGCCAAGAAGGACGTGAGGCTGCCCGACTACCACTTCGTG
GACCACAGGATCGAGATCCTGAAGCACGACAAGGACTACAACAAGGTGAAGCTGTACGAGAACGCCGTGG
CCAGGTACAGCATGCTGCCCAGCCAGGCCAAGGGTACCGGAACTGCAGCAGAGAATTCGGGAAACTCGAG
AACAAAGCTTATGATCCTCTGGCATGAGATGTGGCATGAAGGCCTGGAAGAGGCATCTCGTTTGTACTTT
GGGGAAAGGAACGTGAAAGGCATGTTTGAGGTGCTGGAGCCCTTGCATGCTATGATGGAACGGGGCCCCC
AGACTCTGAAGGAAACATCCTTTAATCAGGCCTATGGTCGAGATTTAATGGAGGCCCAAGAGTGGTGCAG
GAAGTACATGAAATCAGGGAATGTCAAGGACCTCCTCCAAGCCTGGGACCTCTATTATCATGTGTTCCGA
CGAATCTCAAAGCCCGGGGGTAGTGGTGCTGGCTCTGGTGCTGGTAGTGGCGCTGGTAACGAATATCTGC
AGTTTCTGGTCGATGGTGGTAGTGCTGGTGGTTCCGGAGGTGTCGACATGGTGTCTAAGGGCGAAGAGCT
GATTAAGGAGAACATGCACATGAAGCTGTACATGGAGGGCACCGTGGACAACCATCACTTCAAGTGCACA
TCCGAGGGCGAAGGCAAGCCCTACGAGGGCACCCAGACCATGAGAATCAAGGTGGTCGAGGGCGGCCCTC
TCCCCTTCGCCTTCGACATCCTGGCTACTAGCTTCCTCTACGGCAGCAAGACCTTCATCAACCACACCCA
GGGCATCCCCGACTTCTTCAAGCAGTCCTTCCCTGAGGGCTTCACATGGGAGAGAGTCACCACATACGAA
GACGGGGGCGTGCTGACCGCTACCCAGGACACCAGCCTCCAGGACGGCTGCCTCATCTACAACGTCAAGA
TCAGAGGGGTGAACTTCACATCCAACGGCCCTGTGATGCAGAAGAAAACACTCGGCTGGGAGGCCTTCAC
CGAGACGCTGTACCCCGCTGACGGCGGCCTGGAAGGCAGAAACGACATGGCCCTGAAGCTCGTGGGCGGG
AGCCATCTGATCGCAAACGCCAAGACCACATATAGATCCAAGAAACCCGCTAAGAACCTCAAGATGCCTG
GCGTCTACTATGTGGACTACAGACTGGAAAGAATCAAGGAGGCCAACAACGAGACCTACGTCGAGCAGCA

CGAGGTGGCAGTGGCCAGATACTGCGACCTCCCTAGCAAACTGGGGCACAAGCTTAATAGATCCGGCTCT
TACCCATACGATGTTCCAGATTACGCTAGATCTTAA

**Annotation**

PB1: PB1 domain from human p62 (residues 1-102)

AG: green fluorescent protein Azami-Green

FRB: FRB domain from human mammalian target of rapamycin (mTOR) (residues 2021-2113 with a T2098L mutation)

TEV$_{uncs}$: TEV protease uncleavable sequence (NEYLQFL)

mTagBFP2: blue fluorescent protein mTagBFP2

HA: HA epitope tag

**r**

**pCMV-PB1-AG-FRB-TEVcs-BFP-HA-Vav2cat**

>Amino acid sequence
MASLTVKAYLLGKEDAAREIRRFSFCCSPEPEAEAEAAAGPGPCERLLSRVAALFPALRPGGFQAHYRDE
DGDLVAFSSDEELTMAMSYVKDDIFRIYIKEKTGSGSGGSGAGGSAGSGAGGSAGSGAGGSAGSGAGGSA
GSGAGGSASGSSPGSGSGGSGAGGSAGSGAGGSAGSGAGGSAGSGAGGSAGSGAGGSASGSSPGNSADGG
GGSGGSGGSGGGSTQGGSMVSVIKPEMKIKLCMRGTVNGHNFVIEGEGKGNPYEGTQILDLNVTEGAPLP
FAYDILTTVFQYGNRAFTKYPADIQDYFKQTFPEGYHWERSMTYEDQGICTATSNISMRGDCFFYDIRFD
GVNFPPNGPVMQKKTLKWEPSTEKMYVRDGVLKGDVNMALLLEGGGHYRCDFKTTYKAKKDVRLPDYHFV
DHRIEILKHDKDYNKVKLYENAVARYSMLPSQAKGTGTAAENSGNSRTKLMILWHEMWHEGLEEASRLYF
GERNVKGMFEVLEPLHAMMERGPQTLKETSFNQAYGRDLMEAQEWCRKYMKSGNVKDLLQAWDLYYHVFR
RISKPGGSGAGSGAGSGAGENLYFQLVDGGSAGGSGGVDMVSKGEELIKENMHMKLYMEGTVDNHHFKCT
SEGEGKPYEGTQTMRIKVVEGGPLPFAFDILATSFLYGSKTFINHTQGIPDFFKQSFPEGFTWERVTTYE
DGGVLTATQDTSLQDGCLIYNVKIRGVNFTSNGPVMQKKTLGWEAFTETLYPADGGLEGRNDMALKLVGG
SHLIANAKTTYRSKKPAKNLKMPGVYYVDYRLERIKEANNETYVEQHEVAVARYCDLPSKLGHKLNRSGS
YPYDVPDYARSRGPMKMGMTEDDKRSCCLLEIQETEAKYYRTLEDIEKNYMGPLRLVLSPADMAAVFINL
EDLIKVHHSFLRAIDVSMMAGGSTLAKVFLEFKERLLIYGEYCSHMEHAQSTLNQLLASREDFRQKVEEC
TLRVQDGKFKLQDLLVVPMQRVLKYHLLLKELLSHSADRPERQQLKEALEAMQDLAMYINEVKRDKETLK
KISEFQCSIENLQVKLEEFGRPKIDGELKVRSIVNHTKQDRYLFLFDKVVIVCKRKGYSYELKEVIELLF
HKMTDDPMHNKDIKKWSYGFYLIHLQGKQGFQFFCKTEDMKRKWMEQFEMAMSNIKPDKANANHHSFQMY
TFDKTTNCKACKMFLRGTFYQGYLCTRCGVGAHKECLEVIPPCK*

>DNA sequence
ATGGCGTCGCTCACCGTGAAGGCCTACCTTCTGGGCAAGGAGGACGCGGCGCGCGAGATTCGCCGCTTCA
GCTTCTGTTGCAGCCCCGAGCCTGAGGCGGAAGCCGAGGCTGCGGCGGGTCCGGGACCCTGCGAGCGGCT
GCTGAGCCGGGTGGCCGCCCTGTTCCCCGCGCTGCGGCCTGGCGGCTTCCAGGCGCACTACCGCGATGAG
GACGGGGACTTGGTTGCCTTTTCCAGTGACGAGGAATTGACAATGGCCATGTCCTACGTGAAGGATGACA
TCTTCCGAATCTACATTAAAGAGAAAACCGGTTCTGGGAGTGGCGGTAGTGGTGCTGGAGGCAGCGCAGG
CTCTGGAGCAGGCGGCAGTGCCGGAAGTGGCGCTGGAGGGTCTGCCGGATCTGGAGCCGGTGGTTCTGCC
GGCTCCGGCGCTGGTGGGAGCGCTTCCGGAAGTAGTCCCGGTTCTGGGAGTGGCGGTAGTGGTGCTGGAG
GCAGCGCAGGCTCTGGAGCAGGCGGCAGTGCCGGAAGTGGCGCTGGAGGGTCTGCCGGATCTGGAGCCGG
TGGTTCTGCCGGCTCCGGCGCTGGTGGGAGCGCTTCCGGAAGTAGTCCCGGTAATTCCGCTGACGGCGGC
GGAGGATCGGGTGGTAGTGGTGGTTCAGGAGGAGGATCGACCCAAGGAGGATCCATGGTGAGCGTGATCA
AGCCCGAGATGAAGATCAAGCTGTGCATGAGGGGCACCGTGAACGGCCACAACTTCGTGATCGAGGGCGA
GGGCAAGGGCAACCCCTACGAGGGCACCCAGATCCTGGACCTGAACGTGACCGAGGGCGCCCCCCTGCCC
TTCGCCTACGACATCCTGACCACCGTGTTCCAGTACGGCAACAGGGCCTTCACCAAGTACCCCGCCGACA
TCCAGGACTACTTCAAGCAGACCTTCCCCGAGGGCTACCACTGGGAGAGGAGCATGACCTACGAGGACCA
GGGCATCTGCACCGCCACCAGCAACATCAGCATGAGGGGCGACTGCTTCTTCTACGACATCAGGTTCGAC
GGCGTGAACTTCCCCCCCAACGGCCCCGTGATGCAGAAGAAGACCCTGAAGTGGGAGCCCAGCACCGAGA
AGATGTACGTGAGGGACGGCGTGCTGAAGGGCGACGTGAACATGGCCCTGCTGCTGGAGGGCGGCGGCCA
CTACAGGTGCGACTTCAAGACCACCTACAAGGCCAAGAAGGACGTGAGGCTGCCCGACTACCACTTCGTG
GACCACAGGATCGAGATCCTGAAGCACGACAAGGACTACAACAAGGTGAAGCTGTACGAGAACGCCGTGG
CCAGGTACAGCATGCTGCCCAGCCAGGCCAAGGGTACCGGAACTGCAGCAGAGAATTCGGGAAACTCGAG
AACAAAGCTTATGATCCTCTGGCATGAGATGTGGCATGAAGGCCTGGAAGAGGCATCTCGTTTGTACTTT
GGGGAAAGGAACGTGAAAGGCATGTTTGAGGTGCTGGAGCCCTTGCATGCTATGATGGAACGGGGCCCCC
AGACTCTGAAGGAAACATCCTTTAATCAGGCCTATGGTCGAGATTTAATGGAGGCCCAAGAGTGGTGCAG
GAAGTACATGAAATCAGGGAATGTCAAGGACCTCCTCCAAGCCTGGGACCTCTATTATCATGTGTTCCGA
CGAATCTCAAAGCCCGGGGGTAGTGGTGCTGGCTCTGGTGCTGGTAGTGGCGCTGGTGAAAACCTGTATT
TTCAGCTGGTCGATGGTGGTAGTGCTGGTGGTTCCGGAGGTGTCGACATGGTGTCTAAGGGCGAAGAGCT
GATTAAGGAGAACATGCACATGAAGCTGTACATGGAGGGCACCGTGGACAACCATCACTTCAAGTGCACA
TCCGAGGGCGAAGGCAAGCCCTACGAGGGCACCCAGACCATGAGAATCAAGGTGGTCGAGGGCGGCCCTC
TCCCCTTCGCCTTCGACATCCTGGCTACTAGCTTCCTCTACGGCAGCAAGACCTTCATCAACCACACCCA
GGGCATCCCCGACTTCTTCAAGCAGTCCTTCCCTGAGGGCTTCACATGGGAGAGAGTCACCACATACGAA

GACGGGGGCGTGCTGACCGCTACCCAGGACACCAGCCTCCAGGACGGCTGCCTCATCTACAACGTCAAGA
TCAGAGGGGTGAACTTCACATCCAACGGCCCTGTGATGCAGAAGAAAACACTCGGCTGGGAGGCCTTCAC
CGAGACGCTGTACCCCGCTGACGGCGGCCTGGAAGGCAGAAACGACATGGCCCTGAAGCTCGTGGGCGGG
AGCCATCTGATCGCAAACGCCAAGACCACATATAGATCCAAGAAACCCGCTAAGAACCTCAAGATGCCTG
GCGTCTACTATGTGGACTACAGACTGGAAAGAATCAAGGAGGCCAACAACGAGACCTACGTCGAGCAGCA
CGAGGTGGCAGTGGCCAGATACTGCGACCTCCCTAGCAAACTGGGGCACAAGCTTAATAGATCCGGCTCT
TACCCATACGATGTTCCAGATTACGCTAGATCTCGAGGGCCCATGAAAATGGGCATGACTGAGGACGACA
AGAGAAGCTGCTGCTTGTTAGAGATTCAGGAGACCGAGGCCAAGTACTACCGCACCCTGGAGGACATTGA
GAAGAACTACATGGGTCCCTTGCGGCTGGTGCTGAGCCCGGCGGATATGGCTGCTGTCTTCATCAACCTG
GAGGACCTCATCAAGGTGCATCACAGCTTTCTGCGAGCCATCGATGTGTCCATGATGGCTGGTGGCAGTA
CCCTGGCTAAGGTCTTTCTGGAGTTTAAGGAAAGGCTCCTGATCTATGGAGAGTACTGTAGCCACATGGA
ACACGCTCAGAGTACACTGAACCAGCTCCTCGCCAGCCGAGAGGACTTCAGGCAGAAAGTGGAGGAGTGC
ACACTCAGGGTTCAGGATGGCAAGTTCAAGCTGCAAGACCTGCTGGTGGTGCCCATGCAACGGGTGCTGA
AGTACCACCTGCTGCTCAAGGAGCTCCTGAGCCATTCTGCAGACCGACCAGAAAGACAACAGCTCAAAGA
AGCCCTGGAAGCCATGCAGGACTTGGCCATGTACATCAATGAAGTGAAGCGGGACAAGGAGACCTTGAAG
AAGATTAGCGAGTTCCAGTGCTCCATAGAAAACCTGCAAGTGAAGCTGGAGGAATTTGGGAGGCCAAAGA
TTGACGGGGAGCTTAAAGTCCGGTCCATAGTCAACCACACCAAGCAAGACAGGTACCTGTTCCTATTTGA
CAAGGTGGTCATCGTGTGCAAGAGGAAGGGCTACAGCTATGAGCTGAAGGAGGTCATTGAGCTGCTCTTC
CACAAGATGACCGATGACCCGATGCACAACAAGGACATCAAGAAGTGGTCCTATGGCTTCTACCTGATTC
ACCTCCAAGGAAAGCAAGGCTTTCAGTTCTTCTGCAAGACGGAAGACATGAAGCGGAAGTGGATGGAGCA
GTTCGAGATGGCCATGTCAAACATCAAGCCAGATAAGGCCAATGCCAACCATCATAGCTTCCAGATGTAC
ACATTCGACAAGACTACCAACTGCAAAGCCTGCAAGATGTTTCTCAGGGGTACCTTCTACCAGGGATACC
TGTGTACCAGATGTGGCGTCGGGGCACACAAGGAATGCCTGGAGGTGATCCCCCCCTGCAAGTAA

**Annotation**
PB1: PB1 domain from human p62 (residues 1-102)
AG: green fluorescent protein Azami-Green
FRB: FRB domain from human mammalian target of rapamycin (mTOR) (residues 2021-2113 with a T2098L mutation)
$TEV_{cs}$: TEV protease cleavage sequence (ENLYFQ/L)
mTagBFP2: blue fluorescent protein mTagBFP2
HA: HA epitope tag
$Vav2_{cat}$: DH-PH-CR domain from mouse Vav2 (residues 183-563)

**s**

**pPBbsr2-Lifeact-tdiRFP670**

>Amino acid sequence
MGVADLIKKFESISKEEGDPPVATMARKVDLTSCDREPIHIPGSIQPCGCLLACDAQAVRITRITENAGA
FFGRETPRVGELLADYFGETEAHALRNALAQSSDPKRPALIFGWRDGLTGRTFDISLHRHDGTSIIEFEP
AAAEQADNPLRLTRQIIARTKELKSLEEMAARVPRYLQAMLGYHRVMLYRFADDGSGMVIGEAKRSDLES
FLGQHFPASLVPQQARLLYLKNAIRVVSDSRGISSRIVPEHDASGAALDLSFAHLRSISPCHLEFLRNMG
VSASMSLSIIIDGTLWGLIICHHYEPRAVPMAQRVAAEMFADFLSLHFTAAHHQRGHATGSTGSGSAEGG
TASSEDNMARKVDLTSCDREPIHIPGSIQPCGCLLACDAQAVRITRITENAGAFFGRETPRVGELLADYF
GETEAHALRNALAQSSDPKRPALIFGWRDGLTGRTFDISLHRHDGTSIIEFEPAAAEQADNPLRLTRQII
ARTKELKSLEEMAARVPRYLQAMLGYHRVMLYRFADDGSGMVIGEAKRSDLESFLGQHFPASLVPQQARL
LYLKNAIRVVSDSRGISSRIVPEHDASGAALDLSFAHLRSISPCHLEFLRNMGVSASMSLSIIIDGTLWG
LIICHHYEPRAVPMAQRVAAEMFADFLSLHFTAAHHQRMYK*[EMCV_IRES]MLYEDNKHHVGAAIRTK
TGEIISAVHIEAYIGRVTVCAEAIAIGSAVSNGQKDFDTIVAVRHPYSDEVDRSIRVVSPCGMCRELISD
YAPDCFVLIEMNGKLVKTTIEELIPLKYTRN*

>DNA sequence
ATGGGCGTGGCCGACTTGATCAAGAAGTTCGAGTCCATCTCCAAGGAGGAGGGGGATCCACCGGTCGCCA
CCATGGCTCGCAAGGTGGACCTGACCAGCTGCGACCGCGAGCCCATCCACATCCCCGGCAGCATCCAGCC
CTGCGGCTGCCTGCTGGCCTGCGACGCCCAGGCCGTGCGCATCACCCGCATCACCGAGAACGCCGGCGCC
TTCTTCGGCCGCGAGACCCCCCGCGTGGGCGAGCTGCTGGCCGACTACTTCGGCGAGACCGAGGCCCACG
CCCTGCGCAACGCCCTGGCCCAGAGCAGCGACCCCAAGCGCCCCGCCCTGATCTTCGGCTGGCGCGACGG
CCTGACCGGCCGCACCTTCGACATCAGCCTGCACCGCCACGACGGCACCAGCATCATCGAGTTCGAGCCC
GCCGCCGCCGAGCAGGCCGACAACCCCCTGCGCCTGACCCGCCAGATCATCGCCCGCACCAAGGAGCTGA
AGAGCCTGGAGGAGATGGCCGCCCGCGTGCCCCGCTACCTGCAGGCCATGCTGGGCTACCACCGCGTGAT
GCTGTACCGCTTCGCCGACGACGGCAGCGGCATGGTGATCGGCGAGGCCAAGCGCAGCGACCTGGAGAGC
TTCCTGGGCCAGCACTTCCCCGCCAGCCTGGTGCCCCAGCAGGCCCGCCTGCTGTACCTGAAGAACGCCA
TCCGCGTGGTGAGCGACAGCCGCGGCATCAGCAGCCGCATCGTGCCCGAGCACGACGCCAGCGGCGCCGC
CCTGGACCTGAGCTTCGCCCACCTGCGCAGCATCAGCCCCTGCCACCTGGAGTTCCTGCGCAACATGGGC
GTGAGCGCCAGCATGAGCCTGAGCATCATCATCGACGGCACCCTGTGGGGCCTGATCATCTGCCACCACT
ACGAGCCCCGCGCCGTGCCCATGGCCCAGCGCGTGGCCGCCGAGATGTTCGCCGACTTCCTGAGCCTGCA
CTTCACCGCCGCTCACCATCAGAGAGGACATGCTACTGGAAGCACTGGAAGCGGCAGCGCTGAGGGAGGC
ACAGCTTCTAGCGAAGATAATATGGCTCGCAAGGTGGACCTGACCAGCTGCGACCGCGAGCCCATCCACA
TCCCCGGCAGCATCCAGCCCTGCGGCTGCCTGCTGGCCTGCGACGCCCAGGCCGTGCGCATCACCCGCAT
CACCGAGAACGCCGGCGCCTTCTTCGGCCGCGAGACCCCCCGCGTGGGCGAGCTGCTGGCCGACTACTTC
GGCGAGACCGAGGCCCACGCCCTGCGCAACGCCCTGGCCCAGAGCAGCGACCCCAAGCGCCCCGCCCTGA
TCTTCGGCTGGCGCGACGGCCTGACCGGCCGCACCTTCGACATCAGCCTGCACCGCCACGACGGCACCAG
CATCATCGAGTTCGAGCCCGCCGCCGCCGAGCAGGCCGACAACCCCCTGCGCCTGACCCGCCAGATCATC
GCCCGCACCAAGGAGCTGAAGAGCCTGGAGGAGATGGCCGCCCGCGTGCCCCGCTACCTGCAGGCCATGC
TGGGCTACCACCGCGTGATGCTGTACCGCTTCGCCGACGACGGCAGCGGCATGGTGATCGGCGAGGCCAA
GCGCAGCGACCTGGAGAGCTTCCTGGGCCAGCACTTCCCCGCCAGCCTGGTGCCCCAGCAGGCCCGCCTG
CTGTACCTGAAGAACGCCATCCGCGTGGTGAGCGACAGCCGCGGCATCAGCAGCCGCATCGTGCCCGAGC
ACGACGCCAGCGGCGCCGCCCTGGACCTGAGCTTCGCCCACCTGCGCAGCATCAGCCCCTGCCACCTGGA
GTTCCTGCGCAACATGGGCGTGAGCGCCAGCATGAGCCTGAGCATCATCATCGACGGCACCCTGTGGGGC
CTGATCATCTGCCACCACTACGAGCCCCGCGCCGTGCCCATGGCCCAGCGCGTGGCCGCCGAGATGTTCG
CCGACTTCCTGAGCCTGCACTTCACCGCCGCTCACCATCAGAGAATGTACAAGTAAAGCGGCCGCTCTAG
AGTCGACGGGCCGCGGTAACAATTGTTAACTAACTTAAGCTAGCAACGGTTTCCCTCTAGCGGGATCAAT
TCCGCCCCCCCCCCCTAACGTTACTGGCCGAAGCCGCTTGGAATAAGGCCGGTGTGCGTTTGTCTATATG
TTATTTTCCACCATATTGCCGTCTTTTGGCAATGTGAGGGCCCGGAAACCTGGCCCTGTCTTCTTGACGA
GCATTCCTAGGGGTCTTTCCCCTCTCGCCAAAGGAATGCAAGGTCTGTTGAATGTCGTGAAGGAAGCAGT
TCCTCTGGAAGCTTCTTGAAGACAAACAACGTCTGTAGCGACCCTTTGCAGGCAGCGGAACCCCCCACCT
GGCGACAGGTGCCTCTGCGGCCAAAAGCCACGTGTATAAGATACACCTGCAAAGGCGGCACAACCCCAGT
GCCACGTTGTGAGTTGGATAGTTGTGGAAAGAGTCAAATGGCTCTCCTCAAGCGTATTCAACAAGGGGCT

GAAGGATGCCCAGAAGGTACCCCATTGTATGGGATCTGATCTGGGGCCTCGGTGCACATGCTTTACATGT
GTTTAGTCGAGGTTAAAAAACGTCTAGGCCCCCCGAACCACGGGGACGTGGTTTTCCTTTGAAAAACACG
ATAATACCATGGTCATGAAAACATTTAACATTTCTCAACAAGATCTAGAATTAGTAGAAGTAGCGACAGA
GAAGATTACAATGCTTTATGAGGATAATAAACATCATGTGGGAGCGGCAATTCGTACGAAAACAGGAGAA
ATCATTTCGGCAGTACATATTGAAGCGTATATAGGACGAGTAACTGTTTGTGCAGAAGCCATTGCGATTG
GTAGTGCAGTTTCGAATGGACAAAAGGATTTTGACACGATTGTAGCTGTTAGACACCCTTATTCTGACGA
AGTAGATAGAAGTATTCGAGTGGTAAGTCCTTGTGGTATGTGTAGGGAGTTGATTTCAGACTATGCACCA
GATTGTTTTGTGTTAATAGAAATGAATGGCAAGTTAGTCAAAACTACGATTGAAGAACTCATTCCACTCA
AATATACCCGAAATTAA

**Annotation**
Lifeact: actin-binding peptide Lifeact
iRFP670: near-infrared fluorescent protein iRFP670
ECMV IRES: internal ribosomal entry site from *Encephalomyocarditis* virus
Bsr: blasticidin S-deaminase

**t**

**pCMV-PB1-AG-FRB-TEVuncs-BFP-HA-Vav2**

>Amino acid sequence
MASLTVKAYLLGKEDAAREIRRFSFCCSPEPEAEAEAAAGPGPCERLLSRVAALFPALRPGGFQAHYRDE
DGDLVAFSSDEELTMAMSYVKDDIFRIYIKEKTGSGSGGSGAGGSAGSGAGGSAGSGAGGSAGSGAGGSA
GSGAGGSASGSSPGSGSGGSGAGGSAGSGAGGSAGSGAGGSAGSGAGGSAGSGAGGSASGSSPGNSADGG
GGSGGSGGSGGGSTQGGSMVSVIKPEMKIKLCMRGTVNGHNFVIEGEGKGNPYEGTQILDLNVTEGAPLP
FAYDILTTVFQYGNRAFTKYPADIQDYFKQTFPEGYHWERSMTYEDQGICTATSNISMRGDCFFYDIRFD
GVNFPPNGPVMQKKTLKWEPSTEKMYVRDGVLKGDVNMALLLEGGGHYRCDFKTTYKAKKDVRLPDYHFV
DHRIEILKHDKDYNKVKLYENAVARYSMLPSQAKGTGTAAENSGNSRTKLMILWHEMWHEGLEEASRLYF
GERNVKGMFEVLEPLHAMMERGPQTLKETSFNQAYGRDLMEAQEWCRKYMKSGNVKDLLQAWDLYYHVFR
RISKPGGSGAGSGAGSGAGNEYLQFLVDGGSAGGSGGVDMVSKGEELIKENMHMKLYMEGTVDNHHFKCT
SEGEGKPYEGTQTMRIKVVEGGPLPFAFDILATSFLYGSKTFINHTQGIPDFFKQSFPEGFTWERVTTYE
DGGVLTATQDTSLQDGCLIYNVKIRGVNFTSNGPVMQKKTLGWEAFTETLYPADGGLEGRNDMALKLVGG
SHLIANAKTTYRSKKPAKNLKMPGVYYVDYRLERIKEANNETYVEQHEVAVARYCDLPSKLGHKLNRSGS
YPYDVPDYARSRGPMKMGMTEDDKRSCCLLEIQETEAKYYRTLEDIEKNYMGPLRLVLSPADMAAVFINL
EDLIKVHHSFLRAIDVSMMAGGSTLAKVFLEFKERLLIYGEYCSHMEHAQSTLNQLLASREDFRQKVEEC
TLRVQDGKFKLQDLLVVPMQRVLKYHLLLKELLSHSADRPERQQLKEALEAMQDLAMYINEVKRDKETLK
KISEFQCSIENLQVKLEEFGRPKIDGELKVRSIVNHTKQDRYLFLFDKVVIVCKRKGYSYELKEVIELLF
HKMTDDPMHNKDIKKWSYGFYLIHLQGKQGFQFFCKTEDMKRKWMEQFEMAMSNIKPDKANANHHSFQMY
TFDKTTNCKACKMFLRGTFYQGYLCTRCGVGAHKECLEVIPPCK*

>DNA sequence
ATGGCGTCGCTCACCGTGAAGGCCTACCTTCTGGGCAAGGAGGACGCGGCGCGCGAGATTCGCCGCTTCA
GCTTCTGTTGCAGCCCCGAGCCTGAGGCGGAAGCCGAGGCTGCGGCGGGTCCGGGACCCTGCGAGCGGCT
GCTGAGCCGGGTGGCCGCCCTGTTCCCCGCGCTGCGGCCTGGCGGCTTCCAGGCGCACTACCGCGATGAG
GACGGGGACTTGGTTGCCTTTTCCAGTGACGAGGAATTGACAATGGCCATGTCCTACGTGAAGGATGACA
TCTTCCGAATCTACATTAAAGAGAAAACCGGTTCTGGGAGTGGCGGTAGTGGTGCTGGAGGCAGCGCAGG
CTCTGGAGCAGGCGGCAGTGCCGGAAGTGGCGCTGGAGGGTCTGCCGGATCTGGAGCCGGTGGTTCTGCC
GGCTCCGGCGCTGGTGGGAGCGCTTCCGGAAGTAGTCCCGGTTCTGGGAGTGGCGGTAGTGGTGCTGGAG
GCAGCGCAGGCTCTGGAGCAGGCGGCAGTGCCGGAAGTGGCGCTGGAGGGTCTGCCGGATCTGGAGCCGG
TGGTTCTGCCGGCTCCGGCGCTGGTGGGAGCGCTTCCGGAAGTAGTCCCGGTAATTCCGCTGACGGCGGC
GGAGGATCGGGTGGTAGTGGTGGTTCAGGAGGAGGATCGACCCAAGGAGGATCCATGGTGAGCGTGATCA
AGCCCGAGATGAAGATCAAGCTGTGCATGAGGGGCACCGTGAACGGCCACAACTTCGTGATCGAGGGCGA
GGGCAAGGGCAACCCCTACGAGGGCACCCAGATCCTGGACCTGAACGTGACCGAGGGCGCCCCCCTGCCC
TTCGCCTACGACATCCTGACCACCGTGTTCCAGTACGGCAACAGGGCCTTCACCAAGTACCCCGCCGACA
TCCAGGACTACTTCAAGCAGACCTTCCCCGAGGGCTACCACTGGGAGAGGAGCATGACCTACGAGGACCA
GGGCATCTGCACCGCCACCAGCAACATCAGCATGAGGGGCGACTGCTTCTTCTACGACATCAGGTTCGAC
GGCGTGAACTTCCCCCCCAACGGCCCCGTGATGCAGAAGAAGACCCTGAAGTGGGAGCCCAGCACCGAGA
AGATGTACGTGAGGGACGGCGTGCTGAAGGGCGACGTGAACATGGCCCTGCTGCTGGAGGGCGGCGGCCA
CTACAGGTGCGACTTCAAGACCACCTACAAGGCCAAGAAGGACGTGAGGCTGCCCGACTACCACTTCGTG
GACCACAGGATCGAGATCCTGAAGCACGACAAGGACTACAACAAGGTGAAGCTGTACGAGAACGCCGTGG
CCAGGTACAGCATGCTGCCCAGCCAGGCCAAGGGTACCGGAACTGCAGCAGAGAATTCGGGAAACTCGAG
AACAAAGCTTATGATCCTCTGGCATGAGATGTGGCATGAAGGCCTGGAAGAGGCATCTCGTTTGTACTTT
GGGGAAAGGAACGTGAAAGGCATGTTTGAGGTGCTGGAGCCCTTGCATGCTATGATGGAACGGGGCCCCC
AGACTCTGAAGGAAACATCCTTTAATCAGGCCTATGGTCGAGATTTAATGGAGGCCCAAGAGTGGTGCAG
GAAGTACATGAAATCAGGGAATGTCAAGGACCTCCTCCAAGCCTGGGACCTCTATTATCATGTGTTCCGA
CGAATCTCAAAGCCCGGGGGTAGTGGTGCTGGCTCTGGTGCTGGTAGTGGCGCTGGTAACGAATATCTGC
AGTTTCTGGTCGATGGTGGTAGTGCTGGTGGTTCCGGAGGTGTCGACATGGTGTCTAAGGGCGAAGAGCT
GATTAAGGAGAACATGCACATGAAGCTGTACATGGAGGGCACCGTGGACAACCATCACTTCAAGTGCACA
TCCGAGGGCGAAGGCAAGCCCTACGAGGGCACCCAGACCATGAGAATCAAGGTGGTCGAGGGCGGCCCTC
TCCCCTTCGCCTTCGACATCCTGGCTACTAGCTTCCTCTACGGCAGCAAGACCTTCATCAACCACACCCA
GGGCATCCCCGACTTCTTCAAGCAGTCCTTCCCTGAGGGCTTCACATGGGAGAGAGTCACCACATACGAA

GACGGGGGCGTGCTGACCGCTACCCAGGACACCAGCCTCCAGGACGGCTGCCTCATCTACAACGTCAAGA
TCAGAGGGGTGAACTTCACATCCAACGGCCCTGTGATGCAGAAGAAAACACTCGGCTGGGAGGCCTTCAC
CGAGACGCTGTACCCCGCTGACGGCGGCCTGGAAGGCAGAAACGACATGGCCCTGAAGCTCGTGGGCGGG
AGCCATCTGATCGCAAACGCCAAGACCACATATAGATCCAAGAAACCCGCTAAGAACCTCAAGATGCCTG
GCGTCTACTATGTGGACTACAGACTGGAAAGAATCAAGGAGGCCAACAACGAGACCTACGTCGAGCAGCA
CGAGGTGGCAGTGGCCAGATACTGCGACCTCCCTAGCAAACTGGGGCACAAGCTTAATAGATCCGGCTCT
TACCCATACGATGTTCCAGATTACGCTAGATCTCGAGGGCCCATGAAAATGGGCATGACTGAGGACGACA
AGAGAAGCTGCTGCTTGTTAGAGATTCAGGAGACCGAGGCCAAGTACTACCGCACCCTGGAGGACATTGA
GAAGAACTACATGGGTCCCTTGCGGCTGGTGCTGAGCCCGGCGGATATGGCTGCTGTCTTCATCAACCTG
GAGGACCTCATCAAGGTGCATCACAGCTTTCTGCGAGCCATCGATGTGTCCATGATGGCTGGTGGCAGTA
CCCTGGCTAAGGTCTTTCTGGAGTTTAAGGAAAGGCTCCTGATCTATGGAGAGTACTGTAGCCACATGGA
ACACGCTCAGAGTACACTGAACCAGCTCCTCGCCAGCCGAGAGGACTTCAGGCAGAAAGTGGAGGAGTGC
ACACTCAGGGTTCAGGATGGCAAGTTCAAGCTGCAAGACCTGCTGGTGGTGCCCATGCAACGGGTGCTGA
AGTACCACCTGCTGCTCAAGGAGCTCCTGAGCCATTCTGCAGACCGACCAGAAAGACAACAGCTCAAAGA
AGCCCTGGAAGCCATGCAGGACTTGGCCATGTACATCAATGAAGTGAAGCGGGACAAGGAGACCTTGAAG
AAGATTAGCGAGTTCCAGTGCTCCATAGAAAACCTGCAAGTGAAGCTGGAGGAATTTGGAGGCCAAAGA
TTGACGGGGAGCTTAAAGTCCGGTCCATAGTCAACCACACCAAGCAAGACAGGTACCTGTTCCTATTTGA
CAAGGTGGTCATCGTGTGCAAGAGGAAGGGCTACAGCTATGAGCTGAAGGAGGTCATTGAGCTGCTCTTC
CACAAGATGACCGATGACCCGATGCACAACAAGGACATCAAGAAGTGGTCCTATGGCTTCTACCTGATTC
ACCTCCAAGGAAAGCAAGGCTTTCAGTTCTTCTGCAAGACGGAAGACATGAAGCGGAAGTGGATGGAGCA
GTTCGAGATGGCCATGTCAAACATCAAGCCAGATAAGGCCAATGCCAACCATCATAGCTTCCAGATGTAC
ACATTCGACAAGACTACCAACTGCAAAGCCTGCAAGATGTTTCTCAGGGGTACCTTCTACCAGGGATACC
TGTGTACCAGATGTGGCGTCGGGGCACACAAGGAATGCCTGGAGGTGATCCCCCCCTGCAAGTAA

**Annotation**

PB1: PB1 domain  from human p62 (residues 1-102)
AG: green fluorescent protein Azami-Green
FRB: FRB domain from human mammalian target of rapamycin (mTOR) (residues 2021-2113 with a T2098L mutation)
$TEV_{uncs}$: TEV protease uncleavable sequence (NEYLQFL)
mTagBFP2: blue fluorescent protein mTagBFP2
HA: HA epitope tag
$Vav2_{cat}$: DH-PH-CR domain from mouse Vav2 (residues 183-563)

**u**

**pCMV-tdiRFP670-FKBP-TEVp**

>Amino acid sequence
MARKVDLTSCDREPIHIPGSIQPCGCLLACDAQAVRITRITENAGAFFGRETPRVGELLADYFGETEAHA
LRNALAQSSDPKRPALIFGWRDGLTGRTFDISLHRHDGTSIIEFEPAAAEQADNPLRLTRQIIARTKELK
SLEEMAARVPRYLQAMLGYHRVMLYRFADDGSGMVIGEAKRSDLESFLGQHFPASLVPQQARLLYLKNAI
RVVSDSRGISSRIVPEHDASGAALDLSFAHLRSISPCHLEFLRNMGVSASMSLSIIIDGTLWGLIICHHY
EPRAVPMAQRVAAEMFADFLSLHFTAAHHQRGHATGSTGSGSAEGGTASSEDNMARKVDLTSCDREPIHI
PGSIQPCGCLLACDAQAVRITRITENAGAFFGRETPRVGELLADYFGETEAHALRNALAQSSDPKRPALI
FGWRDGLTGRTFDISLHRHDGTSIIEFEPAAAEQADNPLRLTRQIIARTKELKSLEEMAARVPRYLQAML
GYHRVMLYRFADDGSGMVIGEAKRSDLESFLGQHFPASLVPQQARLLYLKNAIRVVSDSRGISSRIVPEH
DASGAALDLSFAHLRSISPCHLEFLRNMGVSASMSLSIIIDGTLWGLIICHHYEPRAVPMAQRVAAEMFA
DFLSLHFTAAHHQRMYKSGLRSRAMGVQVETISPGDGRTFPKRGQTCVVHYTGMLEDGKKFDSSRDRNKP
FKFMLGKQEVIRGWEEGVAQMSVGQRAKLTISPDYAYGATGHPGIIPPHATLVFDVELLKLEEFCSRRYR
MGESLFKGPRDYNPISSTICHLTNESDGHTTSLYGIGFGPFIITNKHLFRRNNGTLLVQSLHGVFKVKNT
TTLQQHLIDGRDMIIIRMPKDFPPFPQKLKFREPQREERICLVTTNFQTKSMSSMVSDTSCTFPSSDGIF
WKHWIQTKDGQCGSPLVSTRDGFIVGIHSASNFTNTNNYFTSVPKNFMELLTNQEAQQWVSGWRLNADSV
LWGGHKVFMVGSTGSR*

>DNA sequence
ATGGCTCGCAAGGTGGACCTGACCAGCTGCGACCGCGAGCCCATCCACATCCCCGGCAGCATCCAGCCCT
GCGGCTGCCTGCTGGCCTGCGACGCCCAGGCCGTGCGCATCACCCGCATCACCGAGAACGCCGGCGCCTT
CTTCGGCCGCGAGACCCCCCGCGTGGGCGAGCTGCTGGCCGACTACTTCGGCGAGACCGAGGCCCACGCC
CTGCGCAACGCCCTGGCCCAGAGCAGCGACCCCAAGCGCCCCGCCCTGATCTTCGGCTGGCGCGACGGCC
TGACCGGCCGCACCTTCGACATCAGCCTGCACCGCCACGACGGCACCAGCATCATCGAGTTCGAGCCCGC
CGCCGCCGAGCAGGCCGACAACCCCCTGCGCCTGACCCGCCAGATCATCGCCCGCACCAAGGAGCTGAAG
AGCCTGGAGGAGATGGCCGCCCGCGTGCCCCGCTACCTGCAGGCCATGCTGGGCTACCACCGCGTGATGC
TGTACCGCTTCGCCGACGACGGCAGCGGCATGGTGATCGGCGAGGCCAAGCGCAGCGACCTGGAGAGCTT
CCTGGGCCAGCACTTCCCCGCCAGCCTGGTGCCCCAGCAGGCCCGCCTGCTGTACCTGAAGAACGCCATC
CGCGTGGTGAGCGACAGCCGCGGCATCAGCAGCCGCATCGTGCCCGAGCACGACGCCAGCGGCGCCGCCC
TGGACCTGAGCTTCGCCCACCTGCGCAGCATCAGCCCCTGCCACCTGGAGTTCCTGCGCAACATGGGCGT
GAGCGCCAGCATGAGCCTGAGCATCATCATCGACGGCACCCTGTGGGGCCTGATCATCTGCCACCACTAC
GAGCCCCGCGCCGTGCCCATGGCCCAGCGCGTGGCCGCCGAGATGTTCGCCGACTTCCTGAGCCTGCACT
TCACCGCCGCTCACCATCAGAGAGGACATGCTACTGGAAGCACTGGAAGCGGCAGCGCTGAGGGAGGCAC
AGCTTCTAGCGAAGATAATATGGCTCGCAAGGTGGACCTGACCAGCTGCGACCGCGAGCCCATCCACATC
CCCGGCAGCATCCAGCCCTGCGGCTGCCTGCTGGCCTGCGACGCCCAGGCCGTGCGCATCACCCGCATCA
CCGAGAACGCCGGCGCCTTCTTCGGCCGCGAGACCCCCCGCGTGGGCGAGCTGCTGGCCGACTACTTCGG
CGAGACCGAGGCCCACGCCCTGCGCAACGCCCTGGCCCAGAGCAGCGACCCCAAGCGCCCCGCCCTGATC
TTCGGCTGGCGCGACGGCCTGACCGGCCGCACCTTCGACATCAGCCTGCACCGCCACGACGGCACCAGCA
TCATCGAGTTCGAGCCCGCCGCCGCCGAGCAGGCCGACAACCCCCTGCGCCTGACCCGCCAGATCATCGC
CCGCACCAAGGAGCTGAAGAGCCTGGAGGAGATGGCCGCCCGCGTGCCCCGCTACCTGCAGGCCATGCTG
GGCTACCACCGCGTGATGCTGTACCGCTTCGCCGACGACGGCAGCGGCATGGTGATCGGCGAGGCCAAGC
GCAGCGACCTGGAGAGCTTCCTGGGCCAGCACTTCCCCGCCAGCCTGGTGCCCCAGCAGGCCCGCCTGCT
GTACCTGAAGAACGCCATCCGCGTGGTGAGCGACAGCCGCGGCATCAGCAGCCGCATCGTGCCCGAGCAC
GACGCCAGCGGCGCCGCCCTGGACCTGAGCTTCGCCCACCTGCGCAGCATCAGCCCCTGCCACCTGGAGT
TCCTGCGCAACATGGGCGTGAGCGCCAGCATGAGCCTGAGCATCATCATCGACGGCACCCTGTGGGGCCT
GATCATCTGCCACCACTACGAGCCCCGCGCCGTGCCCATGGCCCAGCGCGTGGCCGCCGAGATGTTCGCC
GACTTCCTGAGCCTGCACTTCACCGCCGCTCACCATCAGAGAATGTACAAGTCCGGACTCAGATCTCGAG
CTATGGGAGTGCAGGTGGAAACCATCTCCCCAGGAGACGGGCGCACCTTCCCCAAGCGCGGCCAGACCTG
CGTGGTGCACTACACCGGGATGCTTGAAGATGGAAAGAAATTTGATTCCTCCCGGGACAGAAACAAGCCC
TTTAAGTTTATGCTAGGCAAGCAGGAGGTGATCCGAGGCTGGGAAGAAGGGGTTGCCCAGATGAGTGTGG
GTCAGAGAGCCAAACTGACTATATCTCCAGATTATGCCTATGGTGCCACTGGGCACCCAGGCATCATCCC
ACCACATGCCACTCTCGTCTTCGATGTGGAGCTTCTAAAACTGGAAGAATTCTGCAGTCGACGGTACCGC

ATGGGCGAGAGCCTTTTCAAGGGCCCGAGGGACTACAACCCGATCTCCAGCACCATCTGTCACCTGACCA
ACGAGAGCGACGGTCACACCACTAGTCTGTACGGCATCGGCTTCGGCCCCTTCATCATCACCAACAAGCA
TCTGTTCAGGAGGAATAACGGCACACTGCTGGTGCAAAGCCTGCACGGCGTGTTCAAAGTGAAGAACACA
ACCACCCTGCAACAGCACCTGATCGACGGCAGGGACATGATTATCATCAGGATGCCCAAGGACTTCCCCC
CCTTTCCCCAGAAACTGAAGTTCAGGGAGCCACAAAGGGAGGAGCGAATCTGCCTGGTGACCACCAACTT
CCAGACCAAGTCCATGAGCAGCATGGTCTCTGATACCAGCTGCACCTTCCCCAGCAGCGACGGCATCTTC
TGGAAGCACTGGATTCAGACGAAGGATGGCCAATGCGGCAGCCCATTGGTGAGCACTAGGGACGGCTTCA
TCGTGGGCATCCACAGCGCCAGCAATTTTACCAATACCAACAACTACTTCACGAGCGTGCCGAAAAACTT
CATGGAGCTGTTGACCAATCAAGAGGCGCAGCAGTGGGTGAGCGGCTGGAGGCTGAACGCCGACAGCGTT
CTTTGGGGCGGACATAAGGTGTTCATGGTCGGATCCACCGGATCTAGATAA

**Annotation**

iRFP670: near-infrared fluorescent protein iRFP670

FKBP: FKBP12

TEVp: TEV protease lacking the C-terminal residues 220-242

**v**

**pCMV-PB1-AG-FRB-TEVcs-BFP-HA-SOS**

>Amino acid sequence
MASLTVKAYLLGKEDAAREIRRFSFCCSPEPEAEAEAAAGPGPCERLLSRVAALFPALRPGGFQAHYRDE
DGDLVAFSSDEELTMAMSYVKDDIFRIYIKEKTGSGSGGSGAGGSAGSGAGGSAGSGAGGSAGSGAGGSA
GSGAGGSASGSSPGSGSGGSGAGGSAGSGAGGSAGSGAGGSAGSGAGGSAGSGAGGSASGSSPGNSADGG
GGSGGSGGSGGGSTQGGSMVSVIKPEMKIKLCMRGTVNGHNFVIEGEGKGNPYEGTQILDLNVTEGAPLP
FAYDILTTVFQYGNRAFTKYPADIQDYFKQTFPEGYHWERSMTYEDQGICTATSNISMRGDCFFYDIRFD
GVNFPPNGPVMQKKTLKWEPSTEKMYVRDGVLKGDVNMALLLEGGGHYRCDFKTTYKAKKDVRLPDYHFV
DHRIEILKHDKDYNKVKLYENAVARYSMLPSQAKGTGTAAENSGNSRTKLMILWHEMWHEGLEEASRLYF
GERNVKGMFEVLEPLHAMMERGPQTLKETSFNQAYGRDLMEAQEWCRKYMKSGNVKDLLQAWDLYYHVFR
RISKPGGSGAGSGAGSGAGENLYFQLVDGGSAGGSGGVDMVSKGEELIKENMHMKLYMEGTVDNHHFKCT
SEGEGKPYEGTQTMRIKVVEGGPLPFAFDILATSFLYGSKTFINHTQGIPDFFKQSFPEGFTWERVTTYE
DGGVLTATQDTSLQDGCLIYNVKIRGVNFTSNGPVMQKKTLGWEAFTETLYPADGGLEGRNDMALKLVGG
SHLIANAKTTYRSKKPAKNLKMPGVYYVDYRLERIKEANNETYVEQHEVAVARYCDLPSKLGHKLNRSGS
YPYDVPDYARSQMRLPSADVYRFAEPDSEENIIFEENMQPKAGIPIIKAGTVIKLIERLTYHMYADPNFV
RTFLTTYRSFCKPQELLSLIIERFEIPEPEPTEADRIAIENGDQPLSAELKRFRKEYIQPVQLRVLNVCR
HWVEHHFYDFERDAYLLQRMEEFIGTVRGKAMKKWVESITKIIQRKKIARDNGPGHNITFQSSPPTVEWH
ISRPGHIETFDLLTLHPIEIARQLTLLESDLYRAVQPSELVGSVWTKEDKEINSPNLLKMIRHTTNLTLW
FEKCIVETENLEERVAVVSRIIEILQVFQELNNFNGVLEVVSAMNSSPVYRLDHTFEQIPSRQKKILEEA
HELSEDHYKKYLAKLRSINPPCVPFFGIYLTNILKTEEGNPEVLKRHGKELINFSKRRKVAEITGEIQQY
QNQPYCLRVESDIKRFFENLNPMGNSMEKEFTDYLFNKSLEIEPRNPKPLPRFPKKYSYPLKSPGVRPSN
PRPGT*

>DNA sequence
ATGGCGTCGCTCACCGTGAAGGCCTACCTTCTGGGCAAGGAGGACGCGGCGCGCGAGATTCGCCGCTTCA
GCTTCTGTTGCAGCCCCGAGCCTGAGGCGGAAGCCGAGGCTGCGGCGGGTCCGGGACCCTGCGAGCGGCT
GCTGAGCCGGGTGGCCGCCCTGTTCCCCGCGCTGCGGCCTGGCGGCTTCCAGGCGCACTACCGCGATGAG
GACGGGGACTTGGTTGCCTTTTCCAGTGACGAGGAATTGACAATGGCCATGTCCTACGTGAAGGATGACA
TCTTCCGAATCTACATTAAAGAGAAAACCGGTTCTGGGAGTGGCGGTAGTGGTGCTGGAGGCAGCGCAGG
CTCTGGAGCAGGCGGCAGTGCCGGAAGTGGCGCTGGAGGGTCTGCCGGATCTGGAGCCGGTGGTTCTGCC
GGCTCCGGCGCTGGTGGGAGCGCTTCCGGAAGTAGTCCCGGTTCTGGGAGTGGCGGTAGTGGTGCTGGAG
GCAGCGCAGGCTCTGGAGCAGGCGGCAGTGCCGGAAGTGGCGCTGGAGGGTCTGCCGGATCTGGAGCCGG
TGGTTCTGCCGGCTCCGGCGCTGGTGGGAGCGCTTCCGGAAGTAGTCCCGGTAATTCCGCTGACGGCGGC
GGAGGATCGGGTGGTAGTGGTGGTTCAGGAGGAGGATCGACCCAAGGAGGATCCATGGTGAGCGTGATCA
AGCCCGAGATGAAGATCAAGCTGTGCATGAGGGGCACCGTGAACGGCCACAACTTCGTGATCGAGGGCGA
GGGCAAGGGCAACCCCTACGAGGGCACCCAGATCCTGGACCTGAACGTGACCGAGGGCGCCCCCCTGCCC
TTCGCCTACGACATCCTGACCACCGTGTTCCAGTACGGCAACAGGGCCTTCACCAAGTACCCCGCCGACA
TCCAGGACTACTTCAAGCAGACCTTCCCCGAGGGCTACCACTGGGAGAGGAGCATGACCTACGAGGACCA
GGGCATCTGCACCGCCACCAGCAACATCAGCATGAGGGGCGACTGCTTCTTCTACGACATCAGGTTCGAC
GGCGTGAACTTCCCCCCCAACGGCCCCGTGATGCAGAAGAAGACCCTGAAGTGGGAGCCCAGCACCGAGA
AGATGTACGTGAGGGACGGCGTGCTGAAGGGCGACGTGAACATGGCCCTGCTGCTGGAGGGCGGCGGCCA
CTACAGGTGCGACTTCAAGACCACCTACAAGGCCAAGAAGGACGTGAGGCTGCCCGACTACCACTTCGTG
GACCACAGGATCGAGATCCTGAAGCACGACAAGGACTACAACAAGGTGAAGCTGTACGAGAACGCCGTGG
CCAGGTACAGCATGCTGCCCAGCCAGGCCAAGGGTACCGGAACTGCAGCAGAGAATTCGGGAAACTCGAG
AACAAAGCTTATGATCCTCTGGCATGAGATGTGGCATGAAGGCCTGGAAGAGGCATCTCGTTTGTACTTT
GGGGAAAGGAACGTGAAAGGCATGTTTGAGGTGCTGGAGCCCTTGCATGCTATGATGGAACGGGGCCCCC
AGACTCTGAAGGAAACATCCTTTAATCAGGCCTATGGTCGAGATTTAATGGAGGCCCAAGAGTGGTGCAG
GAAGTACATGAAATCAGGGAATGTCAAGGACCTCCTCCAAGCCTGGGACCTCTATTATCATGTGTTCCGA
CGAATCTCAAAGCCCGGGGGTAGTGGTGCTGGCTCTGGTGCTGGTAGTGGCGCTGGTGAAAACCTGTATT
TTCAGCTGGTCGATGGTGGTAGTGCTGGTGGTTCCGGAGGTGTCGACATGGTGTCTAAGGGCGAAGAGCT
GATTAAGGAGAACATGCACATGAAGCTGTACATGGAGGGCACCGTGGACAACCATCACTTCAAGTGCACA
TCCGAGGGCGAAGGCAAGCCCTACGAGGGCACCCAGACCATGAGAATCAAGGTGGTCGAGGGCGGCCCTC

```
TCCCCTTCGCCTTCGACATCCTGGCTACTAGCTTCCTCTACGGCAGCAAGACCTTCATCAACCACACCCA
GGGCATCCCCGACTTCTTCAAGCAGTCCTTCCCTGAGGGCTTCACATGGGAGAGAGTCACCACATACGAA
GACGGGGGCGTGCTGACCGCTACCCAGGACACCAGCCTCCAGGACGGCTGCCTCATCTACAACGTCAAGA
TCAGAGGGGTGAACTTCACATCCAACGGCCCTGTGATGCAGAAGAAAACACTCGGCTGGGAGGCCTTCAC
CGAGACGCTGTACCCCGCTGACGGCGGCCTGGAAGGCAGAAACGACATGGCCCTGAAGCTCGTGGGCGGG
AGCCATCTGATCGCAAACGCCAAGACCACATATAGATCCAAGAAACCCGCTAAGAACCTCAAGATGCCTG
GCGTCTACTATGTGGACTACAGACTGGAAAGAATCAAGGAGGCCAACAACGAGACCTACGTCGAGCAGCA
CGAGGTGGCAGTGGCCAGATACTGCGACCTCCCTAGCAAACTGGGGCACAAGCTTAATAGATCCGGCTCT
TACCCATACGATGTTCCAGATTACGCTAGATCTCAGATGAGGCTGCCTAGTGCTGATGTTTATAGATTTG
CAGAGCCTGACTCTGAAGAGAATATTATATTTGAAGAGAACATGCAGCCCAAGGCTGGAATTCCAATTAT
CAAAGCAGGAACTGTTATTAAACTTATAGAGAGGCTTACGTACCATATGTACGCAGATCCCAATTTTGTT
CGGACATTTCTTACAACATACAGATCCTTTTGCAAACCTCAAGAACTACTGAGTCTTATAATAGAAAGGT
TTGAAATTCCAGAGCCTGAGCCAACAGAAGCTGATCGCATAGCTATAGAGAATGGAGATCAACCCTTGAG
TGCAGAACTGAAAAGATTTAGAAAAGAATATATACAGCCTGTGCAACTGCGAGTATTAAATGTATGTCGG
CACTGGGTAGAGCACCACTTCTATGATTTTGAAAGAGATGCATATCTTTTGCAACGAATGGAAGAATTTA
TTGGAACAGTAAGAGGTAAAGCAATGAAAAAATGGGTTGAATCCATCACTAAAATAATCCAAAGGAAAAA
AATTGCAAGAGACAATGGACCAGGTCATAATATTACATTTCAGAGTTCACCTCCCACAGTTGAGTGGCAT
ATAAGCAGACCTGGGCACATAGAGACTTTTGACCTGCTCACCTTACACCCAATAGAAATTGCTCGACAAC
TCACTTTACTTGAATCAGATCTATACCGAGCTGTACAGCCATCAGAATTAGTTGGAAGTGTGTGGACAAA
AGAAGACAAAGAAATTAACTCTCCTAATCTTCTGAAAATGATTCGACATACCACCAACCTCACTCTGTGG
TTTGAGAAATGTATTGTAGAAACTGAAAATTTAGAAGAAAGAGTAGCTGTGGTGAGTCGAATTATTGAGA
TTCTACAAGTCTTTCAAGAGTTGAACAACTTTAATGGTGTCCTTGAGGTTGTCAGTGCTATGAATTCATC
ACCTGTTTACAGACTAGACCACACATTTGAGCAAATACCAAGTCGCCAGAAGAAAATTTTAGAAGAAGCT
CATGAATTGAGTGAAGATCACTATAAGAAATATTTGGCAAAACTCAGGTCTATTAATCCACCATGTGTGC
CTTTCTTTGGAATTTATCTCACTAATATCTTGAAAACAGAAGAAGGCAACCCTGAGGTCCTAAAAAGACA
TGGAAAAGAGCTTATAAACTTTAGCAAAAGGAGGAAAGTAGCAGAAATAACAGGAGAGATCCAGCAGTAC
CAAAATCAGCCTTACTGTTTACGAGTAGAATCAGATATCAAAAGGTTCTTTGAAAACTTGAATCCGATGG
GAAATAGCATGGAGAAGGAATTTACAGATTATCTTTTCAACAAATCCCTAGAAATAGAACCACGAAACCC
TAAGCCTCTCCCAAGATTTCCAAAAAAATATAGCTATCCCCTAAAATCTCCTGGTGTTCGTCCATCAAAC
CCAAGACCAGGTACCTAA
```

**Annotation**

PB1: PB1 domain from human p62 (residues 1-102)

AG: green fluorescent protein Azami-Green

FRB: FRB domain from human mammalian target of rapamycin (mTOR) (residues 2021-2113 with a T2098L mutation)

$TEV_{cs}$: TEV protease cleavage sequence (ENLYFQ/L)

mTagBFP2: blue fluorescent protein mTagBFP2

HA: HA epitope tag

$SOS1_{cat}$: Rem-Cdc25 (RasGEF) domain from human SOS1 (residues 566-1049)

**w**

**pCMV-PB1-AG-FRB-TEVuncs-BFP-HA-SOS**

>Amino acid sequence
MASLTVKAYLLGKEDAAREIRRFSFCCSPEPEAEAEAAAGPGPCERLLSRVAALFPALRPGGFQAHYRDE
DGDLVAFSSDEELTMAMSYVKDDIFRIYIKEKTGSGSGGSGAGGSAGSGAGGSAGSGAGGSAGSGAGGSA
GSGAGGSASGSSPGSGSGGSGAGGSAGSGAGGSAGSGAGGSAGSGAGGSAGSGAGGSASGSSPGNSADGG
GGSGGSGGSGGGSTQGGSMVSVIKPEMKIKLCMRGTVNGHNFVIEGEGKGNPYEGTQILDLNVTEGAPLP
FAYDILTTVFQYGNRAFTKYPADIQDYFKQTFPEGYHWERSMTYEDQGICTATSNISMRGDCFFYDIRFD
GVNFPPNGPVMQKKTLKWEPSTEKMYVRDGVLKGDVNMALLLEGGGHYRCDFKTTYKAKKDVRLPDYHFV
DHRIEILKHDKDYNKVKLYENAVARYSMLPSQAKGTGTAAENSGNSRTKLMILWHEMWHEGLEEASRLYF
GERNVKGMFEVLEPLHAMMERGPQTLKETSFNQAYGRDLMEAQEWCRKYMKSGNVKDLLQAWDLYYHVFR
RISKPGGSGAGSGAGSGAGNEYLQFLVDGGSAGGSGGVDMVSKGEELIKENMHMKLYMEGTVDNHHFKCT
SEGEGKPYEGTQTMRIKVVEGGPLPFAFDILATSFLYGSKTFINHTQGIPDFFKQSFPEGFTWERVTTYE
DGGVLTATQDTSLQDGCLIYNVKIRGVNFTSNGPVMQKKTLGWEAFTETLYPADGGLEGRNDMALKLVGG
SHLIANAKTTYRSKKPAKNLKMPGVYYVDYRLERIKEANNETYVEQHEVAVARYCDLPSKLGHKLNRSGS
YPYDVPDYARSQMRLPSADVYRFAEPDSEENIIFEENMQPKAGIPIIKAGTVIKLIERLTYHMYADPNFV
RTFLTTYRSFCKPQELLSLIIERFEIPEPEPTEADRIAIENGDQPLSAELKRFRKEYIQPVQLRVLNVCR
HWVEHHFYDFERDAYLLQRMEEFIGTVRGKAMKKWVESITKIIQRKKIARDNGPGHNITFQSSPPTVEWH
ISRPGHIETFDLLTLHPIEIARQLTLLESDLYRAVQPSELVGSVWTKEDKEINSPNLLKMIRHTTNLTLW
FEKCIVETENLEERVAVVSRIIEILQVFQELNNFNGVLEVVSAMNSSPVYRLDHTFEQIPSRQKKILEEA
HELSEDHYKKYLAKLRSINPPCVPFFGIYLTNILKTEEGNPEVLKRHGKELINFSKRRKVAEITGEIQQY
QNQPYCLRVESDIKRFFENLNPMGNSMEKEFTDYLFNKSLEIEPRNPKPLPRFPKKYSYPLKSPGVRPSN
PRPGT*

>DNA sequence
ATGGCGTCGCTCACCGTGAAGGCCTACCTTCTGGGCAAGGAGGACGCGGCGCGCGAGATTCGCCGCTTCA
GCTTCTGTTGCAGCCCCGAGCCTGAGGCGGAAGCCGAGGCTGCGGCGGGTCCGGGACCCTGCGAGCGGCT
GCTGAGCCGGGTGGCCGCCCTGTTCCCCGCGCTGCGGCCTGGCGGCTTCCAGGCGCACTACCGCGATGAG
GACGGGGACTTGGTTGCCTTTTCCAGTGACGAGGAATTGACAATGGCCATGTCCTACGTGAAGGATGACA
TCTTCCGAATCTACATTAAAGAGAAAACCGGTTCTGGGAGTGGCGGTAGTGGTGCTGGAGGCAGCGCAGG
CTCTGGAGCAGGCGGCAGTGCCGGAAGTGGCGCTGGAGGGTCTGCCGGATCTGGAGCCGGTGGTTCTGCC
GGCTCCGGCGCTGGTGGGAGCGCTTCCGGAAGTAGTCCCGGTTCTGGGAGTGGCGGTAGTGGTGCTGGAG
GCAGCGCAGGCTCTGGAGCAGGCGGCAGTGCCGGAAGTGGCGCTGGAGGGTCTGCCGGATCTGGAGCCGG
TGGTTCTGCCGGCTCCGGCGCTGGTGGGAGCGCTTCCGGAAGTAGTCCCGGTAATTCCGCTGACGGCGGC
GGAGGATCGGGTGGTAGTGGTGGTTCAGGAGGAGGATCGACCCAAGGAGGATCCATGGTGAGCGTGATCA
AGCCCGAGATGAAGATCAAGCTGTGCATGAGGGGCACCGTGAACGGCCACAACTTCGTGATCGAGGGCGA
GGGCAAGGGCAACCCCTACGAGGGCACCCAGATCCTGGACCTGAACGTGACCGAGGGCGCCCCCCTGCCC
TTCGCCTACGACATCCTGACCACCGTGTTCCAGTACGGCAACAGGGCCTTCACCAAGTACCCCGCCGACA
TCCAGGACTACTTCAAGCAGACCTTCCCCGAGGGCTACCACTGGGAGAGGAGCATGACCTACGAGGACCA
GGGCATCTGCACCGCCACCAGCAACATCAGCATGAGGGGCGACTGCTTCTTCTACGACATCAGGTTCGAC
GGCGTGAACTTCCCCCCCAACGGCCCCGTGATGCAGAAGAAGACCCTGAAGTGGGAGCCCAGCACCGAGA
AGATGTACGTGAGGGACGGCGTGCTGAAGGGCGACGTGAACATGGCCCTGCTGCTGGAGGGCGGCGGCCA
CTACAGGTGCGACTTCAAGACCACCTACAAGGCCAAGAAGGACGTGAGGCTGCCCGACTACCACTTCGTG
GACCACAGGATCGAGATCCTGAAGCACGACAAGGACTACAACAAGGTGAAGCTGTACGAGAACGCCGTGG
CCAGGTACAGCATGCTGCCCAGCCAGGCCAAGGGTACCGGAACTGCAGCAGAGAATTCGGGAAACTCGAG
AACAAAGCTTATGATCCTCTGGCATGAGATGTGGCATGAAGGCCTGGAAGAGGCATCTCGTTTGTACTTT
GGGGAAAGGAACGTGAAAGGCATGTTTGAGGTGCTGGAGCCCTTGCATGCTATGATGGAACGGGGCCCCC
AGACTCTGAAGGAAACATCCTTTAATCAGGCCTATGGTCGAGATTTAATGGAGGCCCAAGAGTGGTGCAG
GAAGTACATGAAATCAGGGAATGTCAAGGACCTCCTCCAAGCCTGGGACCTCTATTATCATGTGTTCCGA
CGAATCTCAAAGCCCGGGGGTAGTGGTGCTGGCTCTGGTGCTGGTAGTGGCGCTGGTAACGAATATCTGC
AGTTTCTGGTCGATGGTGGTAGTGCTGGTGGTTCCGGAGGTGTCGACATGGTGTCTAAGGGCGAAGAGCT
GATTAAGGAGAACATGCACATGAAGCTGTACATGGAGGGCACCGTGGACAACCATCACTTCAAGTGCACA
TCCGAGGGCGAAGGCAAGCCCTACGAGGGCACCCAGACCATGAGAATCAAGGTGGTCGAGGGCGGCCCTC

TCCCCTTCGCCTTCGACATCCTGGCTACTAGCTTCCTCTACGGCAGCAAGACCTTCATCAACCACACCCA
GGGCATCCCCGACTTCTTCAAGCAGTCCTTCCCTGAGGGCTTCACATGGGAGAGAGTCACCACATACGAA
GACGGGGGCGTGCTGACCGCTACCCAGGACACCAGCCTCCAGGACGGCTGCCTCATCTACAACGTCAAGA
TCAGAGGGGTGAACTTCACATCCAACGGCCCTGTGATGCAGAAGAAAACACTCGGCTGGGAGGCCTTCAC
CGAGACGCTGTACCCCGCTGACGGCGGCCTGGAAGGCAGAAACGACATGGCCCTGAAGCTCGTGGGCGGG
AGCCATCTGATCGCAAACGCCAAGACCACATATAGATCCAAGAAACCCGCTAAGAACCTCAAGATGCCTG
GCGTCTACTATGTGGACTACAGACTGGAAAGAATCAAGGAGGCCAACAACGAGACCTACGTCGAGCAGCA
CGAGGTGGCAGTGGCCAGATACTGCGACCTCCCTAGCAAACTGGGGCACAAGCTTAATAGATCCGGCTCT
TACCCATACGATGTTCCAGATTACGCTAGATCTCAGATGAGGCTGCCTAGTGCTGATGTTTATAGATTTG
CAGAGCCTGACTCTGAAGAGAATATTATATTTGAAGAGAACATGCAGCCCAAGGCTGGAATTCCAATTAT
CAAAGCAGGAACTGTTATTAAACTTATAGAGAGGCTTACGTACCATATGTACGCAGATCCCAATTTTGTT
CGGACATTTCTTACAACATACAGATCCTTTTGCAAACCTCAAGAACTACTGAGTCTTATAATAGAAAGGT
TTGAAATTCCAGAGCCTGAGCCAACAGAAGCTGATCGCATAGCTATAGAGAATGGAGATCAACCCTTGAG
TGCAGAACTGAAAAGATTTAGAAAAGAATATATACAGCCTGTGCAACTGCGAGTATTAAATGTATGTCGG
CACTGGGTAGAGCACCACTTCTATGATTTTGAAAGAGATGCATATCTTTTGCAACGAATGGAAGAATTTA
TTGGAACAGTAAGAGGTAAAGCAATGAAAAAATGGGTTGAATCCATCACTAAAATAATCCAAAGGAAAAA
AATTGCAAGAGACAATGGACCAGGTCATAATATTACATTTCAGAGTTCACCTCCCACAGTTGAGTGGCAT
ATAAGCAGACCTGGGCACATAGAGACTTTTGACCTGCTCACCTTACACCCAATAGAAATTGCTCGACAAC
TCACTTTACTTGAATCAGATCTATACCGAGCTGTACAGCCATCAGAATTAGTTGGAAGTGTGTGGACAAA
AGAAGACAAAGAAATTAACTCTCCTAATCTTCTGAAAATGATTCGACATACCACCAACCTCACTCTGTGG
TTTGAGAAATGTATTGTAGAAACTGAAAATTTAGAAGAAAGAGTAGCTGTGGTGAGTCGAATTATTGAGA
TTCTACAAGTCTTTCAAGAGTTGAACAACTTTAATGGTGTCCTTGAGGTTGTCAGTGCTATGAATTCATC
ACCTGTTTACAGACTAGACCACACATTTGAGCAAATACCAAGTCGCCAGAAGAAAATTTTAGAAGAAGCT
CATGAATTGAGTGAAGATCACTATAAGAAATATTTGGCAAAACTCAGGTCTATTAATCCACCATGTGTGC
CTTTCTTTGGAATTTATCTCACTAATATCTTGAAAACAGAAGAAGGCAACCCTGAGGTCCTAAAAAGACA
TGGAAAAGAGCTTATAAACTTTAGCAAAAGGAGGAAAGTAGCAGAAATAACAGGAGAGATCCAGCAGTAC
CAAAATCAGCCTTACTGTTTACGAGTAGAATCAGATATCAAAAGGTTCTTTGAAAACTTGAATCCGATGG
GAAATAGCATGGAGAAGGAATTTACAGATTATCTTTTCAACAAATCCCTAGAAATAGAACCACGAAACCC
TAAGCCTCTCCCAAGATTTCCAAAAAAATATAGCTATCCCCTAAAATCTCCTGGTGTTCGTCCATCAAAC
CCAAGACCAGGTACCTAA

**Annotation**

PB1: PB1 domain from human p62 (residues 1-102)

AG: green fluorescent protein Azami-Green

FRB: FRB domain from human mammalian target of rapamycin (mTOR) (residues 2021-2113 with a T2098L mutation)

$TEV_{uncs}$: TEV protease uncleavable sequence (NEYLQFL)

mTagBFP2: blue fluorescent protein mTagBFP2

HA: HA epitope tag

$SOS1_{cat}$: Rem-Cdc25 (RasGEF) domain from human SOS1 (residues 566-1049)

**x**

**pCMV-tdiRFP670-FKBP**

```
>Amino acid sequence
MARKVDLTSCDREPIHIPGSIQPCGCLLACDAQAVRITRITENAGAFFGRETPRVGELLADYFGETEAHA
LRNALAQSSDPKRPALIFGWRDGLTGRTFDISLHRHDGTSIIEFEPAAAEQADNPLRLTRQIIARTKELK
SLEEMAARVPRYLQAMLGYHRVMLYRFADDGSGMVIGEAKRSDLESFLGQHFPASLVPQQARLLYLKNAI
RVVSDSRGISSRIVPEHDASGAALDLSFAHLRSISPCHLEFLRNMGVSASMSLSIIIDGTLWGLIICHHY
EPRAVPMAQRVAAEMFADFLSLHFTAAHHQRGHATGSTGSGSAEGGTASSEDNMARKVDLTSCDREPIHI
PGSIQPCGCLLACDAQAVRITRITENAGAFFGRETPRVGELLADYFGETEAHALRNALAQSSDPKRPALI
FGWRDGLTGRTFDISLHRHDGTSIIEFEPAAAEQADNPLRLTRQIIARTKELKSLEEMAARVPRYLQAML
GYHRVMLYRFADDGSGMVIGEAKRSDLESFLGQHFPASLVPQQARLLYLKNAIRVVSDSRGISSRIVPEH
DASGAALDLSFAHLRSISPCHLEFLRNMGVSASMSLSIIIDGTLWGLIICHHYEPRAVPMAQRVAAEMFA
DFLSLHFTAAHHQRMYKSGLRSRAMGVQVETISPGDGRTFPKRGQTCVVHYTGMLEDGKKFDSSRDRNKP
FKFMLGKQEVIRGWEEGVAQMSVGQRAKLTISPDYAYGATGHPGIIPPHATLVFDVELLKLEEFCSRRYR
GPGIHRI*

>DNA sequence
ATGGCTCGCAAGGTGGACCTGACCAGCTGCGACCGCGAGCCCATCCACATCCCCGGCAGCATCCAGCCCT
GCGGCTGCCTGCTGGCCTGCGACGCCCAGGCCGTGCGCATCACCCGCATCACCGAGAACGCCGGCGCCTT
CTTCGGCCGCGAGACCCCCCGCGTGGGCGAGCTGCTGGCCGACTACTTCGGCGAGACCGAGGCCCACGCC
CTGCGCAACGCCCTGGCCCAGAGCAGCGACCCCAAGCGCCCCGCCCTGATCTTCGGCTGGCGCGACGGCC
TGACCGGCCGCACCTTCGACATCAGCCTGCACCGCCACGACGGCACCAGCATCATCGAGTTCGAGCCCGC
CGCCGCCGAGCAGGCCGACAACCCCCTGCGCCTGACCCGCCAGATCATCGCCCGCACCAAGGAGCTGAAG
AGCCTGGAGGAGATGGCCGCCCGCGTGCCCCGCTACCTGCAGGCCATGCTGGGCTACCACCGCGTGATGC
TGTACCGCTTCGCCGACGACGGCAGCGGCATGGTGATCGGCGAGGCCAAGCGCAGCGACCTGGAGAGCTT
CCTGGGCCAGCACTTCCCCGCCAGCCTGGTGCCCCAGCAGGCCCGCCTGCTGTACCTGAAGAACGCCATC
CGCGTGGTGAGCGACAGCCGCGGCATCAGCAGCCGCATCGTGCCCGAGCACGACGCCAGCGGCGCCGCCC
TGGACCTGAGCTTCGCCCACCTGCGCAGCATCAGCCCCTGCCACCTGGAGTTCCTGCGCAACATGGGCGT
GAGCGCCAGCATGAGCCTGAGCATCATCATCGACGGCACCCTGTGGGGCCTGATCATCTGCCACCACTAC
GAGCCCCGCGCCGTGCCCATGGCCCAGCGCGTGGCCGCCGAGATGTTCGCCGACTTCCTGAGCCTGCACT
TCACCGCCGCTCACCATCAGAGAGGACATGCTACTGGAAGCACTGGAAGCGGCAGCGCTGAGGGAGGCAC
AGCTTCTAGCGAAGATAATATGGCTCGCAAGGTGGACCTGACCAGCTGCGACCGCGAGCCCATCCACATC
CCCGGCAGCATCCAGCCCTGCGGCTGCCTGCTGGCCTGCGACGCCCAGGCCGTGCGCATCACCCGCATCA
CCGAGAACGCCGGCGCCTTCTTCGGCCGCGAGACCCCCCGCGTGGGCGAGCTGCTGGCCGACTACTTCGG
CGAGACCGAGGCCCACGCCCTGCGCAACGCCCTGGCCCAGAGCAGCGACCCCAAGCGCCCCGCCCTGATC
TTCGGCTGGCGCGACGGCCTGACCGGCCGCACCTTCGACATCAGCCTGCACCGCCACGACGGCACCAGCA
TCATCGAGTTCGAGCCCGCCGCCGCCGAGCAGGCCGACAACCCCCTGCGCCTGACCCGCCAGATCATCGC
CCGCACCAAGGAGCTGAAGAGCCTGGAGGAGATGGCCGCCCGCGTGCCCCGCTACCTGCAGGCCATGCTG
GGCTACCACCGCGTGATGCTGTACCGCTTCGCCGACGACGGCAGCGGCATGGTGATCGGCGAGGCCAAGC
GCAGCGACCTGGAGAGCTTCCTGGGCCAGCACTTCCCCGCCAGCCTGGTGCCCCAGCAGGCCCGCCTGCT
GTACCTGAAGAACGCCATCCGCGTGGTGAGCGACAGCCGCGGCATCAGCAGCCGCATCGTGCCCGAGCAC
GACGCCAGCGGCGCCGCCCTGGACCTGAGCTTCGCCCACCTGCGCAGCATCAGCCCCTGCCACCTGGAGT
TCCTGCGCAACATGGGCGTGAGCGCCAGCATGAGCCTGAGCATCATCATCGACGGCACCCTGTGGGGCCT
GATCATCTGCCACCACTACGAGCCCCGCGCCGTGCCCATGGCCCAGCGCGTGGCCGCCGAGATGTTCGCC
GACTTCCTGAGCCTGCACTTCACCGCCGCTCACCATCAGAGAATGTACAAGTCCGGACTCAGATCTCGAG
CTATGGGAGTGCAGGTGGAAACCATCTCCCCAGGAGACGGGCGCACCTTCCCCAAGCGCGGCCAGACCTG
CGTGGTGCACTACACCGGGATGCTTGAAGATGGAAAGAAATTTGATTCCTCCCGGGACAGAAACAAGCCC
TTTAAGTTTATGCTAGGCAAGCAGGAGGTGATCCGAGGCTGGGAAGAAGGGGTTGCCCAGATGAGTGTGG
GTCAGAGAGCCAAACTGACTATATCTCCAGATTATGCCTATGGTGCCACTGGGCACCCAGGCATCATCCC
ACCACATGCCACTCTCGTCTTCGATGTGGAGCTTCTAAAACTGGAAGAATTCTGCAGTCGACGGTACCGC
GGGCCCGGGATCCACCGGATCTAG
```

**Annotation**

iRFP670: near-infrared fluorescent protein iRFP670
FKBP: FKBP12

**y**

**pCAGGS-PB1-MR-LOV2**

>Amino acid sequence
MASLTVKAYLLGKEDAAREIRRFSFCCSPEPEAEAEAAAGPGPCERLLSRVAALFPALRPGGFQAHYRDE
DGDLVAFSSDEELTMAMSYVKDDIFRIYIKEKTGSGSGGSGAGGSAGSGAGGSAGSGAGGSAGSGAGGSA
GSGAGGSASGSSPGSGSGGSGAGGSAGSGAGGSAGSGAGGSAGSGAGGSAGSGAGGSASGSSPGNSADGG
GGSGGSGGSGGGSTQGGSMVGVIAKQMTYKVYMSGTVNGHYFEVEGDGKGKPYEGEQTVRLTVTKGGPLP
FAWDILSPQLMYGSITFTKYPEDIPDYFKQSFPEGFTWERIMNFEDGAVCTVSNDSSIQGNCFIYNVKIS
GVNFPPNGPVMQKKTQGWEPSTERLFARDGMLIGYNYMALKLEGGGHYLCEFKTTYKAKKPVRMPGFHFI
DRKLDVTSHNRDYTSVEQCEIAIARHSLLGGTGTAAENSGNSRTKLATTLERIEKNFVITDPRLPDNPII
FASDSFLQLTEYSREEILGRNCRFLQGPETDRATVRKIRDAIDNQTEVTVQLINYTKSGKKFWNLFHLQP
MRDQKGDVQYFIGVQLDGTEHVRDAAEREGVMLIKKTAENIDEAAKEL*

>DNA sequence
ATGGCGTCGCTCACCGTGAAGGCCTACCTTCTGGGCAAGGAGGACGCGGCGCGCGAGATTCGCCGCTTCA
GCTTCTGTTGCAGCCCCGAGCCTGAGGCGGAAGCCGAGGCTGCGGCGGGTCCGGGACCCTGCGAGCGGCT
GCTGAGCCGGGTGGCCGCCCTGTTCCCCGCGCTGCGGCCTGGCGGCTTCCAGGCGCACTACCGCGATGAG
GACGGGGACTTGGTTGCCTTTTCCAGTGACGAGGAATTGACAATGGCCATGTCCTACGTGAAGGATGACA
TCTTCCGAATCTACATTAAAGAGAAAACCGGTTCTGGGAGTGGCGGTAGTGGTGCTGGAGGCAGCGCAGG
CTCTGGAGCAGGCGGCAGTGCCGGAAGTGGCGCTGGAGGGTCTGCCGGATCTGGAGCCGGTGGTTCTGCC
GGCTCCGGCGCTGGTGGGAGCGCTTCCGGAAGTAGTCCCGGTTCTGGGAGTGGCGGTAGTGGTGCTGGAG
GCAGCGCAGGCTCTGGAGCAGGCGGCAGTGCCGGAAGTGGCGCTGGAGGGTCTGCCGGATCTGGAGCCGG
TGGTTCTGCCGGCTCCGGCGCTGGTGGGAGCGCTTCCGGAAGTAGTCCCGGTAATTCCGCTGACGGCGGC
GGAGGATCGGGTGGTAGTGGTGGTTCAGGAGGAGGATCGACCCAAGGAGGATCCATGGTGGGCGTGATCG
CCAAGCAGATGACCTACAAGGTGTACATGAGCGGCACCGTGAACGGCCACTACTTCGAGGTGGAGGGCGA
CGGCAAGGGCAAGCCCTACGAGGGCGAGCAGACCGTGAGGCTGACCGTGACCAAGGGTGGCCCCCTGCCC
TTCGCCTGGGACATCCTGAGCCCCCAGCTCATGTACGGCAGCATCACCTTCACCAAGTACCCCGAGGACA
TCCCCGACTACTTCAAACAGAGCTTCCCCGAGGGCTTCACCTGGGAGCGCATCATGAACTTCGAGGACGG
CGCCGTGTGCACCGTGAGCAACGACAGCAGCATCCAGGGCAACTGCTTCATCTACAACGTGAAGATCAGC
GGCGTGAACTTCCCCCCCAACGGCCCCGTGATGCAGAAGAAGACCCAGGGCTGGGAGCCCAGCACCGAGC
GCCTGTTCGCCCGCGACGGAATGCTGATCGGCTACAACTACATGGCCCTGAAGCTGGAGGGCGGCGGCCA
CTACCTGTGCGAGTTCAAGACCACCTACAAGGCCAAGAAGCCCGTGAGGATGCCCGGCTTCCACTTCATC
GACCGCAAGCTGGACGTGACCAGCCACAACCGCGACTACACCAGCGTGGAGCAGTGCGAGATCGCCATCG
CCCGCCACAGCCTGCTGGGCGGTACCGGAACTGCAGCAGAGAATTCGGGAAACTCGAGAACAAAGCTTGC
TACTACACTTGAACGTATTGAGAAGAACTTTGTCATTACTGACCCAAGATTGCCAGATAATCCCATTATA
TTCGCGTCCGATAGTTTCTTGCAGTTGACAGAATATAGCCGTGAAGAAATTTTGGGAAGAAACTGCAGGT
TTCTACAAGGTCCTGAAACTGATCGCGCGACAGTGAGAAAAATTAGAGATGCCATAGATAACCAAACAGA
GGTCACTGTTCAGCTGATTAATTATACAAAGAGTGGTAAAAAGTTCTGGAACCTCTTTCACTTGCAGCCT
ATGCGAGATCAGAAGGGAGATGTCCAGTACTTTATTGGGGTTCAGTTGGATGGAACTGAGCATGTCCGAG
ATGCTGCCGAGAGAGAGGGAGTCATGCTGATTAAGAAAACTGCAGAAAATATTGATGAGGCGGCAAAAGA
ACTTTAA

**Annotation**
PB1: PB1 domain from human p62 (residues 1-102)
MR: red fluorescent protein Monti-Red
LOV2: photoreceptor LOV2 domain

**z**

**pCMV-miRFP670-Zdk1**

>Amino acid sequence
MVAGHASGSPAFGTASHSNCEHEEIHLAGSIQPHGALLVVSEHDHRVIQASANAAEFLNLGSVLGVPLAE
IDGDLLIKILPHLDPTAEGMPVAVRCRIGNPSTEYCGLMHRPPEGGLIIELERAGPSIDLSGTLAPALER
IRTAGSLRALCDDTVLLFQQCTGYDRVMVYRFDEQGHGLVFSECHVPGLESYFGNRYPSSTVPQMARQLY
VRQRVRVLVDVTYQPVPLEPRLSPLTGRDLDMSGCFLRSMSPCHLQFLKDMGVRATLAVSLVVGGKLWGL
VVCHHYLPRFIRFELRAICKRLAERIATRITALESLYKSGLRSRAMVDNKFNKEKTRAGAEIHSLPNLNV
EQKFAFIVSLFDDPSQSANLLAEAKKLNDAQAPKTS*

>DNA sequence
ATGGTAGCAGGTCATGCCTCTGGCAGCCCCGCATTCGGGACCGCCTCTCATTCGAATTGCGAACATGAAG
AGATCCACCTCGCCGGCTCGATCCAGCCGCATGGCGCGCTTCTGGTCGTCAGCGAACATGATCATCGCGT
CATCCAGGCCAGCGCCAACGCCGCGGAATTTCTGAATCTCGGAAGCGTACTCGGCGTTCCGCTCGCCGAG
ATCGACGGCGATCTGTTGATCAAGATCCTGCCGCATCTCGATCCCACCGCCGAAGGCATGCCGGTCGCGG
TGCGCTGCCGGATCGGCAATCCCTCTACGGAGTACTGCGGTCTGATGCATCGGCCTCCGGAAGGCGGGCT
GATCATCGAACTCGAACGTGCCGGCCCGTCGATCGATCTGTCAGGCACGCTGGCGCCGGCGCTGGAGCGG
ATCCGCACGGCGGGTTCACTGCGCGCGCTGTGCGATGACACCGTGCTGCTGTTTCAGCAGTGCACCGGCT
ACGACCGGGTGATGGTGTATCGTTTCGATGAGCAAGGCCACGGCCTGGTATTCTCCGAGTGCCATGTGCC
TGGGCTCGAATCCTATTTCGGCAACCGCTATCCGTCGTCGACTGTCCCGCAGATGGCGCGGCAGCTGTAC
GTGCGGCAGCGCGTCCGCGTGCTGGTCGACGTCACCTATCAGCCGGTGCCGCTGGAGCCGCGGCTGTCGC
CGCTGACCGGGCGCGATCTCGACATGTCGGGCTGCTTCCTGCGCTCGATGTCGCCGTGCCATCTGCAGTT
CCTGAAGGACATGGGCGTGCGCGCCACCCTGGCGGTGTCGCTGGTGGTCGGCGGCAAGCTGTGGGGCCTG
GTTGTCTGTCACCATTATCTGCCGCGCTTCATCCGTTTCGAGCTGCGGGCGATCTGCAAACGGCTCGCCG
AAAGGATCGCGACGCGGATCACCGCGCTTGAGAGCCTGTACAAGTCCGGACTCAGATCTCGAGCTATGGT
GGATAACAAATTCAATAAAGAAAAGACGCGTGCCGGTGCGGAAATCCATTCTCTGCCAAACCTGAATGTT
GAGCAGAAGTTTGCCTTCATCGTGAGCCTGTTTGATGATCCATCTCAGAGCGCCAATCTGCTGGCCGAAG
CCAAAAAACTGAACGATGCCCAGGCCCCAAAAACTAGTTAA

**Annotation**
miRFP670: monomeric iRFP670
Zdk1: LOV2-binding protein Zdk1

**aa**

**pCAGGS-PB1-MontiRed-LOV2(superdark)**

>Amino acid sequence
MASLTVKAYLLGKEDAAREIRRFSFCCSPEPEAEAEAAAGPGPCERLLSRVAALFPALRPGGFQAHYRDE
DGDLVAFSSDEELTMAMSYVKDDIFRIYIKEKTGSGSGGSGAGGSAGSGAGGSAGSGAGGSAGSGAGGSA
GSGAGGSASGSSPGSGSGGSGAGGSAGSGAGGSAGSGAGGSAGSGAGGSAGSGAGGSASGSSPGNSADGG
GGSGGSGGSGGGSTQGGSMVGVIAKQMTYKVYMSGTVNGHYFEVEGDGKGKPYEGEQTVRLTVTKGGPLP
FAWDILSPQLMYGSITFTKYPEDIPDYFKQSFPEGFTWERIMNFEDGAVCTVSNDSSIQGNCFIYNVKIS
GVNFPPNGPVMQKKTQGWEPSTERLFARDGMLIGYNYMALKLEGGGHYLCEFKTTYKAKKPVRMPGFHFI
DRKLDVTSHNRDYTSVEQCEIAIARHSLLGGTGTAAENSGNSRTKLATTLERIEKNFVITDPRLPDNPII
FASDSFLQLTEYSREEILGRNARFLQGPETDRATVRKIRDAIDNQTEVTVQLINYTKSGKKFWNLFHLQP
MRDQKGDVQYFIGVQKDGTEHVRDAAEREAVMEIKKTAEEIDEAAKEL*

>DNA sequence
ATGGCGTCGCTCACCGTGAAGGCCTACCTTCTGGGCAAGGAGGACGCGGCGCGCGAGATTCGCCGCTTCA
GCTTCTGTTGCAGCCCCGAGCCTGAGGCGGAAGCCGAGGCTGCGGCGGGTCCGGGACCCTGCGAGCGGCT
GCTGAGCCGGGTGGCCGCCCTGTTCCCCGCGCTGCGGCCTGGCGGCTTCCAGGCGCACTACCGCGATGAG
GACGGGGACTTGGTTGCCTTTTCCAGTGACGAGGAATTGACAATGGCCATGTCCTACGTGAAGGATGACA
TCTTCCGAATCTACATTAAAGAGAAAACCGGTTCTGGGAGTGGCGGTAGTGGTGCTGGAGGCAGCGCAGG
CTCTGGAGCAGGCGGCAGTGCCGGAAGTGGCGCTGGAGGGTCTGCCGGATCTGGAGCCGGTGGTTCTGCC
GGCTCCGGCGCTGGTGGGAGCGCTTCCGGAAGTAGTCCCGGTTCTGGGAGTGGCGGTAGTGGTGCTGGAG
GCAGCGCAGGCTCTGGAGCAGGCGGCAGTGCCGGAAGTGGCGCTGGAGGGTCTGCCGGATCTGGAGCCGG
TGGTTCTGCCGGCTCCGGCGCTGGTGGGAGCGCTTCCGGAAGTAGTCCCGGTAATTCCGCTGACGGCGGC
GGAGGATCGGGTGGTAGTGGTGGTTCAGGAGGAGGATCGACCCAAGGAGGATCCATGGTGGGCGTGATCG
CCAAGCAGATGACCTACAAGGTGTACATGAGCGGCACCGTGAACGGCCACTACTTCGAGGTGGAGGGCGA
CGGCAAGGGCAAGCCCTACGAGGGCGAGCAGACCGTGAGGCTGACCGTGACCAAGGGTGGCCCCCTGCCC
TTCGCCTGGGACATCCTGAGCCCCCAGCTCATGTACGGCAGCATCACCTTCACCAAGTACCCCGAGGACA
TCCCCGACTACTTCAAACAGAGCTTCCCCGAGGGCTTCACCTGGGAGCGCATCATGAACTTCGAGGACGG
CGCCGTGTGCACCGTGAGCAACGACAGCAGCATCCAGGGCAACTGCTTCATCTACAACGTGAAGATCAGC
GGCGTGAACTTCCCCCCCAACGGCCCCGTGATGCAGAAGAAGACCCAGGGCTGGGAGCCCAGCACCGAGC
GCCTGTTCGCCCGCGACGGAATGCTGATCGGCTACAACTACATGGCCCTGAAGCTGGAGGGCGGCGGCCA
CTACCTGTGCGAGTTCAAGACCACCTACAAGGCCAAGAAGCCCGTGAGGATGCCCGGCTTCCACTTCATC
GACCGCAAGCTGGACGTGACCAGCCACAACCGCGACTACACCAGCGTGGAGCAGTGCGAGATCGCCATCG
CCCGCCACAGCCTGCTGGGCGGTACCGGAACTGCAGCAGAGAATTCGGGAAACTCGAGAACAAAGCTTGC
TACTACACTTGAACGTATTGAGAAGAACTTTGTCATTACTGACCCAAGATTGCCAGATAATCCCATTATA
TTCGCGTCCGATAGTTTCTTGCAGTTGACAGAATATAGCCGTGAAGAAATTTTGGGAAGAAACGCCAGGT
TTCTACAAGGTCCTGAAACTGATCGCGCGACAGTGAGAAAAATTAGAGATGCCATAGATAACCAAACAGA
GGTCACTGTTCAGCTGATTAATTATACAAAGAGTGGTAAAAAGTTCTGGAACCTCTTTCACTTGCAGCCT
ATGCGAGATCAGAAGGGAGATGTCCAGTACTTTATTGGGGTTCAGAAGGATGGAACTGAGCATGTCCGAG
ATGCTGCCGAGAGAGAGGCCGTCATGGAGATTAAGAAAACTGCAGAAGAGATTGATGAGGCGGCAAAAGA
ACTTTAA

**Annotation**
PB1: PB1 domain from human p62 (residues 1-102)
MR: red fluorescent protein Monti-Red
LOV2 (superdark): light insensitive LOV2 domain (with C450A/L514K/G528A/L531E/N538E mutations)

**ab**

**pCAG-PB1-AzamiGreen(Y63L)-LOV2**

>Amino acid sequence
MASLTVKAYLLGKEDAAREIRRFSFCCSPEPEAEAEAAAGPGPCERLLSRVAALFPALRPGGFQAHYRDE
DGDLVAFSSDEELTMAMSYVKDDIFRIYIKEKTGSGSGGSGAGGSAGSGAGGSAGSGAGGSAGSGAGGSA
GSGAGGSASGSSPGSGSGGSGAGGSAGSGAGGSAGSGAGGSAGSGAGGSAGSGAGGSASGSSPGNSADGG
GGSGGSGGSGGGSTQGGSMVSVIKPEMKIKLCMRGTVNGHNFVIEGEGKGNPYEGTQILDLNVTEGAPLP
FAYDILTTVFQLGNRAFTKYPADIQDYFKQTFPEGYHWERSMTYEDQGICTATSNISMRGDCFFYDIRFD
GVNFPPNGPVMQKKTLKWEPSTEKMYVRDGVLKGDVNMALLLEGGGHYRCDFKTTYKAKKDVRLPDYHFV
DHRIEILKHDKDYNKVKLYENAVARYSMLPSQAKGTGTAAENSGNSRTKLATTLERIEKNFVITDPRLPD
NPIIFASDSFLQLTEYSREEILGRNCRFLQGPETDRATVRKIRDAIDNQTEVTVQLINYTKSGKKFWNLF
HLQPMRDQKGDVQYFIGVQLDGTEHVRDAAEREGVMLIKKTAENIDEAAKEL*

>DNA sequence
ATGGCGTCGCTCACCGTGAAGGCCTACCTTCTGGGCAAGGAGGACGCGGCGCGCGAGATTCGCCGCTTCA
GCTTCTGTTGCAGCCCCGAGCCTGAGGCGGAAGCCGAGGCTGCGGCGGGTCCGGGACCCTGCGAGCGGCT
GCTGAGCCGGGTGGCCGCCCTGTTCCCCGCGCTGCGGCCTGGCGGCTTCCAGGCGCACTACCGCGATGAG
GACGGGGACTTGGTTGCCTTTTCCAGTGACGAGGAATTGACAATGGCCATGTCCTACGTGAAGGATGACA
TCTTCCGAATCTACATTAAAGAGAAAACCGGTTCTGGGAGTGGCGGTAGTGGTGCTGGAGGCAGCGCAGG
CTCTGGAGCAGGCGGCAGTGCCGGAAGTGGCGCTGGAGGGTCTGCCGGATCTGGAGCCGGTGGTTCTGCC
GGCTCCGGCGCTGGTGGGAGCGCTTCCGGAAGTAGTCCCGGTTCTGGGAGTGGCGGTAGTGGTGCTGGAG
GCAGCGCAGGCTCTGGAGCAGGCGGCAGTGCCGGAAGTGGCGCTGGAGGGTCTGCCGGATCTGGAGCCGG
TGGTTCTGCCGGCTCCGGCGCTGGTGGGAGCGCTTCCGGAAGTAGTCCCGGTAATTCCGCTGACGGCGGC
GGAGGATCGGGTGGTAGTGGTGGTTCAGGAGGAGGATCGACCCAAGGAGGATCCATGGTGAGCGTGATCA
AGCCCGAGATGAAGATCAAGCTGTGCATGAGGGGCACCGTGAACGGCCACAACTTCGTGATCGAGGGCGA
GGGCAAGGGCAACCCCTACGAGGGCACCCAGATCCTGGACCTGAACGTGACCGAGGGCGCCCCCCTGCCC
TTCGCCTACGACATCCTGACCACCGTGTTCCAGCTGGGCAACAGGGCCTTCACCAAGTACCCCGCCGACA
TCCAGGACTACTTCAAGCAGACCTTCCCCGAGGGCTACCACTGGGAGAGGAGCATGACCTACGAGGACCA
GGGCATCTGCACCGCCACCAGCAACATCAGCATGAGGGGCGACTGCTTCTTCTACGACATCAGGTTCGAC
GGCGTGAACTTCCCCCCCAACGGCCCCGTGATGCAGAAGAAGACCCTGAAGTGGGAGCCCAGCACCGAGA
AGATGTACGTGAGGGACGGCGTGCTGAAGGGCGACGTGAACATGGCCCTGCTGCTGGAGGGCGGCGGCCA
CTACAGGTGCGACTTCAAGACCACCTACAAGGCCAAGAAGGACGTGAGGCTGCCCGACTACCACTTCGTG
GACCACAGGATCGAGATCCTGAAGCACGACAAGGACTACAACAAGGTGAAGCTGTACGAGAACGCCGTGG
CCAGGTACAGCATGCTGCCCAGCCAGGCCAAGGGTACCGGAACTGCAGCAGAGAATTCGGGAAACTCGAG
AACAAAGCTTGCTACTACACTTGAACGTATTGAGAAGAACTTTGTCATTACTGACCCAAGATTGCCAGAT
AATCCCATTATATTCGCGTCCGATAGTTTCTTGCAGTTGACAGAATATAGCCGTGAAGAAATTTTGGGAA
GAAACTGCAGGTTTCTACAAGGTCCTGAAACTGATCGCGCGACAGTGAGAAAAATTAGAGATGCCATAGA
TAACCAAACAGAGGTCACTGTTCAGCTGATTAATTATACAAAGAGTGGTAAAAAGTTCTGGAACCTCTTT
CACTTGCAGCCTATGCGAGATCAGAAGGGAGATGTCCAGTACTTTATTGGGGTTCAGTTGGATGGAACTG
AGCATGTCCGAGATGCTGCCGAGAGAGAGGGAGTCATGCTGATTAAGAAAACTGCAGAAAATATTGATGA
GGCGGCAAAAGAACTTTAA

**Annotation**
PB1: PB1 domain from human p62 (residues 1-102)
AG(Y63L): non-fluorescent colorless Azami-Green mutant
LOV2: photoreceptor LOV2 domain

**ac**

**pCMV-mCherry-Zdk1-Vav2$_{cat}$**

>Amino acid sequence
MVSKGEEDNMAIIKEFMRFKVHMEGSVNGHEFEIEGEGEGRPYEGTQTAKLKVTKGGPLPFAWDILSPQF
MYGSKAYVKHPADIPDYLKLSFPEGFKWERVMNFEDGGVVTVTQDSSLQDGEFIYKVKLRGTNFPSDGPV
MQKKTMGWEASSERMYPEDGALKGEIKQRLKLKDGGHYDAEVKTTYKAKKPVQLPGAYNVNIKLDITSHN
EDYTIVEQYERAEGRHSTGGMDELYKSGLRSRAMVDNKFNKEKTRAGAEIHSLPNLNVEQKFAFIVSLFD
DPSQSANLLAEAKKLNDAQAPKTSRSRGPMKMGMTEDDKRSCCLLEIQETEAKYYRTLEDIEKNYMGPLR
LVLSPADMAAVFINLEDLIKVHHSFLRAIDVSMMAGGSTLAKVFLEFKERLLIYGEYCSHMEHAQSTLNQ
LLASREDFRQKVEECTLRVQDGKFKLQDLLVVPMQRVLKYHLLLKELLSHSADRPERQQLKEALEAMQDL
AMYINEVKRDKETLKKISEFQCSIENLQVKLEEFGRPKIDGELKVRSIVNHTKQDRYLFLFDKVVIVCKR
KGYSYELKEVIELLFHKMTDDPMHNKDIKKWSYGFYLIHLQGKQGFQFFCKTEDMKRKWMEQFEMAMSNI
KPDKANANHHSFQMYTFDKTTNCKACKMFLRGTFYQGYLCTRCGVGAHKECLEVIPPCK*

>DNA sequence
ATGGTGAGCAAGGGCGAGGAGGATAACATGGCCATCATCAAGGAGTTCATGCGCTTCAAGGTGCACATGG
AGGGCTCCGTGAACGGCCACGAGTTCGAGATCGAGGGCGAGGGCGAGGGCCGCCCCTACGAGGGCACCCA
GACCGCCAAGCTGAAGGTGACCAAGGGTGGCCCCCTGCCCTTCGCCTGGGACATCCTGTCCCCTCAGTTC
ATGTACGGCTCCAAGGCCTACGTGAAGCACCCCGCCGACATCCCCGACTACTTGAAGCTGTCCTTCCCCG
AGGGCTTCAAGTGGGAGCGCGTGATGAACTTCGAGGACGGCGGCGTGGTGACCGTGACCCAGGACTCCTC
CCTGCAGGACGGCGAGTTCATCTACAAGGTGAAGCTGCGCGGCACCAACTTCCCCTCCGACGGCCCCGTA
ATGCAGAAGAAGACCATGGGCTGGGAGGCCTCCTCCGAGCGGATGTACCCCGAGGACGGCGCCCTGAAGG
GCGAGATCAAGCAGAGGCTGAAGCTGAAGGACGGCGGCCACTACGACGCTGAGGTCAAGACCACCTACAA
GGCCAAGAAGCCCGTGCAGCTGCCCGGCGCCTACAACGTCAACATCAAGTTGGACATCACCTCCCACAAC
GAGGACTACACCATCGTGGAACAGTACGAACGCGCCGAGGGCCGCCACTCCACCGGCGGCATGGACGAGC
TGTACAAGTCCGGACTCAGATCTCGAGCTATGGTGGATAACAAATTCAATAAAGAAAAGACGCGTGCCGG
TGCGGAAATCCATTCTCTGCCAAACCTGAATGTTGAGCAGAAGTTTGCCTTCATCGTGAGCCTGTTTGAT
GATCCATCTCAGAGCGCCAATCTGCTGGCCGAAGCCAAAAAACTGAACGATGCCCAGGCCCCAAAAACTA
GTAGATCTCGAGGGCCCATGAAAATGGGCATGACTGAGGACGACAAGAGAAGCTGCTGCTTGTTAGAGAT
TCAGGAGACCGAGGCCAAGTACTACCGCACCCTGGAGGACATTGAGAAGAACTACATGGGTCCCTTGCGG
CTGGTGCTGAGCCCGGCGGATATGGCTGCTGTCTTCATCAACCTGGAGGACCTCATCAAGGTGCATCACA
GCTTTCTGCGAGCCATCGATGTGTCCATGATGGCTGGTGGCAGTACCCTGGCTAAGGTCTTTCTGGAGTT
TAAGGAAAGGCTCCTGATCTATGGAGAGTACTGTAGCCACATGGAACACGCTCAGAGTACACTGAACCAG
CTCCTCGCCAGCCGAGAGGACTTCAGGCAGAAAGTGGAGGAGTGCACACTCAGGGTTCAGGATGGCAAGT
TCAAGCTGCAAGACCTGCTGGTGGTGCCCATGCAACGGGTGCTGAAGTACCACCTGCTGCTCAAGGAGCT
CCTGAGCCATTCTGCAGACCGACCAGAAAGACAACAGCTCAAAGAAGCCCTGGAAGCCATGCAGGACTTG
GCCATGTACATCAATGAAGTGAAGCGGGACAAGGAGACCTTGAAGAAGATTAGCGAGTTCCAGTGCTCCA
TAGAAAACCTGCAAGTGAAGCTGGAGGAATTTGGGAGGCCAAAGATTGACGGGGAGCTTAAAGTCCGGTC
CATAGTCAACCACACCAAGCAAGACAGGTACCTGTTCCTATTTGACAAGGTGGTCATCGTGTGCAAGAGG
AAGGGCTACAGCTATGAGCTGAAGGAGGTCATTGAGCTGCTCTTCCACAAGATGACCGATGACCCGATGC
ACAACAAGGACATCAAGAAGTGGTCCTATGGCTTCTACCTGATTCACCTCCAAGGAAAGCAAGGCTTTCA
GTTCTTCTGCAAGACGGAAGACATGAAGCGGAAGTGGATGGAGCAGTTCGAGATGGCCATGTCAAACATC
AAGCCAGATAAGGCCAATGCCAACCATCATAGCTTCCAGATGTACACATTCGACAAGACTACCAACTGCA
AAGCCTGCAAGATGTTTCTCAGGGGTACCTTCTACCAGGGATACCTGTGTACCAGATGTGGCGTCGGGGC
ACACAAGGAATGCCTGGAGGTGATCCCCCCCTGCAAGTAA

**Annotation**
mCherry: red fluorescent protein mCherry
Zdk1: LOV2-binding protein Zdk1
Vav2$_{cat}$: DH-PH-CR domain from mouse Vav2 (residues 183-563)

**Supplementary Fig. 17 DNA and amino acid sequences of constructs used in this study.**

#### Supplementary Movies

**Supplementary Movie 1 Rapamycin-triggered FKBP-mCherry recruitment to $^{FRB}$PAC by the SPREC-In system** (time-lapse movie of **Fig. 1b**).

**Supplementary Movie 2 Acute sequestration of Vav2 activity by the SPREC-In system** (time-lapse movie of **Fig. 2a**).

**Supplementary Movie 3 Acute sequestration of SOS activity by the SPREC-In system** (time-lapse movie of **Fig. 2d**).

**Supplementary Movie 4 Rapamycin-triggered BFP-HA release by the SPREC-Out system** (time-lapse movie of **Fig. 3b**).

**Supplementary Movie 5 Acute release of Vav2 activity by the SPREC-Out system** (time-lapse movie of **Fig. 4a**).

**Supplementary Movie 6 Acute release of SOS activity by the SPREC-Out system** (time-lapse movie of **Fig. 4d**).

**Supplementary Movie 7 Light-mediated, reversible release and sequestration of miRFP-Zdk1 by the optoSPREC system** (time-lapse movie of **Fig. 5b**).

**Supplementary Movie 8 Light-mediated reversible control of Vav2 activity by the optoSPREC system** (time-lapse movie of **Fig. 5d**).

For all movies, scale bars are 20 μm.
